# Does the BioEmu2 Preview Improve on BioEmu-1 for Protein Conformational Ensembles?

**DOI:** 10.64898/2026.11.24.769054

**Authors:** Xavier Chee Wezen, Yu Zhu, Lee Sue Han, Adrian JT Teo, Taufiq Rahman

## Abstract

Generative deep-learning models such as BioEmu aim to sample the conformational ensembles of proteins, but it is not known whether BioEmu2 shows improvement over its predecessor BioEmu-1 in reaching experimentally solved states and reproducing measured populations. To test this, we compared BioEmu-1 with the preview release of BioEmu2 on 22 protein systems with 49 experimentally solved states. This is done together with ColabFold and all-atom molecular dynamics as reference methods. To this end, we found that BioEmu2 samples almost never fail a backbone geometry filter (median 0.010% per system, against 5.60% for BioEmu-1). However, BioEmu2 reached every state within 3.0 Å in 10 of the 22 systems, against 15 for BioEmu - 1 (exact McNemar test, *p* = 0.0625). The difference lay mainly in states that need a ligand, nucleic acid, partner protein or phosphorylation that the sampled sequence lacks. When the missing phosphorylation was supplied, BioEmu2 reached the active states of ERK2 and p38α. Additionally, adding DNA raised the percentage of filter-passing DNA polymerase β samples in the DNA-bound arrangement from 2.1% to 60.0%. The two models favoured different states in most systems but neither model reproduced the populations measured in solution. Only one of six comparisons fell within the experimental tolerance, and BioEmu2 tended to place almost all its samples in a single state. Taken together, these results suggest that BioEmu2 produces cleaner samples but does not reach more states, and that the sampling frequencies of either model should not be read as populations.

## 1. Introduction

Many proteins carry out their functions by moving between distinct protein conformational states.^1^ For example, the kinase domain of ABL1 switches between an active state ( adopted by 88% of the population) and two inactive states, namely I1 and I2 (each adopted by about 6%).^2^ These populations determine the function of the kinase, and mutations can shift these populations.^2^ Such minor states matter for drug design, because an inhibitor normally binds one state selectively. For example, the drug imatinib binds the minor I2 state of ABL1. ^2^ In the same way, some cryptic binding pockets would transiently open in certain conformational states only. Knowing the probability that such a pocket is open is important for structure-based drug discovery.^3^

However, these conformational states can interconvert slowly. This makes them difficult to sample with conventional molecular dynamics (MD) simulations. For example, the three states of ABL1 interconvert on a millisecond time scale. Their exchange rates of about 47 and 89 s ⁻¹ correspond to exchange times of about 21 and 11 ms, ^2^ which are more than 100,000 times longer than each of the three 100 ns MD replicas that the ATLAS database provides for each protein.^4^ Machine-learning methods offer another way to find these states. The structure-prediction model AlphaFold2^5^ normally predicts one structure from a multiple sequence alignment, which is a set of sequences of related proteins. It can predict other conformations when it is given only part of this alignment, either a small random subset of the sequences or a cluster of similar sequences. ^6,7^

Generative models of protein ensembles work differently from both MD and AlphaFold2. They are trained to sample a whole ensemble of structures directly, rather than to simulate motion over time or to return a few predictions. One such model is BioEmu-1 (published as BioEmu), which was trained to emulate the equilibrium ensembles of proteins.^8^ It generates the backbone atoms (N, Cα and C) and the Cβ atoms of a protein from its sequence. It was fine-tuned on more than 500,000 experimental protein stabilities,^8,9^ it predicted folding free energies with a mean absolute error below 0.9 kcal/mol, and it sampled 83% of a set of domain-motion reference structures within 3 Å.^8^ However, folding free energies describe the balance between folded and unfolded protein, so they do not show whether the model gives the folded states of a protein their correct relative populations.

Nonetheless, despite these advances, independent tests have found several limitations with BioEmu-1 ensembles. BioEmu-1 reproduced local flexibility well with correlations of root-mean-square fluctuations above 0.7 in 39 of 50 proteins from the ATLAS set. However, it did not separate cancer driver mutations from passenger mutations, and its samples were skewed towards ligand - bound states.^13^ Similarly, a comparison of machine-learning tools for ensemble prediction found only small or negligible shifts for mutations that are known to change conformational populations. ^14^ Two other studies found that such models often sampled mainly one state. In wild-type T4 lysozyme, BioEmu-1 and six other generative methods largely sampled only one of four states. ^15^ In four further proteins with several states, BioEmu-1 and structure-prediction models frequently defaulted to a dominant state represented in the Protein Data Bank (PDB).^16^ When BioEmu-1 structures of 26 kinases were used for ensemble docking in virtual screening, the screening performance did not improve on docking to crystal structures and was often worse. ^17^ Another study noted that it remains unclear how the states generated by BioEmu-1 are populated, and it estimated their populations by seeding short MD simulations with BioEmu-1 ensembles and building Markov state models.^18^

Recently, BioEmu2 (the successor of BioEmu-1) was released as a preview in October 2026.^10,11^ BioEmu2 is a diffusion model of about 500 million parameters that generates all atoms, and it also accepts nucleic acids, ligands and modified residues as input. ^10,11^ To evaluate BioEmu2, the developers of BioEmu2 compared the model with MD simulations of single proteins, protein dimers and protein–ligand complexes.^12^ They found that BioEmu2 sampled about half of the states that MD found in proteins and protein complexes. However, on single proteins, BioEmu-1 still performed best, likely because BioEmu-1 had been fine-tuned on experimental stabilities.^11,12^

With its all-atom output and wider range of inputs, BioEmu2 may address some of the known limitations of BioEmu-1. To our knowledge, there is currently no independent study that has compared the two models against experimental structures and measured populations. Such a comparison has to test several properties of an ensemble separately. Each sample should be a physically valid structure, the model should reach the experimentally solved states, and the populations of these states should match the measured values and change in the same direction as the measured populations when the protein is perturbed. Good performance on one property need not imply good performance on the next. This is because a model that reaches every observed state can still give these states the wrong populations. In addition, when a state was solved with a ligand, nucleic acid, partner protein or phosphorylation, a model given only the protein sequence may miss it without being in error. Therefore, in this work, we compared BioEmu-1 and BioEmu2 on 22 protein systems from 14 families against experimental structures and solution measurements. After checking every sample for broken or clashing backbones, we compared the two models at three levels, namely (a) the recovery of experimentally solved states, (b) the agreement with measured populations and (c) the response of these populations to mutations, phosphorylation and ligands (Figure 1a). We also tested whether supplying the missing partner or phosphorylation helped BioEmu2 to reach states that it had missed, and we scored ColabFold^19^ and all-atom MD in the same way as reference methods. Finally, we used ABL1 to test the response to perturbation, because it is the only system in our set with measured populations for three different experimentally resolved conformational states and for their changes on mutation and phosphorylation.^2^

**Figure 1.**
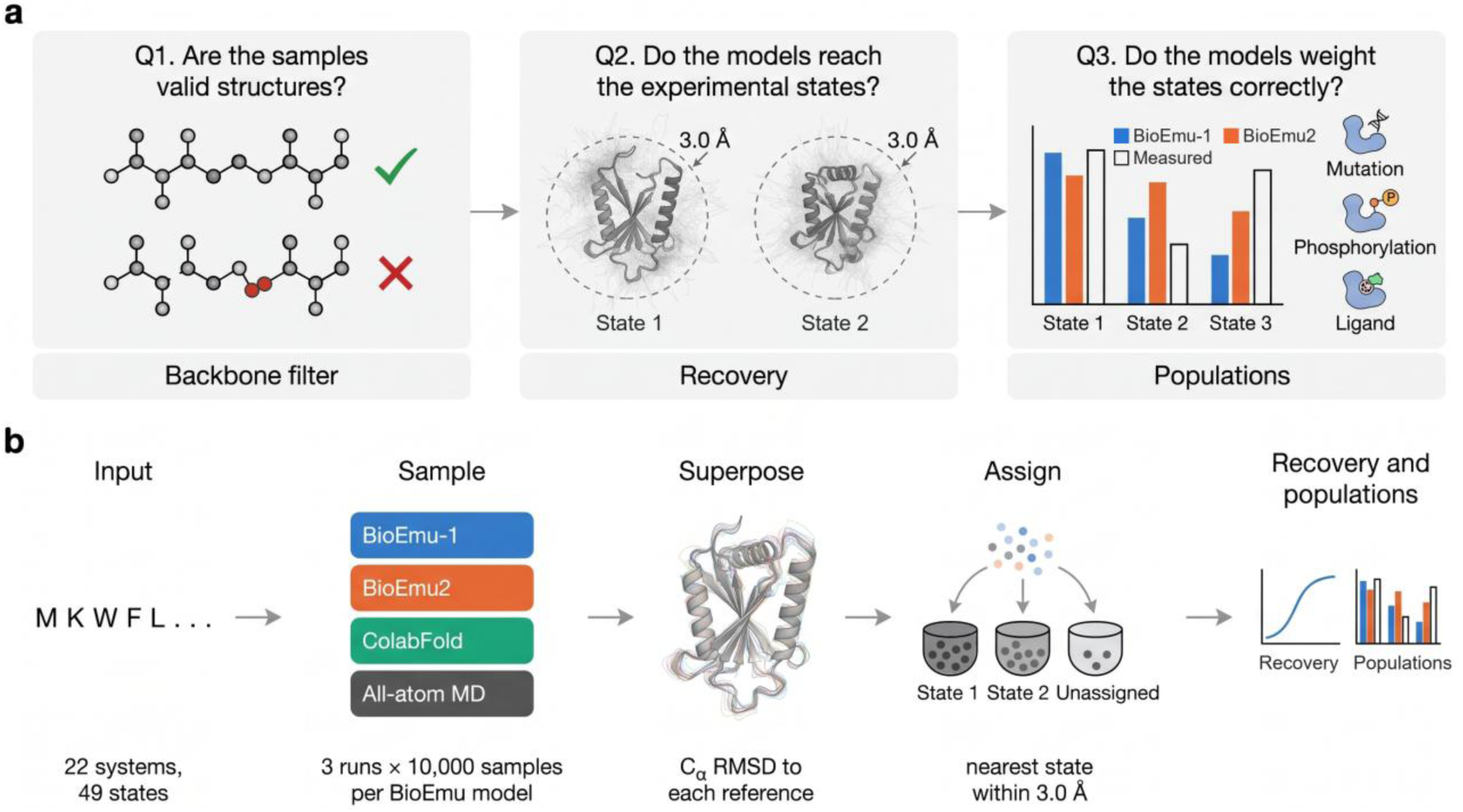
Overview of the study. (a) The three questions addressed in this work, namely whether the samples are physically valid structures (Q1), whether the models reach the experimentally solved states (Q2) and whether the models reproduce the measured populations of these states (Q3). (b) Study workflow. The sequence of each system was the input for sampling with BioEmu-1 and BioEmu2, with ColabFold and all-atom molecular dynamics (MD) run as reference methods. Each sample was superposed on and compared with every reference structure. Samples that passed the backbone filter were then assigned to their nearest state, or by collective variables or backbone rules in five systems. Finally, the samples were analysed for how readily they reached each state (recovery) and for whether the proportions of samples in each state matched the measured values, where available (populations).

## 2. Methods

### 2.1 Study Design

We compared two generative models from Microsoft Research, BioEmu-1^8^ and the preview release of BioEmu2.^10–12^ All BioEmu2 results in this paper refer to this preview release. ColabFold and all-atom MD were run as reference methods, and experimental structures defined the protein conformational states against which all methods were scored.

For clarity, we define three terms. A "model" is one of the two programs, BioEmu-1 or BioEmu2, and a "sample" is a single three-dimensional structure that a model generates. The structures predicted by ColabFold are called ColabFold models. A "state" is a protein conformation that has been solved experimentally.

### 2.2 Data Curation: Systems, Reference States and Condition Labels

The data set comprises 22 protein systems from 14 protein families (Table 1). These systems contain loop motions, side-chain switches, domain closures and rearrangements between domains. We chose systems from diverse families because performance in sampling benchmarks can depend on the nature of the protein.^20^ In total, 11 systems were chosen for this study (the study systems), and the other 11 were taken from a published benchmark (the benchmark systems).

**Table 1.**
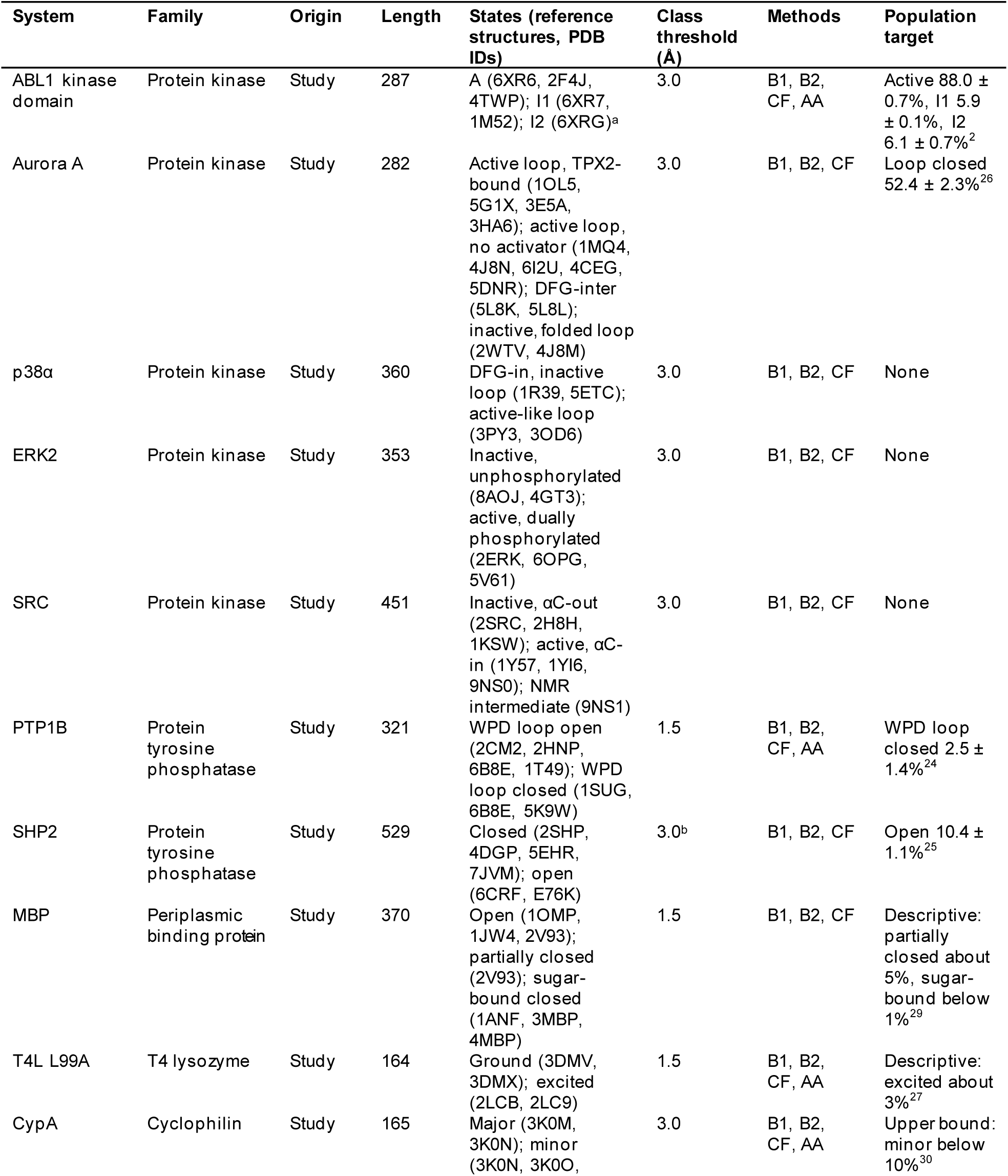
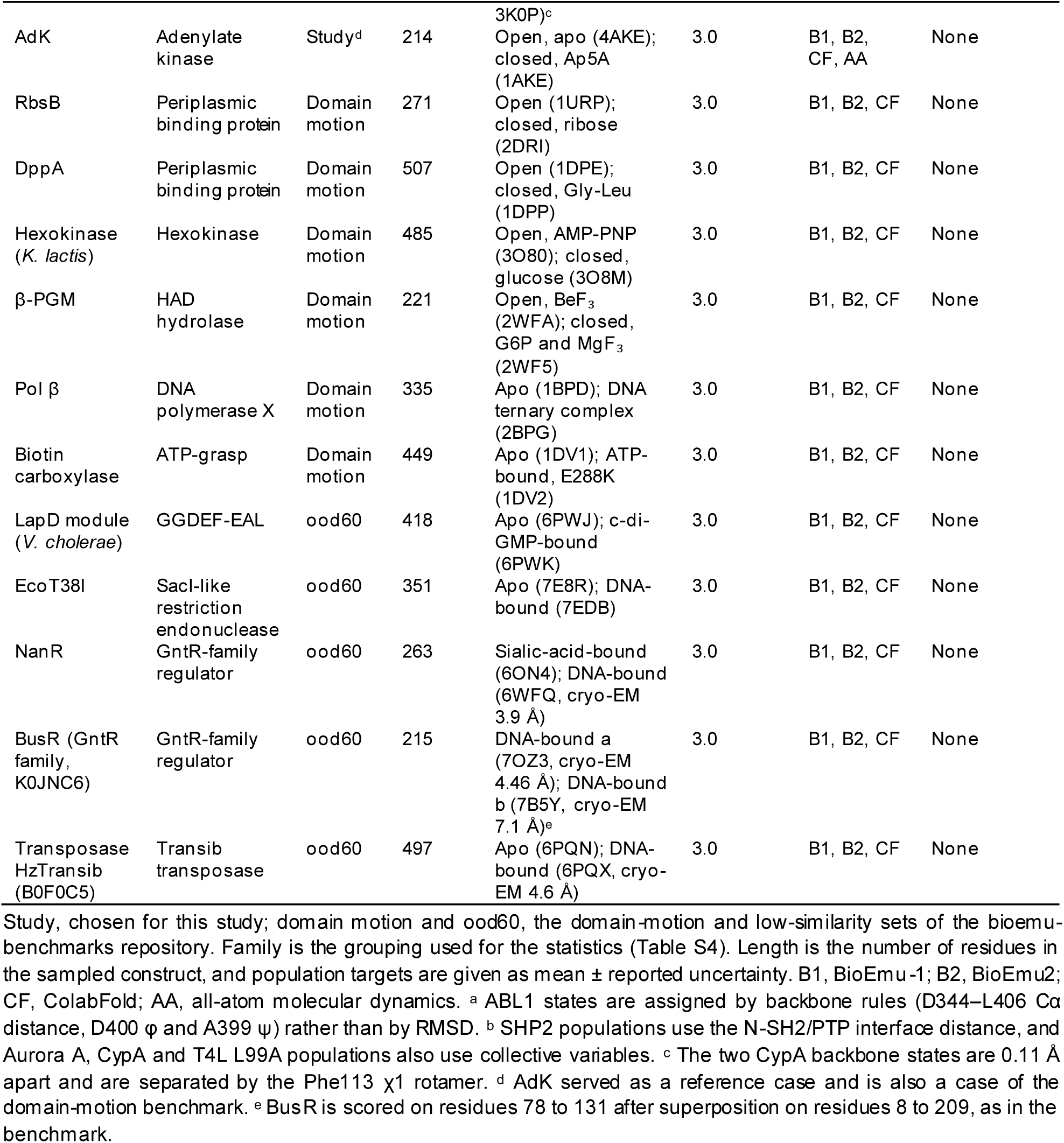
The 22 systems, their reference states and the methods run for each.

Seven study systems were chosen because the populations of their states have been measured in solution. These systems are ABL1, Aurora A, PTP1B, SHP2, maltose-binding protein (MBP), T4 lysozyme L99A (T4L L99A) and cyclophilin A (CypA). Three further kinases without a usable population target were added for state recovery only, namely p38α, ERK2 and SRC. SRC also has a rearrangement between domains and an intermediate state determined by NMR in solution.^21^ Adenylate kinase (AdK) was included as a reference case since it is the worked example in the BioEmu2 documentation.^10^

The benchmark systems were taken unchanged, with their published reference states, from the multi-conformation test cases of the bioemu-benchmarks repository.^22^ Six came from its domain-motion set and five came from a set selected for low sequence similarity to the BioEmu-1 training data (ood60). Two kinds of benchmark protein were left out, namely small fold-switching proteins that change fold only with a binding partner, and proteins longer than 549 residues. One ood60 system is the cytoplasmic module of LapD from *Vibrio cholerae*, which comprises its S helix and its GGDEF and EAL domains and is called the LapD module below.

The 22 systems have 49 states (with two to four states per system). Each state is defined by one to five experimental structures. BioEmu2 was trained partly on PDB structures in the copy prepared for Boltz-1.^11^ This training set contained entries released before 30 September 2021.^23^ Of the 84 reference structures, 79 were released before that date, so BioEmu2 may have seen many of them during training (Table S1). In contrast, BioEmu-1 was not trained on PDB entries, and 14 of the 22 proteins were among its published test proteins, which were removed from its training data. ^8^ Any advantage from overlap with the training data would therefore favour BioEmu2 rather than BioEmu -1.

Many states were solved with a ligand, DNA, a partner protein or a phosphorylation that is not indicated in sampled sequences. Such a state can test whether a model given only the protein sequence reaches the conformation, but not its population, and each state was therefore given a condition label. A state was labelled apo-matched (A) when at least one reference structure was solved without a ligand, nucleic acid, partner, phosphorylation or mutation that the sampled sequence lacks, or when the same protein populates the state in solution. It was labelled condition -dependent (B) when every reference needed one of these condition s, and uncertain (C) when the evidence could not decide between the two.

The labels were assigned from the records of each reference file, and four were then revised after a review of the structural evidence (Table S2). The final labels gave 22 A, 24 B and three C states.

Population targets with a stated uncertainty were available for ABL1, PTP1B, SHP2 and Aurora A. Each was measured for an apo, unphosphorylated, single -chain protein that both models can represent.^2,24–26^ By contrast, the primary papers for T4L L99A and MBP give populations without a standard uncertainty,^27–29^ and only an upper bound is available for CypA.^30^ These three systems were compared with the models descriptively.

### 2.3 Sequences and Alignments

The benchmark systems were sampled with their published test-case sequences. The other systems were sampled with the construct of the solution measurement where one exists (for example, Aurora A residues 122 to 403). Residue numbers follow UniProt numbering throughout. One multiple sequence alignment per sequence was obtained from the public ColabFold MMseqs2 server^19^ and used as input for BioEmu-1, BioEmu2 and ColabFold. The number of sequences in each alignment ranged from 338 for EcoT38I to 24,091 for CypA.

### 2.4 BioEmu-1 and BioEmu2 Sampling

BioEmu-1 was run with bioemu 1.2.0 and its default checkpoint, bioemu-v1.1, using the default DPM-Solver sampler with 50 denoising steps. Its built-in sample filter was switched off, and the same filter was applied to the samples of both models at the scoring stage. BioEmu -1 writes only the backbone and Cβ atoms.

BioEmu2 was run from the preview checkpoint (bioemu2-preview, Hugging Face snapshot dd4b0c1b), the only BioEmu2 weights released as of 24 November 2026, with the code at repository commit 6d1ca744. BioEmu2 was run in equilibrium mode with the default unsteered sampler, a step scale of 1.0 and 200 denoising steps. This mode was chosen because it samples a range of conformations rather than sharpening the distribution towards one structure. ^10,12^ BioEmu2 also writes the side chains, and every sample it generated was kept for scoring.

Each model sampled each system as a single protein chain in three independent runs of 10,000 samples, and run k used the base seed k × 100,000.

ABL1 was also sampled under perturbations whose effect on the populations has been measured^2^ and with two drugs, imatinib and dasatinib, that bind one state selectively. Each condition was sampled in three runs of 10,000 samples with the settings of the wild-type runs. BioEmu2 was given 11 conditions, namely the mutants T408Y, Y272H, F378V, Y272H/T408Y, F378V/T408Y and H415P, phosphorylated Y412, the SH3-SH2-kinase construct (residues 83 to 534) alone and with the allosteric inhibitor GNF-5, and the kinase domain with imatinib or dasatinib. BioEmu-1 cannot take modified residues or ligands and was given the six mutants and the phosphomimetic substitution Y412E in place of phosphorylated Y412.

BioEmu2 can also propagate a structure forward by sampling new structures at a set nominal lag after a given start.^12^ This mode was run for ABL1 from 10 NMR models each of the active state (PDB ID: 6XR6), I1 (PDB ID: 6XR7) and I2 (PDB ID: 6XRG). The models were drawn at random from those that the ABL1 class rules assign to each state. The I2 models carry the substitutions G269E, M309L and T408Y. These substitutions were reverted to the wild type with PDBFixer before a restrained energy minimisation in OpenMM. From each start, 32 endpoints were sampled at nominal lags of 1, 10, 100 and 1,000 ns. One chain of 1,000 consecutive steps of 1 ns was also run from each of the first five starts drawn for each state, which gave 15 chains, and each step started from the previous frame.

### 2.5 Follow-Up BioEmu2 Runs with a Binding Partner or Phosphorylation

BioEmu2 was run again on seven proteins with the partner or phosphorylation of a missed state added. This was done to test whether those states were missed because the partner or modification was absent (Table S3).

Four proteins were given the molecules that touch the reference chain in their bound structure. DNA polymerase β (Pol β) was given the template and primer DNA, the nucleotide analogue ddCTP and one Mg²⁺ ion of its DNA ternary complex (PDB ID: 2BPG). NanR was run in two conditions, namely one with its operator DNA and one with sialic acid and Zn² ⁺. BusR was given DNA with cyclic di-AMP, and the LapD module was given two cyclic di-GMP with Mg²⁺. The reference chain also touches another protein chain in every bound structure except that of Pol β, and these runs did not include that second protein chain.

Three kinases were given the phosphorylation found in the crystal structures of their missed state, namely pT185 and pY187 for ERK2 (active state), pT180 and pY182 for p38α (active-like loop) and pY530 for SRC (inactive αC-out state). Y530 lies in the C-terminal tail that holds SRC inactive.

Each condition kept the construct, alignment, reference states, scored residues, thresholds and model settings of its protein-only condition and was sampled in three runs of 1,000 samples. This smaller budget was used because BioEmu2 had found every state that it reached within about 500 samples in the protein-only runs. Only the protein chain was scored. The phosphorylated residues in ERK2 and p38α lie in the scored loop and were scored as their parent residues in both the modified and the unmodified sequence.

A state was counted as successfully reproduced when a filter-passing sample came within 2.0 Å Cα RMSD of it in at least two of the three runs. This threshold is stricter than the 3.0 Å used for recovery, because these runs ask whether the bound conformation itself is reproduced. As the protein-only runs had ten times as many samples, they were also compared with the follow-up runs at 1,000 samples (Supporting Information).

### 2.6 ColabFold

ColabFold 1.5.4,^19^ the version bundled with bioemu 1.2.0, was run on all 22 systems as a structure-prediction reference. Each system was predicted with the five AlphaFold2 models ^5^ (model type alphafold2_ptm) and 100 random seeds, which gave 500 ColabFold models per system. The alignment was subsampled with the max-msa setting 32:64, which keeps at most 32 cluster centres and 64 extra sequences, because such subsampling can lead AlphaFold2 to produce more than one conformation.^6^ Two shallower depths (16:32 and 8:16) were also run for ABL1. Templates, dropout and Amber relaxation were not used, and every prediction ran the default three recycles.

### 2.7 All-Atom Molecular Dynamics Simulations

All-atom MD was run for five systems to test whether each end state is stable on a time scale that all-atom simulation can reach. These were ABL1, whose state populations and their response to perturbation have been measured, three systems whose states differ in side-chain or loop detail that only an all-atom model resolves (T4L L99A, CypA and PTP1B), and AdK as the reference case.

The simulations contained the protein only, as in the generative runs. Crystal ligands such as Ap5A in closed AdK, water molecules and hydrogen atoms were removed from the starting structures. Each protein was described with the ff19SB force field ^31^ and solvated with the OPC four-point water model^32^ in a truncated octahedral box with at least 10 Å of water around the protein. The systems were neutralised with Na⁺ or Cl⁻ ions, and NaCl was then added to about 0.15 M, with the number of ion pairs estimated from the number of water molecules. They were built with pdb4amber and tleap from AmberTools 22.3^33^ and simulated with pmemd.cuda from Amber 20.^34^ Hydrogen mass repartitioning^35^ with ParmEd set the mass of every non-water hydrogen atom to 3.024 Da, which allowed a 4 fs time step.

Each replica was minimised twice for up to 5,000 steps, first with the protein heavy atoms restrained at 10 kcal mol⁻¹ Å⁻² and then without restraints. Each replica was then heated at constant volume from 0 K to its simulation temperature over 80 ps and held there for 20 ps, with heavy-atom restraints of 5 kcal mol⁻¹ Å⁻², a Langevin collision frequency of 2 ps⁻¹ and a 2 fs time step. The restraints were released at constant pressure in four steps of 250 ps (5.0, 2.0, 1.0 and 0.5 kcal mol ⁻¹ Å⁻²). Unrestrained equilibration at constant pressure then ran for 2 ns with a 4 fs time step before production.

Production runs used a Langevin thermostat with a collision frequency of 1 ps⁻¹, a Monte Carlo barostat at 1 bar with a relaxation time of 1 ps, particle-mesh Ewald summation with a 9 Å direct-space cut-off and SHAKE on bonds to hydrogen. ABL1 and CypA were simulated at 283.15 K and T4L L99A at 298.15 K. These temperatures were chosen to match the temperatures of their NMR measurements. PTP1B and AdK were simulated at 300 K, because no measured population is used for AdK and the PTP1B measurement was made near room temperature. Three replicas of 250 ns were started from each end state. These simulations gave 30 replicas and 7.5 µs in total. Frames were saved every 100 ps. The first 10 ns of each replica were treated as further equilibration and removed before scoring, which left 2,400 frames per replica.

### 2.8 Scoring

All samples, ColabFold models and MD frames were scored with one scoring package built on MDTraj^36^ (Figure 1b). Scoring had four steps. First, each sample was mapped to UniProt numbering, superposed on a fixed set of superposition residues of its system and compared with every reference structure by the Cα root-mean-square deviation (RMSD) over the scored residues. The distance to a state is the smallest RMSD to any of its references. Where states differ only locally, only the moving region was scored, for example the WPD loop of PTP1B (residues 177 to 185) or a kinase activation loop after superposition on the C-lobe.

Second, every sample was checked with the backbone filter that the BioEmu-1 developers apply to its output by default, as implemented in bioemu 1.2.0. ^8,37^ This filter was used for all methods because it defines the samples that a BioEmu-1 user normally receives. A sample passed when every consecutive Cα–Cα distance was below 4.5 Å, every peptide C–N distance was below 2.0 Å and no two N, Cα, C, O or Cβ atoms of residues three or more apart were closer than 1.0 Å. These criteria detect broken peptide bonds, gaps between residues and overlapping atoms. Only samples that passed (filter-passing samples) were used for recovery and populations. The failure fraction of a system is the percentage of all generated samples that failed the filter, out of 30,000 for each BioEmu model (three runs pooled) and 500 for ColabFold.

Two further geometry checks were made on 1,000 filter-passing samples per run. Their backbone bond lengths, bond angles and Ramachandran angles were compared with standard values.^38,39^ The stricter HIGH_QUALITY check that BioEmu2 applies to its own output ^10^ was also applied to the samples of both models. This check rejects a sample with more than five backbone bonds outside set length ranges or with too many clashes or other bonding errors. The side chains of BioEmu-1 samples were first rebuilt with HPacker^40^ for this check (Supporting Information).

Third, each filter-passing sample was assigned to its nearest state if its RMSD to that state was below the class threshold, which is the largest distance at which a sample counts as being in a state. Otherwise, it was assigned to the unassigned class of its system. The class threshold was 3.0 Å, as for recovery. Meanwhile, for MBP, PTP1B and T4L L99A, the class threshold was 1.5 Å because their closest states are only 2.3 to 3.1 Å apart.

Five systems were assigned by other rules for the population comparisons with measurements, because their measured states are not defined by the backbone RMSD alone. These rules used a geometric measure, such as a distance or a dihedral angle, as a collective variable.

The open state of SHP2 has no single crystal arrangement, because its N-terminal SH2 (N-SH2) domain is released from the phosphatase (PTP) domain and is mobile. SHP2 samples were therefore classed as open when the shortest Cα–Cα distance between these two domains was above 10 Å. The Aurora A measurement reports whether two dye-labelled residues touch, so Aurora A samples were classed as closed when the Cα distance between K224 and S283 was below 22 Å, a value in the gap between the folded and extended crystal structures. CypA and T4L L99A were assigned using the χ1 rotamer of Phe113 and of Phe114, respectively, which was read from the rebuilt side chains for BioEmu-1. ABL1 was assigned from three backbone measures, as its active state and I1 lie only 1.71 Å apart. These were the φ angle of D400 in the DFG motif (the Asp-Phe-Gly sequence at the start of the activation loop), the ψ angle of A399 and the Cα distance between D344 and L406.

Fourth, the recovery of a state at a sampling budget b and threshold t was computed as the probability that a random subset of b samples from one run contains at least one filter-passing sample within t of the state. In other words, it is the chance that a user who draws b samples finds at least one of them within t of the state. The subset is drawn from all samples of the run, including those that fail the filter. Recovery was computed exactly from the hypergeometric distribution. It was averaged over runs and then over the states of a system, at budgets from 100 to 10,000 samples and thresholds of 1.5, 2.0 and 3.0 Å.

The population of a class is the percentage of filter-passing samples assigned to it in one run. The populations were renormalised over the measured states where those states sum to one, and differences between populations are given in percentage points (pp). A run matched a measured population when its value lay within the population tolerance, which is twice the reported uncertainty.

As an independent check of our scoring, the runs of the 12 benchmark-derived systems (AdK and the 11 benchmark systems) were also scored with the evaluator of the bioemu-benchmarks package,^22^ which the BioEmu developers released with these test cases. This evaluator compares samples with the references by backbone RMSD over the N, Cα, C and O atoms and uses its own sample filter and scored regions.

### 2.9 Statistical Analysis

Our first comparison was a binary outcome for each system, that is, whether or not a model reached every state of the system successfully. A state is counted as reached when at least two of the three runs of 10,000 samples contained a filter-passing sample within 3.0 Å of it. This threshold was used because Lewis et al. reported the domain-motion recovery of BioEmu-1 at 3.0 Å,^8^ so our results are comparable to theirs.

The two models were compared with the exact McNemar test for paired binary outcomes, ^41^ which uses only the systems in which one model reached every state and the other did not. No interval is given for the difference in the proportion of systems recovered. This is because an interval over systems would ignore the nesting of systems in families. The test was repeated at 1.5, 2.0 and 3.0 Å with three run rules (any run, two of three runs and all three runs), and the nine *p* values were adjusted with Holm’s method.^42^

A paired test of binary outcomes has little power with 22 systems, so graded recovery was also compared. Graded recovery is the recovery averaged over the states of a system, and it was compared as the paired difference per system (BioEmu2 minus BioEmu-1) at each budget and threshold. The same difference was computed for the budget-averaged recovery, which is the graded recovery at budgets of 100, 500, 1,000, 3,000 and 10,000 samples, averaged on a logarithmic scale of the budget by the trapezoid rule. On this scale, the budgets from 100 to 1,000 samples carry about the same weight as those from 1,000 to 10,000 samples.

The systems were grouped into 14 families by homology (Table 1). The 95% percentile confidence interval (CI) of the mean difference over systems came from a two-stage bootstrap with 20,000 resamples that drew families first and then systems within them, as systems are nested within families. The *p* values came from an exact two-sided sign-flip test on the 14 family means, which weights each family equally. This test enumerates all 16,384 possible sign patterns, and its smallest attainable *p* value is 1.2 × 10⁻⁴. The failure fraction was analysed in the same way, and no correction for multiple comparisons was applied to these secondary comparisons.

To test whether one family drove the result, the comparisons at 3.0 Å were repeated with each family left out in turn and without CypA, whose two backbone states are 0.11 Å apart. The graded comparison was also repeated in three other ways, with 20 finer family labels based on Pfam domains, with a criterion that also requires the state to be the nearest one, and without AdK, which BioEmu2 may have seen during training. Results were also split by condition label and by origin. State counts by label are given without a statistical test, because the states of one protein are not independent.

The variation of a class population between runs was measured as its standard deviation (SD) over the three runs of a model. For each model, the width of the interval that holds the middle 90% of its samples (from the 5th to the 95th percentile) was measured along the variable that separates the states of a protein. The populations of the two models within a system were compared on the class whose population differed most between them, as the BioEmu2 population minus the BioEmu-1 population. This class was chosen at the class threshold of each system and then compared at the other thresholds tested (2.0, 2.5, 3.5 and 4.0 Å, or three or four values from 1.25 to 2.5 Å for the systems with a 1.5 Å threshold). Populations were compared with measurements system by system rather than pooled, as only three systems outside ABL1 had a target with a stated uncertainty.

Populations were also converted to apparent free-energy differences at 298.15 K, a common reference temperature for all systems, as ΔG = −RT ln(p₁/p₂) from the mean population over runs. Where a method did not place any filter-passing sample in a state, the exact one-sided 95% upper bound on its population, 1 − 0.05^(1/n) for n filter-passing samples pooled over the three runs, gave a bound on ΔG. These free-energy differences are apparent, because sampling frequencies have not been shown to be Boltzmann probabilities. For the ABL1 perturbations, a change in population was called only when the three run values of the perturbed condition lay entirely abo ve or below those of its comparison condition.

Finally, three measures described the breadth of each ensemble, that is, how widely a model spreads its samples. The reference spread is the median of the largest difference between the RMSDs of two samples to any one reference state, taken over 20,000 random pairs of filter-passing samples. It is computed from the RMSDs directly and does not depend on the class threshold. The pairwise RMSD is the median pairwise Cα RMSD of the scored residues among 1,000 filter-passing samples per run, and it needs no reference structure. The cluster count is the number of clusters at 3.0 Å found with the method of Daura et al.^43^ The log ratio of BioEmu2 to BioEmu-1 was tested with the sign-flip test on the 14 family means for the reference spread and the pairwise RMSD.

### 2.10 Software and Hardware

Scoring used be_scoring 0.1.0 with Python 3.12, MDTraj 1.11.2, NumPy 2.3.5, SciPy 1.18.1 and pandas 3.1.0. The independent check used bioemu-benchmarks 0.0.2.post1 with the multi-conformation test cases of version 0.1. BioEmu2 ran with Python 3.12 and PyTorch 2.13.0 on NVIDIA A100-SXM4-80GB GPUs, and parts of the CypA and T4L L99A runs ran on V100 GPUs with the same installation. BioEmu-1 ran with Python 3.11 and PyTorch 2.13.0, and BioEmu-1, ColabFold and the all-atom simulations ran on NVIDIA Tesla P100-PCIE-12GB GPUs. All software versions and identifiers are listed in Table S17.

## 3. Results and Discussion

### 3.1 Backbone Geometry of the Generated Samples

Before a sample can be compared with an experimental state, it must be a physically plausible structure. This is because a broken or clashing chain cannot count as reaching a state. Therefore, we started the work by counting the samples of each model that fail the backbone filter of BioEmu-1. We found that BioEmu-1 failed in a median of 5.60% of samples per system, from 0.81% for T4L L99A to 35.08% for SHP2. In contrast, BioEmu2 failed in a median of 0.010% of samples, and its failure fraction was lower in all 22 systems (Figure 2a). On average, the failure fraction of BioEmu2 was 8.23 pp lower than that of BioEmu-1 (95% CI 4.57 to 13.27 pp), and the ColabFold models failed in 0% to 2.2% of cases. As BioEmu-1 removes failing samples by default, a user would need about 6% more samples for the median system, and about 54% more for SHP2, to obtain the same number of filter-passing samples.

**Figure 2.**
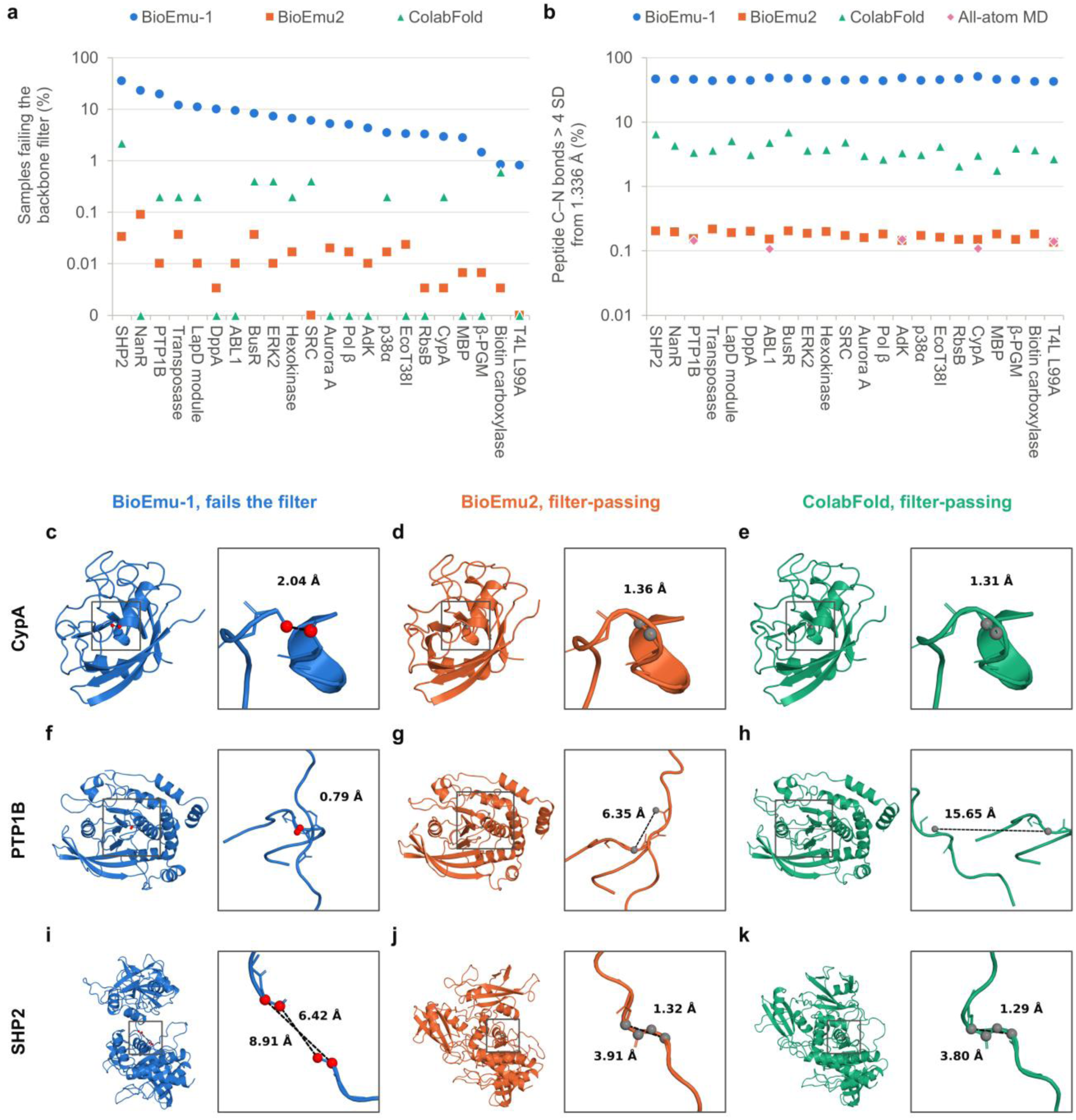
Backbone geometry of the generated samples. (a) Failure fraction under the backbone filter for each system for BioEmu-1 and BioEmu2 (30,000 samples each, three runs pooled) and ColabFold (500 models per system), on a logarithmic axis. Systems are ordered by the BioEmu -1 value. (b) Percentage of peptide C–N bonds more than 4 standard deviations from the standard length of 1.336 Å in filter-passing samples (1,000 samples per run, three runs pooled) for BioEmu-1, BioEmu2, all-atom MD (five systems) and ColabFold. (c–k) A BioEmu-1 sample that fails the filter (blue), a filter-passing BioEmu2 sample (orange) and a filter-passing ColabFold model (green) for CypA (c–e), PTP1B (f–h) and SHP2 (i–k). Red spheres mark the two atoms at the failure site, and grey spheres mark the same atoms in the other two structures. Each panel shows the whole structure on the left and an enlarged view of the failure site on the right. The grey square marks the enlarged region, which shows the distance between the marked atoms. The BioEmu2 sample and the ColabFold model were drawn at random from the filter-passing run-1 samples or models and superposed on the BioEmu-1 sample.

To show what these failures look like, we took one failing BioEmu-1 sample from each of three proteins that span the range of chain lengths, namely CypA (165 residues), PTP1B (321) and SHP2 (529). We found a stretched peptide bond in CypA (C–N distance 2.04 Å), a clash between two distant parts of the chain in PTP1B (two atoms 0.79 Å apart) and a gap in the chain of SHP2 (Cα–Cα distance 8.91 Å; Figure 2c–k). In a BioEmu2 sample and a ColabFold model of each protein, the same atoms lay at normal distances.

However, passing the filter does not guarantee a realistic backbone, because the filter accepts any peptide C–N distance below 2.0 Å, whereas the standard length is 1.336 Å. ^38^ Therefore, we compared two measures of the filter-passing samples with standard values that have a known spread, namely the peptide C–N bond (1.336 ± 0.023 Å), which links one residue to the next, and the N–Cα–C angle (111.0 ± 2.7°), which lies within a residue.^38^ In BioEmu-1, a median of 45.3% of peptide C–N bonds per system lay more than 4 SDs from 1.336 Å, compared with 0.18% in BioEmu2, 0.14% in all-atom MD and 3.7% in ColabFold (Figure 2b, Table S6). Their mean length was on target (1.337 Å), so these bonds were scattered around the correct length rather than shifted from it.

Within residues, we saw the reverse. BioEmu-1 kept the N–Cα–C angle almost fixed (median SD 0.83°), whereas BioEmu2 (3.3°) and all-atom MD (3.5°) varied it by about the spread of the standard value. This is consistent with BioEmu-1 building each residue as a rigid unit without constraining the bond to the next residue, so that its errors gather at the junctions between residues. On both measures, the local geometry of BioEmu2 therefore resembled that of all-atom MD, with 0.18% against 0.14% of peptide C–N bonds beyond 4 SDs and a median N–Cα–C angle SD of 3.3° against 3.5° (Table S6).

Meanwhile, the stricter HIGH_QUALITY check of BioEmu2 gave the same picture. BioEmu2 marked at least 99.98% of its own samples as passing in every system, whereas only a median of 23.4% of BioEmu-1 samples passed once we had rebuilt their side chains (Table S7). Most of these failures came from the BioEmu-1 backbone, which the rebuild leaves unchanged, and a control rebuild of BioEmu2 samples suggested that the rebuild itself contributed only a small part of the failures (Supporting Information). However, BioEmu2 samples contain errors of their own. A cis peptide bond before a residue other than proline is rare in experimental structures (43 such bonds in a set of 571 proteins),^44^ and such bonds made up at most 0.033% of the BioEmu2 peptide bonds in every system except Pol β. In Pol β, 2,984 of the 3,000 BioEmu2 samples that we checked had at least one such bond. In all 50 of these samples that we inspected, it was the Gly274 –Ser275 bond, which is not cis in either of the two Pol β reference structures (the apo enzyme, PDB ID: 1BPD, and the DNA complex, PDB ID: 2BPG).

Taken together, these results suggest that BioEmu2 has largely removed two defects of BioEmu-1, namely the broken and clashing backbones that fail the filter and the distorted peptide bonds that remain in BioEmu-1 samples that pass it. However, BioEmu2 can still make errors, such as the cis peptide bond that it placed in almost every Pol β sample that we checked, so it is still worth checking the local peptide geometry of its samples.

### 3.2 Recovery of the Experimental States

A cleaner backbone is useful only if the samples also reach the biologically relevant conformations that a protein adopts. To test whether BioEmu2 outperforms BioEmu-1 in reaching the 49 experimental states included in our study, we asked whether a model reached every one of its states (the binary outcome) for each system.

In our hands, BioEmu-1 reached every state in 15 of the 22 systems (68%) and BioEmu2 in 10 (45%; Table 2, Figure 3e). The 10 systems in which BioEmu2 reached every state were all among the 15 systems of BioEmu-1. Likewise, the seven systems in which BioEmu-1 failed (ABL1, Aurora A, BusR, EcoT38I, SHP2, SRC and the transposase) were all systems in which BioEmu2 also failed. The two models therefore disagreed only in five systems (ERK2, NanR, p38α, Pol β and the LapD module), and in each of them BioEmu-1 succeeded and BioEmu2 did not. However, this difference was not statistically significant (*p* = 0.0625), because only five systems separated the two models, which is too few for this test to reach *p* < 0.05.

**Figure 3.**
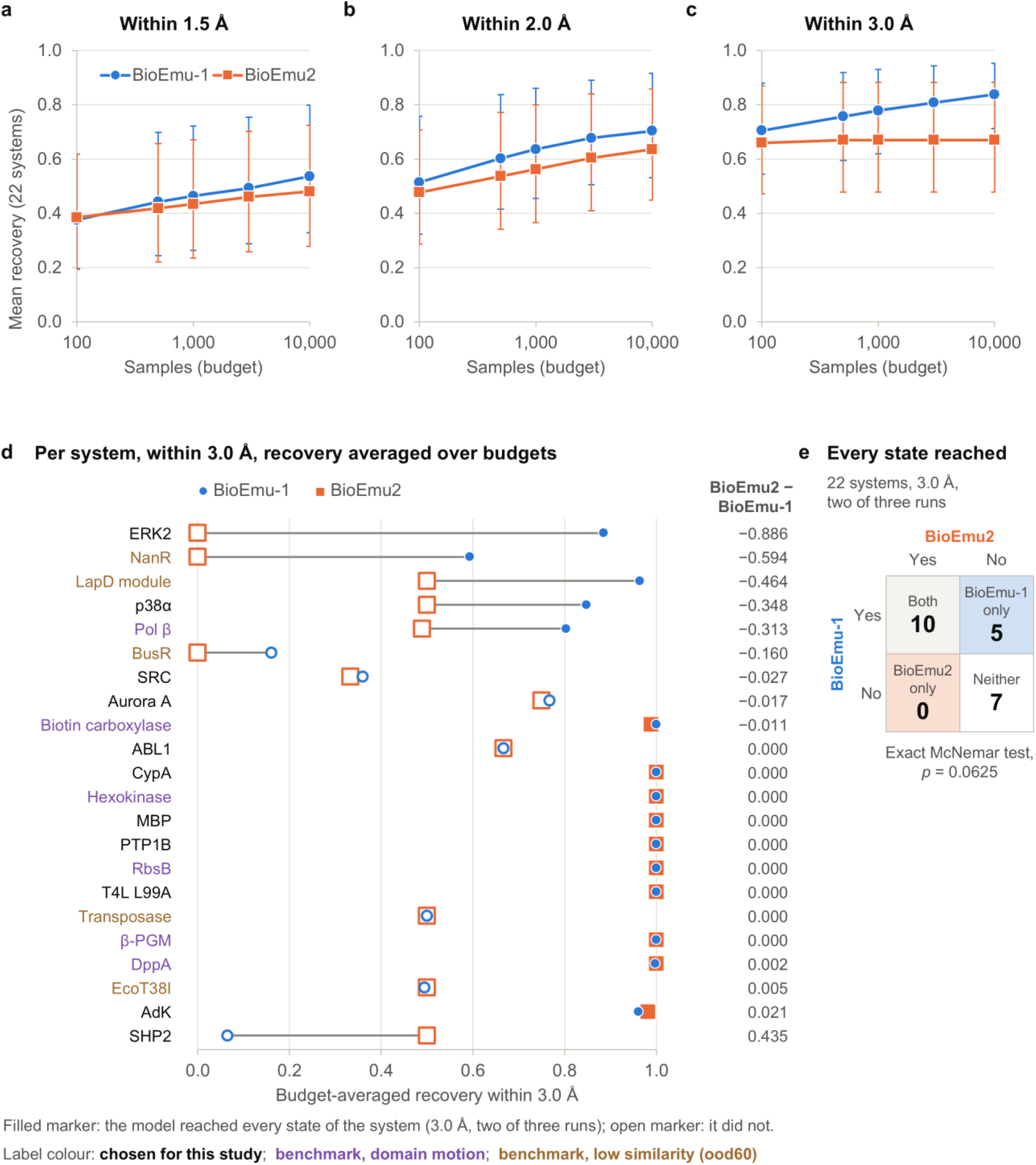
Recovery of the experimental states against sampling budget. (a–c) Mean graded recovery over the 22 systems for BioEmu-1 and BioEmu2 at thresholds of 1.5, 2.0 and 3.0 Å and budgets of 100, 500, 1,000, 3,000 and 10,000 samples on a logarithmic axis. Graded recovery is the probability that a random subset of the given size from one run contains a filter-passing sample within the threshold of a state, averaged over runs and states. Error bars are 95% two-stage family-bootstrap intervals of each mean (20,000 resamples). (d) Budget-averaged recovery within 3.0 Å for each system, with the two models joined by a line and their difference (BioEmu2 minus BioEmu -1) on the right. Systems are ordered by this difference. A filled marker means that the model reached every state of the system within 3.0 Å in at least two of three runs, and the colour of each label gives the origin of the system. (e) Classification of the 22 systems by whether each model reached every state within 3.0 Å in at least two of three runs.

**Table 2.** Number of systems in which every state was reached, under three thresholds and three run rules.

| **Threshold (Å)** | **Run rule** | **BioEmu-1** | **BioEmu2** | **Both** | **Neither** | **BioEmu-1 only** | **BioEmu2 only** | ***p*** | **Holm-adjusted *p*** |
|---|---|---|---|---|---|---|---|---|---|
| 1.5 | Any run | 10 | 7 | 6 | 11 | 4 | 1 | 0.375 | 1.00 |
| 1.5 | 2 of 3 runs | 9 | 6 | 6 | 13 | 3 | 0 | 0.250 | 1.00 |
| 1.5 | All 3 runs | 8 | 6 | 6 | 14 | 2 | 0 | 0.500 | 1.00 |
| 2.0 | Any run | 13 | 9 | 9 | 9 | 4 | 0 | 0.125 | 0.875 |
| 2.0 | 2 of 3 runs | 11 | 9 | 9 | 11 | 2 | 0 | 0.500 | 1.00 |
| 2.0 | All 3 runs | 11 | 9 | 9 | 11 | 2 | 0 | 0.500 | 1.00 |
| 3.0 | Any run | 16 | 10 | 10 | 6 | 6 | 0 | 0.031 | 0.281 |
| 3.0 | 2 of 3 runs | 15 | 10 | 10 | 7 | 5 | 0 | 0.0625 | 0.500 |
| 3.0 | All 3 runs | 14 | 10 | 10 | 8 | 4 | 0 | 0.125 | 0.875 |

Counts are the number of systems (of 22) in which every state has a filter-passing sample within the threshold at 10,000 samples. The *p* values are from the exact McNemar test, and the Holm-adjusted *p* values are across the nine combinations. The two-of-three rule at 3.0 Å is the setting used throughout, and the other eight combinations test the dependence on this choice. The system reached only by BioEmu2 at 1.5 Å in any run is AdK. Without CypA, which both models reached, the BioEmu-1, BioEmu2 and Both counts each fall by one, and the *p* values are unchanged.

To test whether this result depended on the 3.0 Å threshold or on requiring two of the three runs, we repeated the comparison at 1.5, 2.0 and 3.0 Å with three different run rules. BioEmu -1 succeeded in more systems than BioEmu2 across all nine combinations. However, after Holm’s adjustment, none of the differences was statistically significant (Table 2). When we left out one protein family at a time, or CypA alone, the *p* values ranged from 0.0625 to 0.25 (Supporting Information).

Nevertheless, the binary outcome does not show how many samples a model needs to reach a state, or whether it reached some but not all the states of a system. Therefore, we also compared graded recovery against the sampling budget (Figure 3a–c, Table S8). For each threshold, we summarised the comparison by the budget-averaged recovery, which averages graded recovery over budgets from 100 to 10,000 samples. We found that the budget-averaged difference (BioEmu2 minus BioEmu-1) was negative at every threshold. The results were −0.026 (95% CI −0.127 to 0.060) at 1.5 Å, −0.064 (−0.153 to 0.042) at 2.0 Å and −0.107 (−0.244 to 0.028) at 3.0 Å. Because recovery is a probability, the difference of −0.107 at 3.0 Å has a direct meaning. Averaged over states and sampling budgets, the chance that a set of BioEmu2 samples contained a sample within 3.0 Å of a state was 0.107 lower (10.7 percentage points) than for a set of BioEmu-1 samples of the same size. Although these values favour BioEmu-1, every confidence interval included zero, so we cannot conclude that either model recovers states better. In two of four sensitivity analyses, the 3.0 Å interval excluded zero in favour of BioEmu-1, but these intervals come from a bootstrap over few families and may be too narrow (Supporting Information). These differences agree with the BioEmu2 developers’ comparison against MD simulations, where BioEmu-1 outperformed BioEmu2 on single proteins.^12^

We also found that the two models responded differently to more sampling. At 3.0 Å, the mean recovery of BioEmu2 was 0.660 at 100 samples and stayed at 0.670 at every budget from 500 to 10,000 samples, whereas that of BioEmu-1 rose from 0.706 to 0.838 (Figure 3c). The difference between the models therefore widened from −0.046 at 100 samples to −0.168 at 10,000 samples. This suggests that BioEmu2 found the states that it reaches within a few hundred samples, and that further sampling did not add any new state at 3.0 Å.

To find out which systems produced these averages, we compared the budget-averaged recovery within 3.0 Å system by system (Figure 3d). In 15 of the 22 systems, the two models differed by less than 0.03. Among the other seven, BioEmu-1 had the higher recovery in six (ERK2, NanR, the LapD module, p38α, Pol β and BusR), by 0.160 to 0.886, whereas BioEmu2 had the higher recovery only in SHP2, by 0.435. At 2.0 and 3.0 Å, BioEmu-1 had the higher budget-averaged recovery in each of the three groups of systems (study, domain-motion and ood60), and it remained ahead when we left out AdK, which BioEmu2 may have seen during training (Supporting Information). BioEmu-1 also reached more states than BioEmu2 when we counted only its samples that passed the stricter HIGH_QUALITY check of BioEmu2. In random subsets of 1,000 filter-passing samples per run, these samples reached 36 of the 49 states in at least two of the three runs, against 33 for BioEmu2 (Supporting Information).

As an outside reference, we compared both models with ColabFold at an equal number of samples. Within 3.0 Å, the expected number of states recovered in 500 samples was 36.9 for BioEmu-1 and 33.0 for BioEmu2, against 36 for the 500 ColabFold models. Therefore, we concluded that neither generative model recovered clearly more states than ColabFold at this budget (Supporting Information).

To check our scoring, we also scored the 12 benchmark-derived systems with the evaluator that the BioEmu developers released with the bioemu-benchmarks test cases.^22^ The evaluator agreed with our scoring on all 144 reached or not-reached calls at 3.0 Å. Its coverage, a bootstrapped estimate of the fraction of domain-motion reference structures that a model samples within 3.0 Å, was 0.825 to 0.841 per run for BioEmu-1. This range is close to the 83% reported by Lewis et al.,^8^ although the weights, test cases and success rule differ (Table S5).

Taken together, these results suggest that the cleaner geometry of BioEmu2 did not translate into reaching more biologically relevant states, and that sampling BioEmu2 beyond a few hundred samples did not add any new state at 3.0 Å.

### 3.3 States Missed by BioEmu2

To find out which states BioEmu2 missed, we compared the closest filter-passing sample of each model over its three runs for each of the 49 states (Figure 4a). We counted a state as reached when at least one run came within 3.0 Å of it. Using this rule, BioEmu2 reached 33 states and missed the other 16 (Table 3). BioEmu-1 also reached all 33 of these states, as well as 10 of the 16 states that BioEmu2 missed, whereas neither model reached the remaining six. BioEmu2 therefore did not reach any state that BioEmu-1 missed, so it did not gain any state in exchange for those that it missed. One example state from each occupied quadrant of this comparison shows what reaching and missing a state look like (Figure 4c).

**Figure 4.**
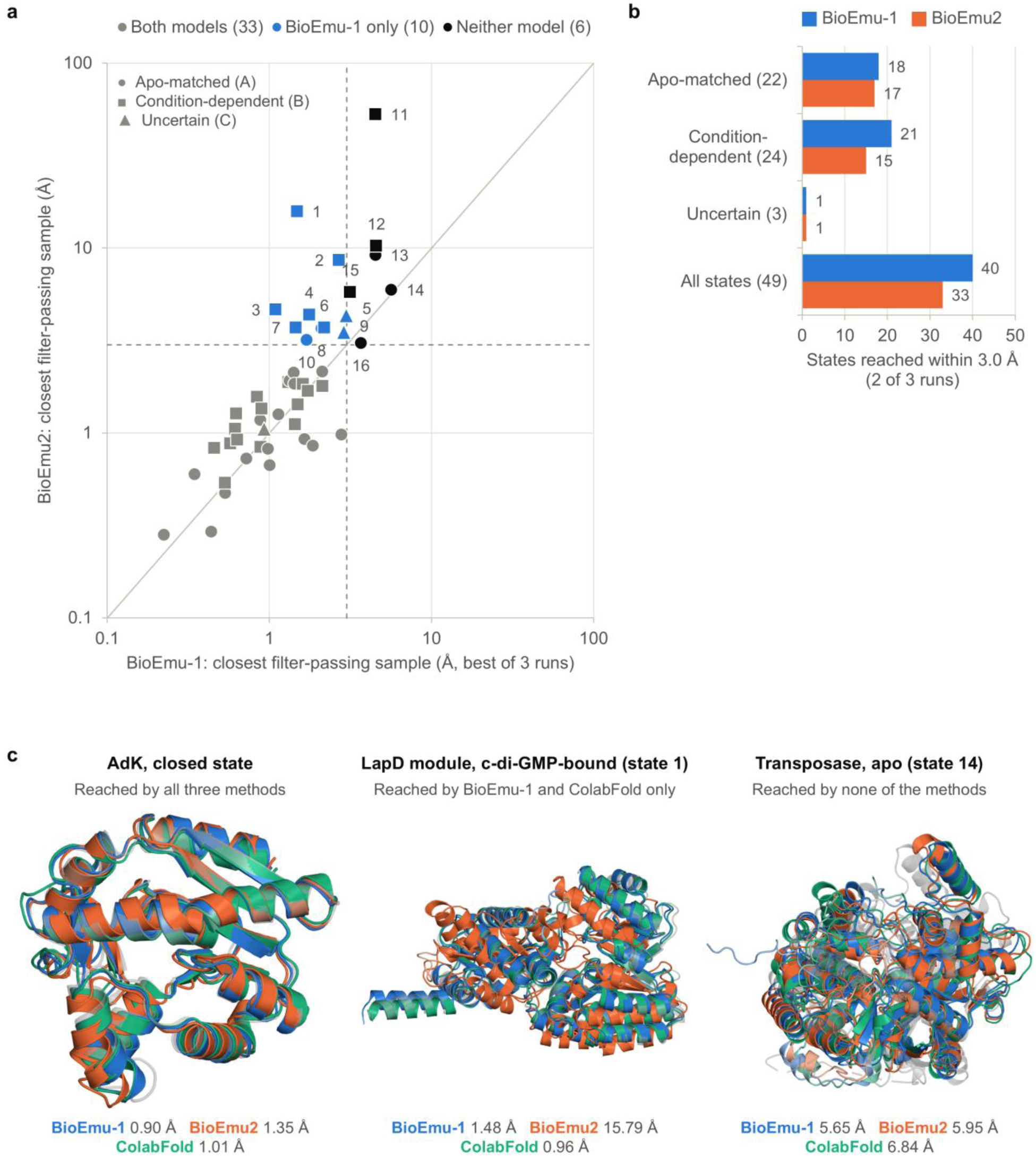
States reached by each model. (a) Distance from each of the 49 states to the closest filter-passing sample of BioEmu-1 and BioEmu2, best of three runs, on logarithmic axes. Dashed lines at 3.0 Å divide the panel into states reached by both models (lower left), by BioEmu-1 only (upper left), by BioEmu2 only (lower right, none) and by neither model (upper right), and the solid line marks equal distances. Colour shows which model reached the state within 3.0 Å in any run, and marker shape shows the condition label (circles for apo-matched, squares for condition-dependent and triangles for uncertain states). Numbers are the state numbers of Table 3. (b) Number of states reached within 3.0 Å in at least two of three runs, grouped by condition label. (c) One example state from each occupied quadrant of (a). Each image shows the closest filter-passing sample of BioEmu-1 (blue), BioEmu2 (orange) and ColabFold (green) superposed on the reference structure (grey), with the scored residues in full colour, and the numbers give the Cα RMSD over the scored residues.

**Table 3.** The 16 states not reached within 3.0 Å by at least one BioEmu model in any run.

| No. | System · state | Reference structures (PDB IDs) and condition | Label | BioEmu-1 | BioEmu2 |
|---|---|---|---|---|---|
| 1 | LapD module · c-di-GMP-bound | 6PWK; second messenger bound | B | 1.48 (3) | 15.79 (0) |
| 2 | BusR · DNA-bound a | 7OZ3; DNA and c-di-AMP bound | B | 2.69 (3) | 8.62 (0) |
| 3 | p38α · active-like loop | 3PY3, 3OD6; phosphorylation-dependent | B | 1.09 (3) | 4.65 (0) |
| 4 | NanR · DNA-bound | 6WFQ; DNA bound | B | 1.77 (3) | 4.36 (0) |
| 5 | SRC · NMR intermediate | 9NS1; solution NMR state | C | 2.99 (1) | 4.32 (0) |
| 6 | NanR · sialic-acid-bound | 6ON4; ligand bound | B | 2.19 (2) | 3.72 (0) |
| 7 | ERK2 · active | 2ERK, 6OPG, 5V61; dually phosphorylated | B | 1.46 (3) | 3.72 (0) |
| 8 | ERK2 · inactive | 8AOJ, 4GT3; unphosphorylated, with an ATP-site ligand in both | A | 2.10 (3) | 3.68 (0) |
| 9 | Aurora A · inactive, folded loop | 2WTV, 4J8M | C | 2.88 (1) | 3.51 (0) |
| 10 | Pol β · apo | 1BPD; apo | A | 1.69 (3) | 3.19 (0) |
| 11 | SHP2 · open | 6CRF; activating mutant E76K | B | 4.52 (0) | 52.85 (0) |
| 12 | BusR · DNA-bound b | 7B5Y; DNA and c-di-AMP bound | B | 4.56 (0) | 10.27 (0) |
| 13 | ABL1 · I2 | 6XRG; solution NMR state | A | 4.51 (0) | 9.17 (0) |
| 14 | Transposase · apo | 6PQN; apo | A | 5.65 (0) | 5.95 (0) |
| 15 | SRC · inactive, αC-out | 2SRC, 2H8H, 1KSW; needs phosphorylated Y530 | B | 3.15 (0) | 5.79 (0) |
| 16 | EcoT38I · apo | 7E8R; apo | A | 3.67 (0) | 3.07 (0) |

Distances are those of the closest filter-passing sample over all runs (Å), with the number of runs (of three) within 3.0 Å given in brackets. States 1 to 10 were reached by BioEmu-1 only and states 11 to 16 by neither model. Label A, apo-matched; B, condition-dependent; C, uncertain (Table S2).

The binary outcome used a stricter rule, under which a state is counted as reached only when at least two of the three runs come within 3.0 Å of it. Under this stricter rule, 32 states were reached by both models, eight only by BioEmu-1, one only by BioEmu2 and eight by neither model. The single state reached only by BioEmu2 was the closed state of SHP2, in which the N-SH2 domain blocks the active site of the phosphatase domain. BioEmu-1 reached this state in only one of its three runs, so it is counted as reached by both models under the first rule but only by BioEmu2 under the stricter rule.

However, we reasoned that a missed state is not always an error, because many states were solved with a ligand, nucleic acid, partner or phosphorylation that the sampled sequence lacks. Nine of the 16 states that BioEmu2 missed were condition-dependent, so these misses are not necessarily errors of the model. Having said that, BioEmu2 also missed five apo-matched states, namely unphosphorylated ERK2, apo Pol β, apo EcoT38I, the apo state of the transposase and the I2 state of ABL1, which wild-type ABL1 populates in solution. BioEmu-1 reached unphosphorylated ERK2 and apo Pol β, but it also missed the other three.

To test whether the advantage of BioEmu-1 lay mainly in condition-dependent states, we counted the states that each model reached by condition label (Figure 4b). Within 3.0 Å in at least two of three runs, BioEmu-1 reached 21 of the 24 condition-dependent states, whereas BioEmu2 reached 15. On the 22 apo-matched states, in contrast, the two models were close, with BioEmu-1 reaching 18 and BioEmu2 reaching 17.

We saw the same picture in graded recovery. On the apo-matched states, BioEmu2 had the higher budget-averaged recovery at 1.5 Å (0.568 against 0.448) and 2.0 Å (0.622 against 0.576), and the two were about equal at 3.0 Å (0.772 against 0.778). On the condition-dependent states, the recovery of BioEmu2 was lower at every threshold, for example 0.623 against 0.809 at 3.0 Å. When only the apo-matched states had to be reached, BioEmu-1 reached every such state in 12 of the 16 systems that contain one, and Bio Emu2 in 11 (*p* = 1.00). The six systems in which BioEmu-1 had the clearly higher recovery fit this pattern (Figure 3d).

Taken together, these results suggest that the gap between the two generations lies mainly in states that need a ligand, nucleic acid, partner or phosphorylation that the sampled sequence lacks, whereas the two models perform similarly on apo-matched states.

### 3.4 Breadth of the Ensembles and Placement of Domains

A model can miss a state for two different reasons. First, it may spread its samples too narrowly to reach the state. Second, it may sample broadly but in the wrong places. At 3.0 Å, the recovery of BioEmu2 stopped rising after about 500 samples, which pointed to the first reason. To understand this further, we measured how widely each model spreads its samples with three measures of breadth, namely (a) the reference spread, (b) the pairwise RMSD and (c) the cluster count.

We first measured the reference spread. For two samples, this measure is the largest difference between their RMSDs to the same reference state. Therefore, it is small when the samples of a model all lie at similar distances from every state and large when they lie at different distances. We found that the reference spread of BioEmu2 was smaller than that of BioEmu -1 in 17 of the 22 systems (median ratio of BioEmu2 to BioEmu-1 of 0.70, *p* = 0.018, Figure 5a). In the other five systems, however, the ratio ranged from 1.00 to 2.51.

**Figure 5.**
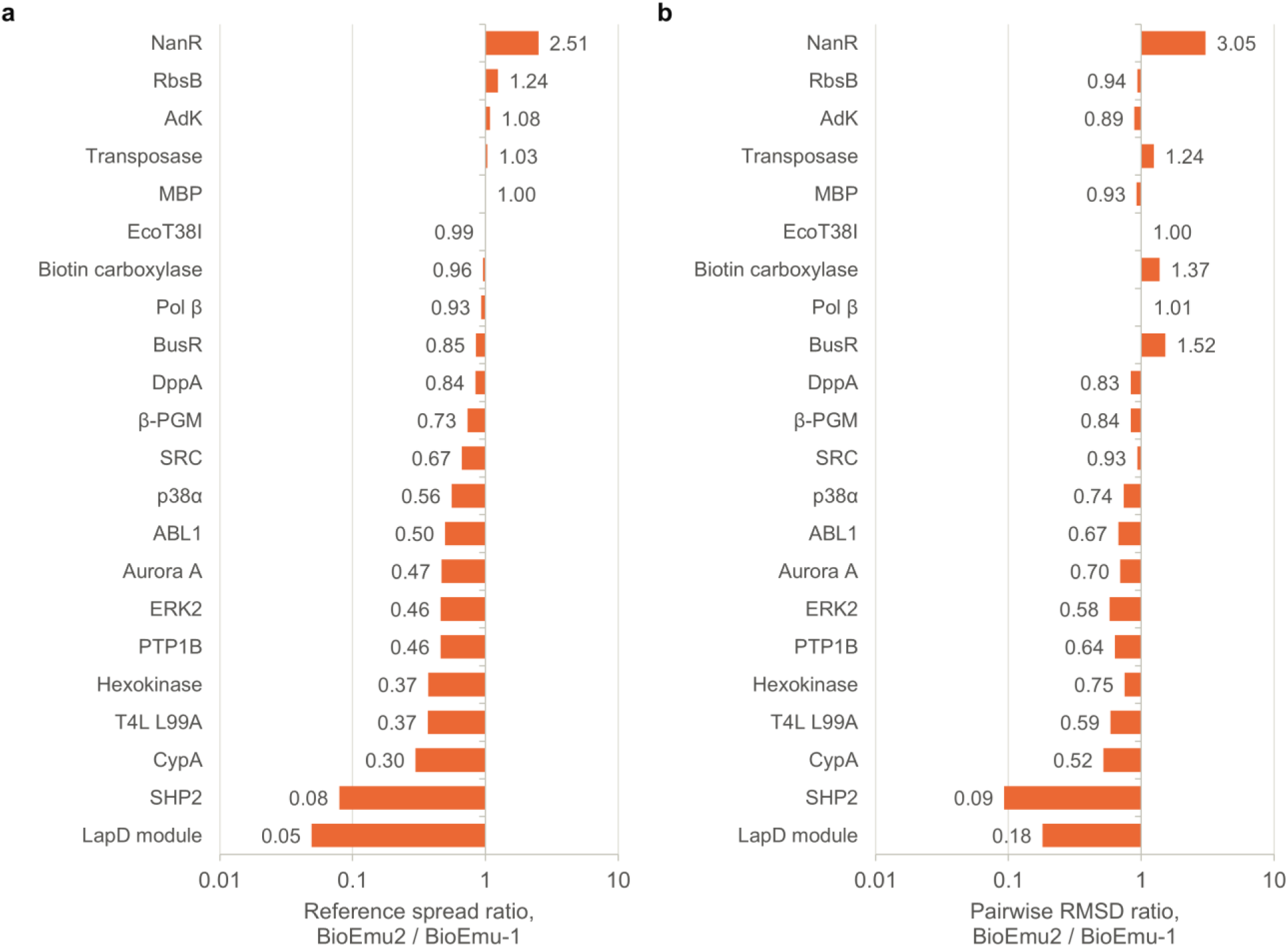
Breadth of the BioEmu2 ensembles relative to BioEmu-1. (a) Ratio of the reference spread of BioEmu2 to that of BioEmu-1 for each system. The reference spread is the median, over 20,000 random pairs of filter-passing samples, of the largest difference between the root-mean-square deviations (RMSDs) of the two samples to the same reference state. (b) Ratio of the pairwise RMSD, which is the median pairwise Cα RMSD of the scored residues among 1,000 random filter-passing samples per run.

The reference spread is measured only relative to the reference states, so we also measured the pairwise RMSD, which allows for the comparison of the samples directly with one another. The pairwise RMSD was smaller for BioEmu2 in 16 of the 22 systems (median ratio 0.83, range 0.09 to 3.05), although this difference was not statistically significant ( *p* = 0.13; Figure 5b, Table S9). BioEmu2 also formed fewer clusters at 3.0 Å in 18 of the 22 systems (median cluster count 6.5 against 85.3 per 1,000 samples). All three measures therefore showed that BioEmu2 generated structures that were more similar to one another than those of BioEmu-1, although the difference was statistically significant only for the reference spread.

A narrow ensemble does not account for every missed state. To find out why BioEmu2 missed states even where its samples were not narrower, we examined the systems in which the pairwise RMSD of BioEmu2 was at least that of BioEmu-1, and in which BioEmu-1 reached a state that BioEmu2 missed. These systems are Pol β, NanR and BusR (ratios 1.01, 3.05 and 1.52; Figure 6). In these systems, BioEmu2 spread its samples at least as widely as BioEmu-1, yet it reached neither reference state of NanR or BusR. Breadth alone cannot explain these misses, so we asked where the samples lay.

**Figure 6.**
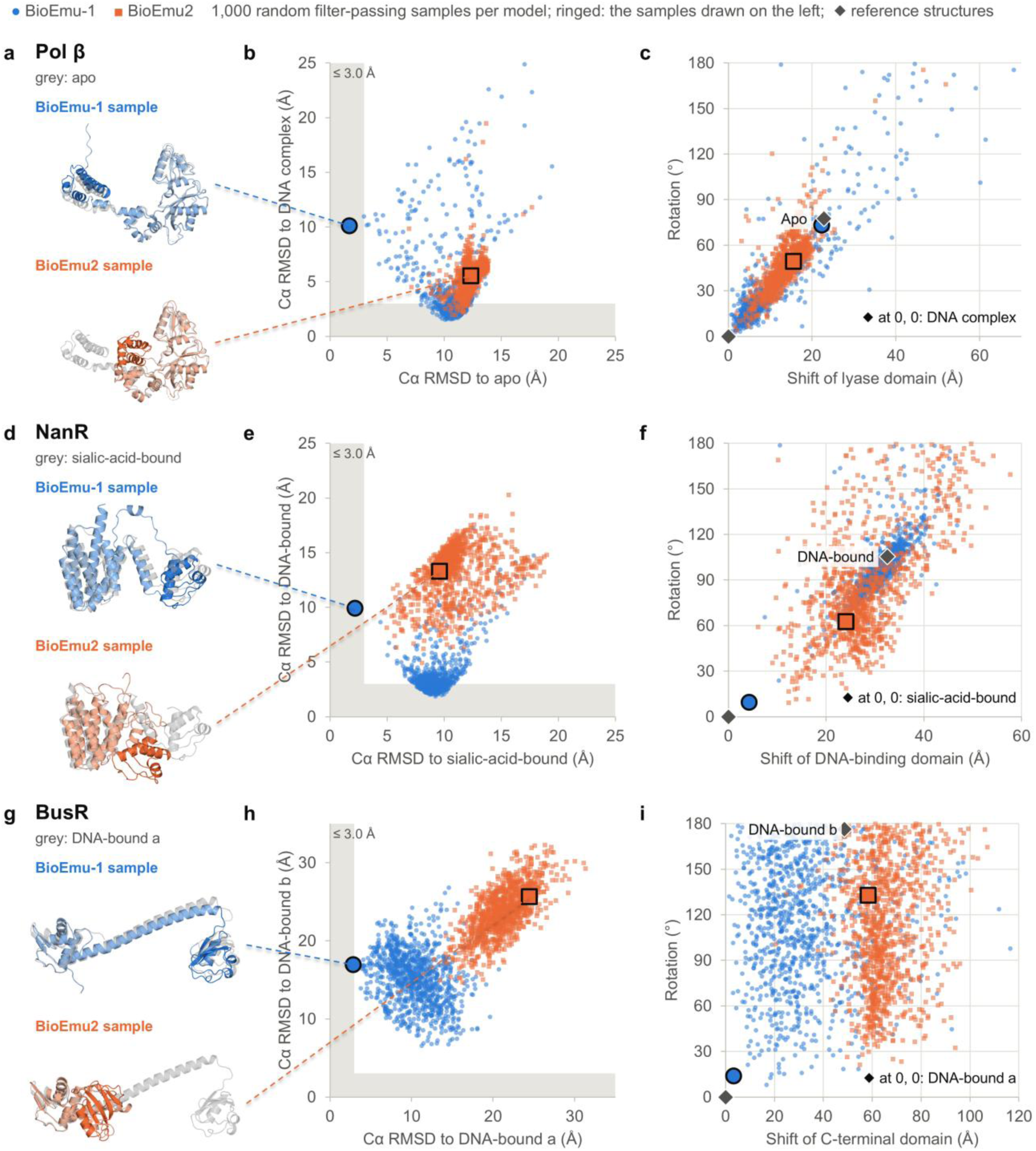
Positions of BioEmu-1 and BioEmu2 samples in Pol β, NanR and BusR. Each row shows one protein in which BioEmu2 missed a state that BioEmu-1 reached. (a, d, g) A BioEmu-1 sample within 3.0 Å of the state that BioEmu2 missed (blue) and a representative BioEmu2 sample (orange), which is the sample closest on average to the others in its group. Each is superposed on the largest domain of the reference structure of that state (transparent grey), namely Pol β (PDB ID: 1BPD), NanR (PDB ID: 6ON4) and BusR (PDB ID: 7OZ3). The moving domain is in full colour and the rest of the chain is lighter. (b, e, h) Cα RMSD of each sample to the two reference states over the scored residues. Shaded bands mark samples within 3.0 Å of a state, and dashed lines link each cartoon to its sample. (c, f, i) Shift and rotation of the moving domain from its position in one reference structure, after superposition on the largest domain. Diamonds mark the two reference structures. Each chart shows 1,000 random filter-passing samples of each model, and ringed markers are the samples drawn on the left.

We found that each domain, on its own, matched the reference structures closely in both models. For instance, in Pol β, when each domain was superposed alone, its median Cα RMSD to the nearer reference structure was 0.53 to 1.05 Å. Instead, the two models differed in how they placed the domains relative to one another. In Pol β, Bio Emu2 placed the N-terminal lyase domain a median of 14.3 Å and 48° from its position in the DNA complex, whereas BioEmu -1 placed it 6.5 Å and 20° from that position (Figure 6c).

In each of the three proteins, a few BioEmu-1 samples came within 3.0 Å of a state that none of the BioEmu2 samples reached. There were 29 such samples for apo Pol β, three for sialic-acid-bound NanR and seven for DNA-bound state a of BusR (Figure 6b, e and h). In these BioEmu-1 samples, the moving domain lies close to its position in the reference structure, whereas in a representative BioEmu2 sample it lies elsewhere (Figure 6a, d and g).

For Pol β, small-angle X-ray scattering data informed the orientation of the lyase domain. Free Pol β has a radius of gyration (Rg) of 28.7 to 29.1 Å in solution, which is consistent with the extended crystal structure of the apo enzyme (PDB ID: 1BPD).^45,46^ To compare the samples with this measurement, we averaged their Cα Rg as a root mean square, which is how a scattering experiment averages Rg over the molecules in solution. This average was 22.96 Å for BioEmu -1 and 21.82 Å for BioEmu2. These values lie about 6 to 7 Å below the measured value and close to the Rg of the DNA complex (PDB ID: 2BPG; 22.7 Å), which suggests that both models make free Pol β too compact. However, we noted that BioEmu-1 occasionally produced the extended conformation seen in solution (29 samples), whereas BioEmu2 never did (Supporting Information).

In NanR and BusR, BioEmu2 also placed the moving domain away from its reference position. In NanR, it packed the DNA-binding domain against its C-terminal domain, 26.6 Å and 77° from its position in the sialic-acid-bound structure (Figure 6f). In BusR, it brought the two terminal domains into contact in 97.4% of its samples, compared with 0.2% for BioEmu-1. However, both proteins function as oligomers in solution,^47,48^ and in their reference structures the chain also contacts another protein chain that our single-chain runs did not include. Therefore, these two misses may stem partially from modelling a single chain in isolation, rather than only from an error of the model.

We saw a related shift in ERK2, whose inactive state is apo-matched. BioEmu2 did not come closer than 3.68 Å to the inactive state or 3.72 Å to the active state. Instead, 90.4% of the BioEmu2 samples and 86.7% of the ColabFold models lay within 3.0 Å of an unphosphorylated ERK2 structure bound to a partner peptide and an ATP analogue (PDB ID: 2Y9Q). Only 4.5% of the BioEmu -1 samples lay this close to it. This structure itself lies 4.9 to 6.2 Å from both reference states. The miss of this active state may be due to the missing phosphorylation.

Taken together, these results suggest that BioEmu2 can miss states for two reasons. First, it concentrates its samples more tightly than BioEmu-1. Second, even where its samples are as broad, it can place whole domains differently from the experimental structures.

### 3.5 BioEmu2 with the Missing Partner or Phosphorylation

Several of the states that BioEmu2 missed, or reached rarely, were solved experimentally with DNA, a small signalling molecule or a phosphorylation that the sampled sequence lacked. Unlike BioEmu-1, BioEmu2 accepts binding partners and modified residues, so we were able to test directly whether these misses depended on the omitted elements. To do this, we re-ran BioEmu2 on seven proteins after adding the binding partner or post-translational modification present in each bound reference structure (Table 4).

**Table 4.**
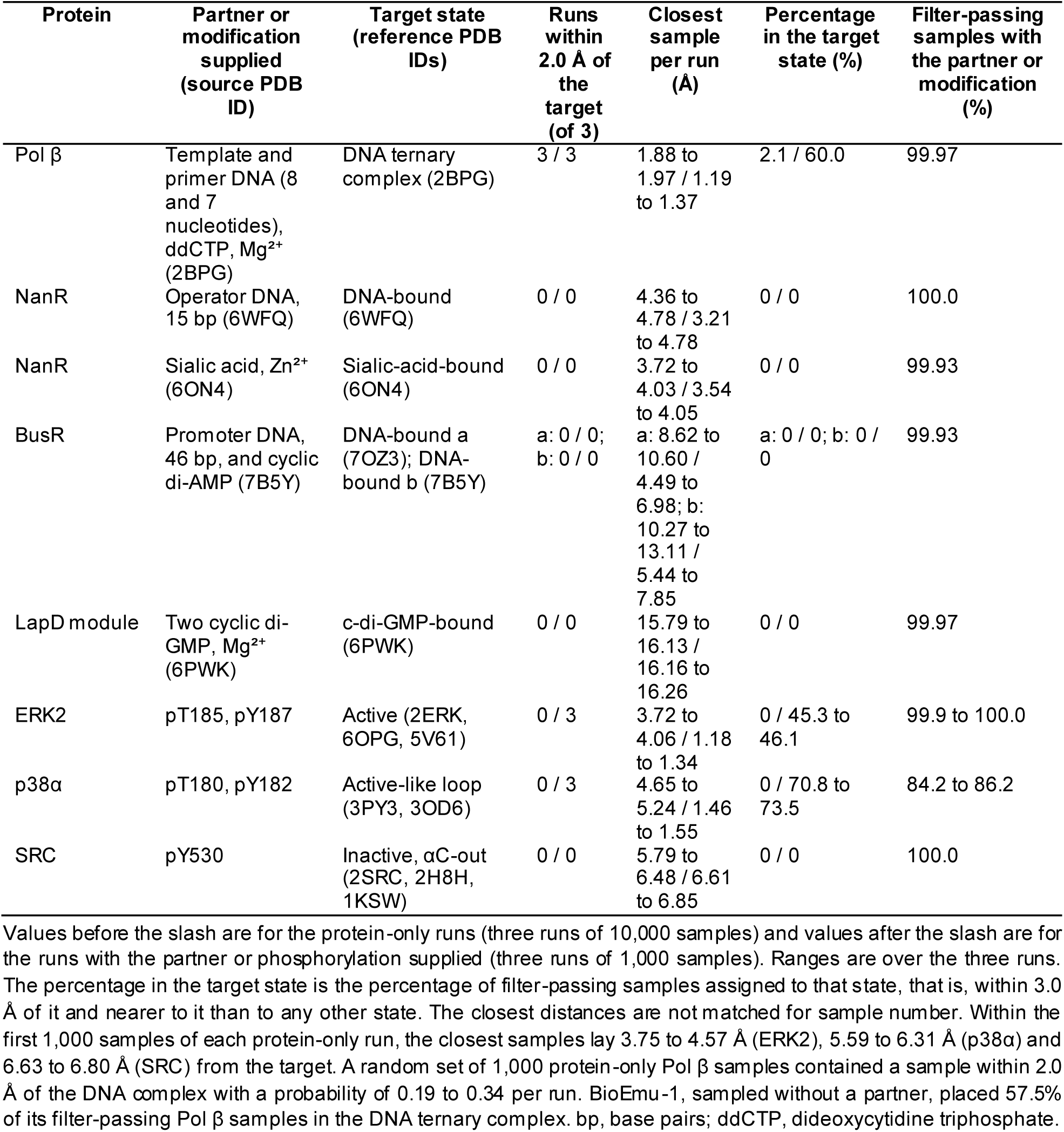
BioEmu2 runs with the missing partner or phosphorylation supplied, against the protein -only runs.

We observed that the addition of DNA shifted the Pol β samples towards the DNA-bound conformation and made it the main form. Without DNA, BioEmu2 rarely generated this arrangement. Each protein-only run contained at least one sample within 2.0 Å of the DNA complex, but only two to four of its 10,000 samples came this close, and only 2.1% of the filter-passing samples were assigned to the DNA-complex state. However, when DNA, ddCTP and Mg²⁺ were added, 53 to 56 of the 1,000 samples in each run lay within 2.0 Å of the complex, with the closest sample at 1.19 Å. Moreover, the percentage of filter-passing samples assigned to the complex rose to 60.0%, close to the 57.5% of BioEmu-1 without any partner (Figure 7a, b and g). The median Cα Rg hardly changed (21.8 Å, against 21.7 Å without the partner), which suggests that DNA changed the relative arrangement of the domains rather than their overall size.

**Figure 7.**
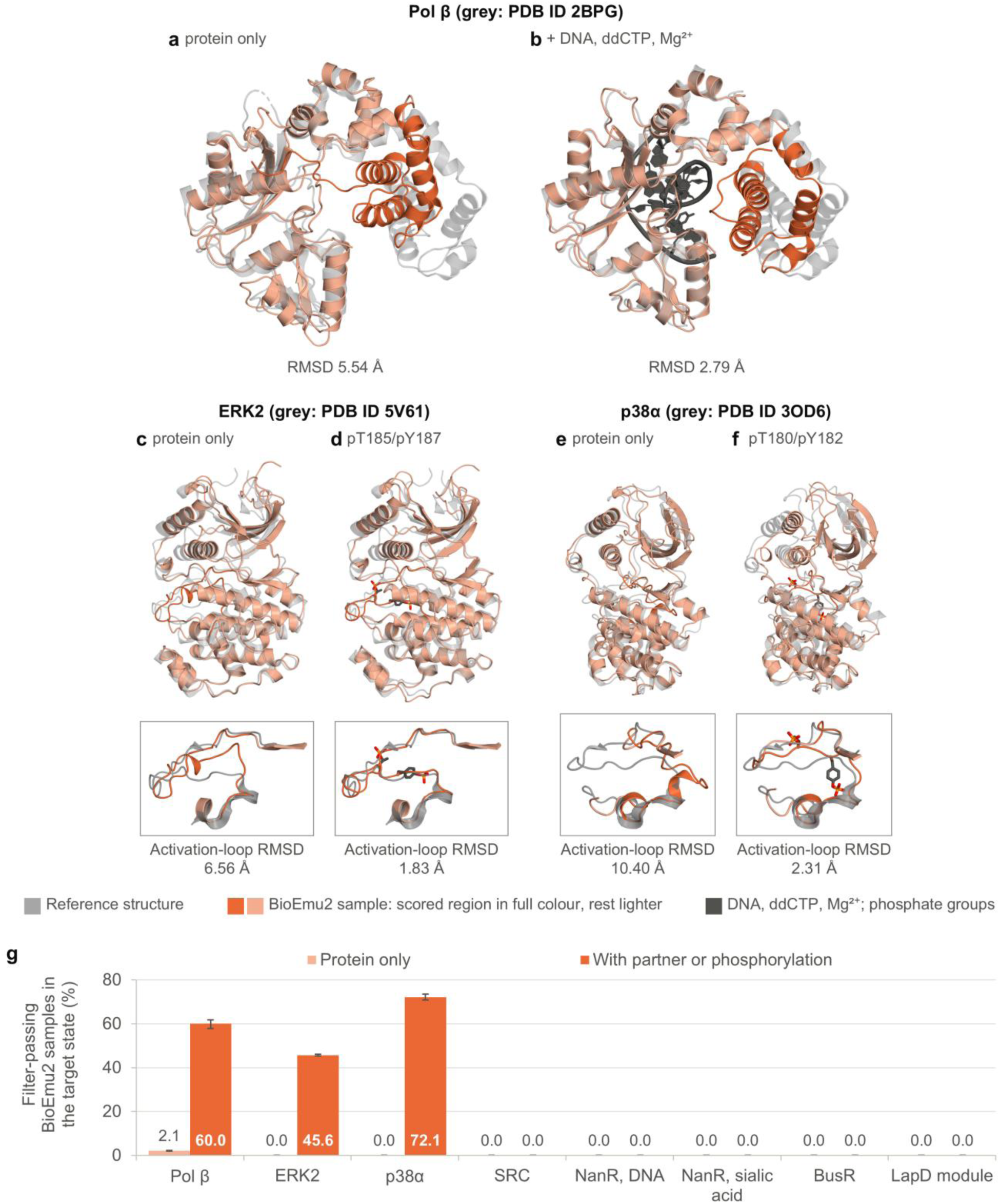
BioEmu2 with and without the missing partner or phosphorylation. (a–f) Pol β, ERK2 and p38α. Each structure panel shows the representative sample of one condition, which is the medoid of the largest cluster of filter - passing samples, and the label below it gives the Cα RMSD of its scored residues to the reference shown. (a, b) Pol β without DNA (a) and with the template and primer DNA, dideoxycytidine triphosphate (ddCTP) and Mg² ⁺ of the ternary complex (PDB ID: 2BPG) (b), superposed on the polymerase core of that complex (transparent grey). The lyase domain (residues 1 to 90) is shown in full colour. After superposition on domain D1, the lyase domain of the representative sample lies 15.6 Å from its position in the ternary complex without DNA and 7.2 Å with DNA. (c –f) ERK2 (c, d) and p38α (e, f) without (c, e) and with (d, f) phosphorylation of the two activation -loop sites, superposed on the C-terminal lobe of an active ERK2 structure (PDB ID: 5V61) or of an active -like structure of an unphosphorylated Y323T variant of p38α (PDB ID: 3OD6). Below each kinase structure, the activation loop is enlarged in the same orientation. The N-terminal lobe of the p38α sample lies closer to the reference in (e) than in (f) (residues 4 to 110, 2.40 and 4.80 Å Cα RMSD after superposition on the C-terminal lobe), but its activation loop does not. (g) Percentage of filter-passing samples in the target state without and with the partner or phosphorylation (mean over three runs, with bars showing the range over runs).

Contrastingly, in NanR, BusR and the LapD module, BioEmu2 still could not generate any of the five partner-bound states that it had missed, even when the binding partners were included. None of the runs contained a filter-passing sample within 3.0 Å of these states (Table 4). For BusR, DNA and cyclic di-AMP moved a minority of samples closer to both DNA-bound states, but the median distances to the two states did not decrease (Supporting Information). In the bound structures of all three proteins, however, the reference chain also contacts another protein chain that these ru ns did not include, so these negative results remain inconclusive.

On the other hand, phosphorylation of the activation loop let BioEmu2 reach the active states of both ERK2 and p38α in all three runs of 1,000 samples. Without phosphorylation, neither state had been reached in any run, and none of the samples had been assigned to either state, even after 10,000 samples per run. With phosphorylation, the active state held 45.3% to 46.1% of the filter -passing ERK2 samples, and the active-like state 70.8% to 73.5% of the filter-passing p38α samples. The representative structures also moved from a distant loop conformation to one close to the reference (Figure 7c–g).

Phosphorylation also changed how the other states were populated. In p38α, it removed the inactive loop state, which had held 33.0% to 37.0% of the first 1,000 samples of each protein -only run. In ERK2, it lowered the percentage of samples near the peptide-bound ERK2 structure from 90.4% to 1.2%, although the inactive state itself stayed unreached (closest samples 3.29 to 3.76 Å).

However, we found a structural defect in the phosphorylated p38α samples. In 14.7% of them, the chain was broken on either side of pY182, so that they failed the backbone filter, whereas only one sample in 3,000 failed in the unmodified p38α runs and in the phosphorylated ERK2 runs. The HIGH_QUALITY check of BioEmu2 still passed 99.7% to 99.9% of these samples per run. The failing samples occupied the active-like state at nearly the same rate as the filter-passing ones (71% and 72%), so filtering does not explain the shift towards the active-like state.

However, phosphorylation of the SRC tail tyrosine did not draw samples closer to the inactive state. The closest distance to it stayed at 6.61 to 6.85 Å, against 6.63 to 6.80 Å for the first 1,000 unmodified samples, and the active state still held 43.3% to 45.6% of samples, against 42.5% to 45.3% (Supporting Information).

Taken together, these results suggest that some of the states missed by BioEmu2 reflect what the model was given rather than what it can generate. Supplying a known partner or modification may therefore be worth trying when a bound or modified state is of interest. However, all six partner-bound structures were released before the Boltz-1 training cut-off, so the positive results may reflect recall of these structures, or of the phosphorylated kinase structures, rather than a prediction of how the partner or phosphorylation changes the structure.

### 3.6 State Preferences of the Two Models

So far, our analyses have asked whether each model reaches the experimentally solved states of each protein. However, reaching a state shows only that a model can generate it, not how often it does so. Therefore, we further examined whether each model assigns these states the correct populations.

First, we asked whether the two models give the same populations as each other. Each run used its own random seed and a finite number of samples, so the population of a class can differ between runs of one model by chance alone. Hence, before comparing the models, we checked that their differences exceed this variation. Across the 71 classes of the 22 systems (49 states and one unassigned class per system), the median SD of a class population over the three runs of one model was 0.17 pp, and the largest was 0.82 pp. By contrast, the two models differed by more than 1 pp in 54 of the 71 classes, and each of these differences was at least 3.6 times the SD of either model (Figure S1). Hence, differences of this magnitude reflect the models themselves rather than the variation between runs.

To compare the two models within each system, we identified the class in which their populations differed the most (Figure 8a). We summarised each system by this single class because the systems have different states, so none of the states can be compared across all systems. A single class is enough because the class populations of each model sum to 100%, so any gain in one class must be balanced by losses in the others. The largest difference therefore gives the minimum percentage of samples that the two models place in different classes. A positive difference means that BioEmu2 placed more samples in the class than BioEmu-1. For example, 96.7% of the filter-passing β-PGM samples from BioEmu2 were open, compared with only 10.1% of those from BioEmu-1, a difference of +87 pp. At the class threshold of each system, the largest difference was between 29 and 87 pp, whether positive or negative, in 13 of the 21 systems shown, and between 0 and 26 pp in the remaining eight. CypA is not shown because its two backbone states are only 0.11 Å apart.

**Figure 8.**
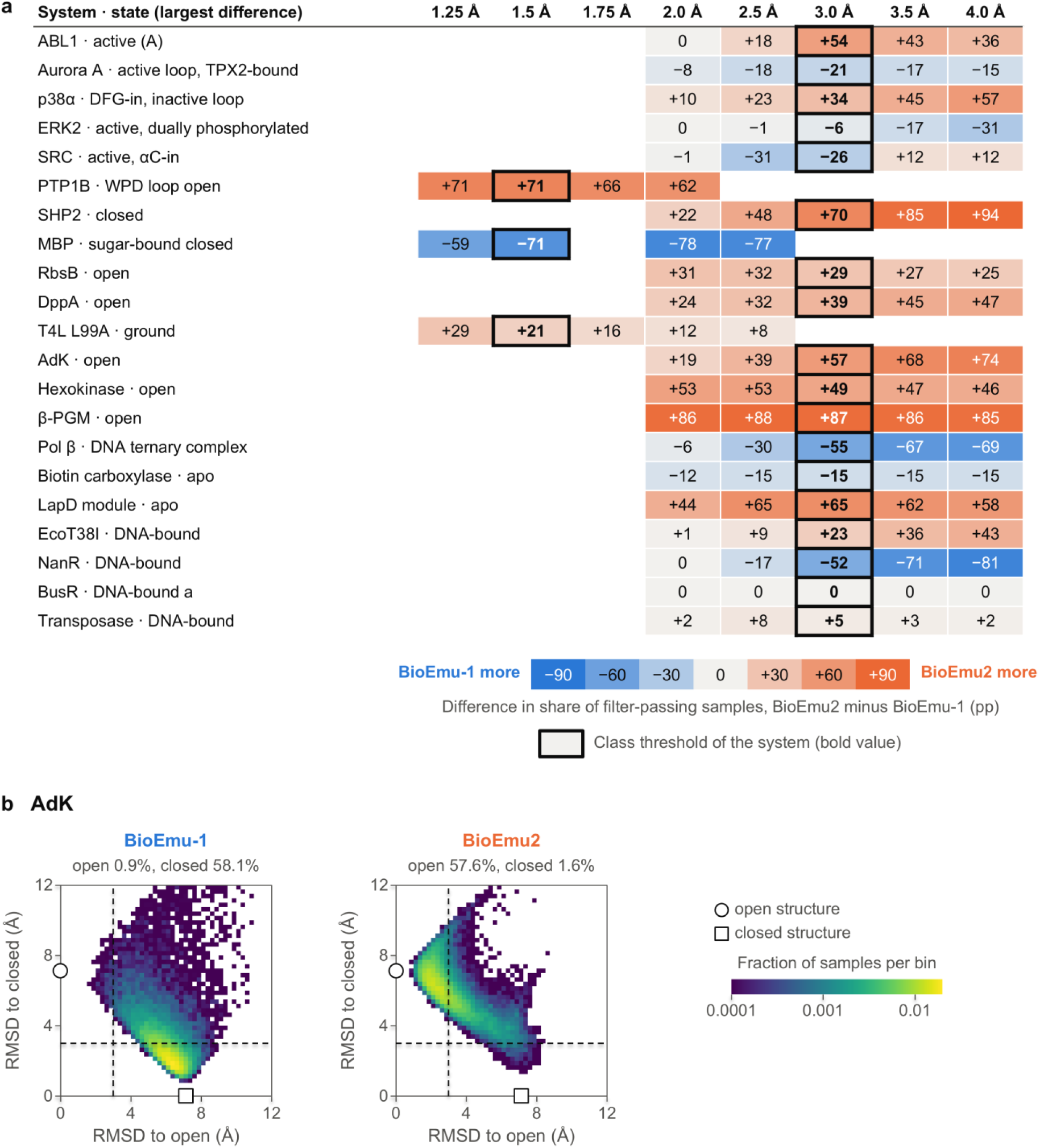
State preferences of BioEmu2 relative to BioEmu-1. (a) For each system, the class with the largest population difference between the two models at the class threshold of that system, shown as BioEmu2 minus BioEmu-1 in percentage points of filter-passing samples at each class threshold tested. Orange cells mean that BioEmu2 places more samples in the class, and blue cells mean that BioEmu -1 does. Bold, outlined cells mark the class threshold of each system, and blank cells were not tested. CypA is not shown, because its two backbone states are 0.11 Å apart. (b) Cα RMSD of AdK samples to the open structure (PDB ID: 4AKE) against Cα RMSD to the closed structure with Ap5A (PDB ID: 1AKE) for the filter-passing samples of BioEmu-1 and BioEmu2 (three pooled runs of 10,000 samples), in 0.25 Å bins on a logarithmic colour scale. Dashed lines mark 3.0 Å.

To find out which states each model favoured, we first examined the five two-state proteins whose closed forms were solved with a bound substrate, analogue or inhibitor. These proteins are AdK, RbsB, DppA, hexokinase and β-PGM. In all five, BioEmu2 placed 29 to 87 pp more filter-passing samples in the open state than Bio Emu-1 (Figure 8a). For example, AdK was 57.6% open with BioEmu2, compared with 0.9% open with BioEmu-1 (Figure 8b). By contrast, ColabFold favoured the closed state in all five proteins (60.8% to 100% of its filter-passing models closed and 0% to 29.0% open). In each protein, BioEmu2 placed more samples in the open state than either of the other two methods. The open state was apo-matched in three of these proteins (AdK, RbsB and DppA), whereas the open structures of hexokinase and β-PGM carry a bound nucleotide analogue and BeF₃⁻ with Mg²⁺, respectively.

We saw the same preference for the open form in PTP1B and MBP. BioEmu2 kept the PTP1B WPD loop mainly open (2.3% of assigned samples closed) and MBP mainly open (62.1% open and 1.22% in the sugar-bound form). Conversely, BioEmu-1 favoured the closed WPD loop (71.8%) and the sugar-bound form of MBP (72.4%), as did ColabFold (96.0% and 54.4%).

Likewise, in p38α, BioEmu2 favoured the apo-matched state. It placed 34.3% of its filter-passing samples in the DFG-in inactive loop and none in the active-like loop, which was solved with phosphorylation or a mutation. BioEmu-1 and ColabFold, in contrast, placed 14.6% and 85.4% of their samples or models in the active-like loop and almost none in the inactive loop. There were two exceptions to this preference of BioEmu2. Relative to BioEmu-1, it placed fewer samples in the apo state of biotin carboxylase (0.9% against 16.1%) and more in the DNA-bound state of EcoT38I (26.2% against 2.9%).

We then checked whether two choices in our scoring could have created these differences. First, the sign of the largest difference stayed the same at every class threshold that we tested in 19 of the 21 systems (Figure 8a). The exceptions were ABL1, where the difference was negative only at 2.0 Å and then by only 0.02 pp, and SRC, where it changed from −26 pp at 3.0 Å to +12 pp at 3.5 and 4.0 Å. Second, removing the samples that fail the backbone filter did not create these differences. For example, BioEmu-1 failed 35.08% of its SHP2 samples. By the N-SH2/PTP interface distance, 94.6% of these failed samples were open, compared with 98.5% of the samples that passed.

BioEmu2 favoured the open form of proteins whose closed form was solved with a ligand, so we investigated whether such a closed form is stable once the ligand is removed. To answer this, we examined AdK, the only one of the five proteins that we also simulated with all-atom MD. Replicas started from the closed structure, simulated without its bound ligand Ap5A, moved to the open state after 12.4, 12.6 and 48.2 ns, whereas replicas started from the open state stayed open (Table S16). This may indicate that the closed form is not stable without its ligand on this time scale, which would be consistent with the preference of BioEmu2 for the open state. The BioEmu2 developers also reported that adding ATP and AMP to the input shifted the BioEmu2 samples of AdK towards the closed state. In their propagation runs, AdK closed in about 40% of 50 trajectories with these ligands.^12^

One possible explanation for the closed preference of BioEmu-1 is that the PDB holds many closed, ligand-bound structures of such proteins. Although BioEmu-1 was not trained on PDB entries, the AlphaFold2 sequence representation that it uses was trained on them, so such a bias could reach it indirectly. Zha et al. also reported that BioEmu-1 samples were skewed towards ligand-bound states,^13^ but we did not test this explanation.

Taken together, these results suggest that the two generations give clearly different populations for the same protein, with BioEmu2 tending to favour the open form of proteins whose closed form was solved with a ligand.

### 3.7 Populations Compared with Solution Measurements

Next, to investigate whether BioEmu-1 and BioEmu2 can reproduce biologically relevant populations measured in solution, we used the three systems other than ABL1 whose populations have been measured with an error bar, namely PTP1B, SHP2 and Aurora A (Table 5). However, it is noteworthy that the sampling frequencies of BioEmu2 are not calibrated Boltzmann probabilities.^11,12^ Instead, the comparison evaluates the extent to which BioEmu2 samples can be interpreted as populations. We first asked how widely each model spreads its samples along the variable that separates the states of each protein, such as the distance between two residues.

**Table 5.** Populations of the measured states outside ABL1 for each method, against the measured values.

| System · state | Measured (tolerance range) | BioEmu-1 | BioEmu2 | ColabFold | All-atom MD |
|---|---|---|---|---|---|
| PTP1B · WPD loop closed, backbone class at 1.5 Å | 2.5% (0 to 5.3%) | 71.8% (0/3) | 2.3% (3/3) | 96.0% (0/1) | By start state<sup>a</sup> |
| SHP2 · open, interface distance above 10 Å | 10.4% (8.2 to 12.6%) | 98.5% (0/3) | 0%<sup>b</sup> (0/3) | 93.7% (0/1) | Not run |
| Aurora A · activation loop closed, K224–S283 Cα distance below 22 Å | 52.4% (47.8 to 56.9%) | 1.20% (0/3) | 0.05% (0/3) | 0.20% (0/1) | Not run |
| MBP · sugar-bound closed (upper bound) | Below 1% | 72.4% | 1.22% (1.04% to 1.42%) | 54.4% | Not run |
| MBP · partially closed (no reported uncertainty) | About 5% | 0.003% (1 sample) | 0% | 0% | Not run |
| T4L L99A · excited, backbone class (no reported uncertainty) | About 3% | 0.02% (0.010% to 0.050%) | 0% (0 of 30,000) | 0% | By start state<sup>a</sup> |
| T4L L99A · Phe114 χ1 trans | About 3% | 0.3% (0.1% to 0.4%)<sup>c</sup> | 0% | 0% | By start state<sup>a</sup> |
| CypA · Phe113 minor rotamer (upper bound) | Below 10% | 3.6% (3.2% to 4.2%)<sup>c</sup> | 7.9% (7.5% to 8.1%) | 66.5% | By start state<sup>a</sup> |

Values are percentages of filter-passing samples (mean over runs, with the range over runs where given). For the three populations measured with an uncertainty (PTP1B, SHP2 and Aurora A), the values are renormalised over the two measured states, and the number of runs within the tolerance (twice the reported uncertainty) is given in brackets. ColabFold has one set of 500 models. Populations before renormalisation, unassigned fractions and comparisons with other definitions of the states are in Tables S10 to S12. <sup>a</sup> The occupancy of each state depended on the start state and varied widely between replicas (Table S15), so a value pooled over replicas would reflect the chosen start states rather than an equilibrium population. <sup>b</sup> 0 of 29,990 filter-passing samples. The exact one-sided 95% upper bound on the sampling frequency is 0.010%. <sup>c</sup> BioEmu-1 outputs backbone and Cβ atoms only, so this value describes the rotamer that HPacker chose for 1,000 random filter-passing BioEmu-1 backbones per run. When BioEmu2 samples were rebuilt in the same way, HPacker kept the Phe113 rotamer of BioEmu2 in 99.5% of samples and its Phe114 rotamer in all samples.

We found that the width of the middle 90% of the samples was smaller for BioEmu2 than for BioEmu-1 in all four proteins with measured populations, at 1.4 against 49.0 Å in SHP2, 6.1 against 13.5 Å in Aurora A, 3.3 against 8.2 Å in PTP1B and 0.8 against 1.6 Å in ABL1 (Figure 9a–d). In SHP2, PTP1B and ABL1, the BioEmu2 samples clustered in the state adopted by most molecules in solution without a ligand. In Aurora A, BioEmu2 sampled the extended loop, although the loop is closed in about half of the measured molecules.

**Figure 9.**
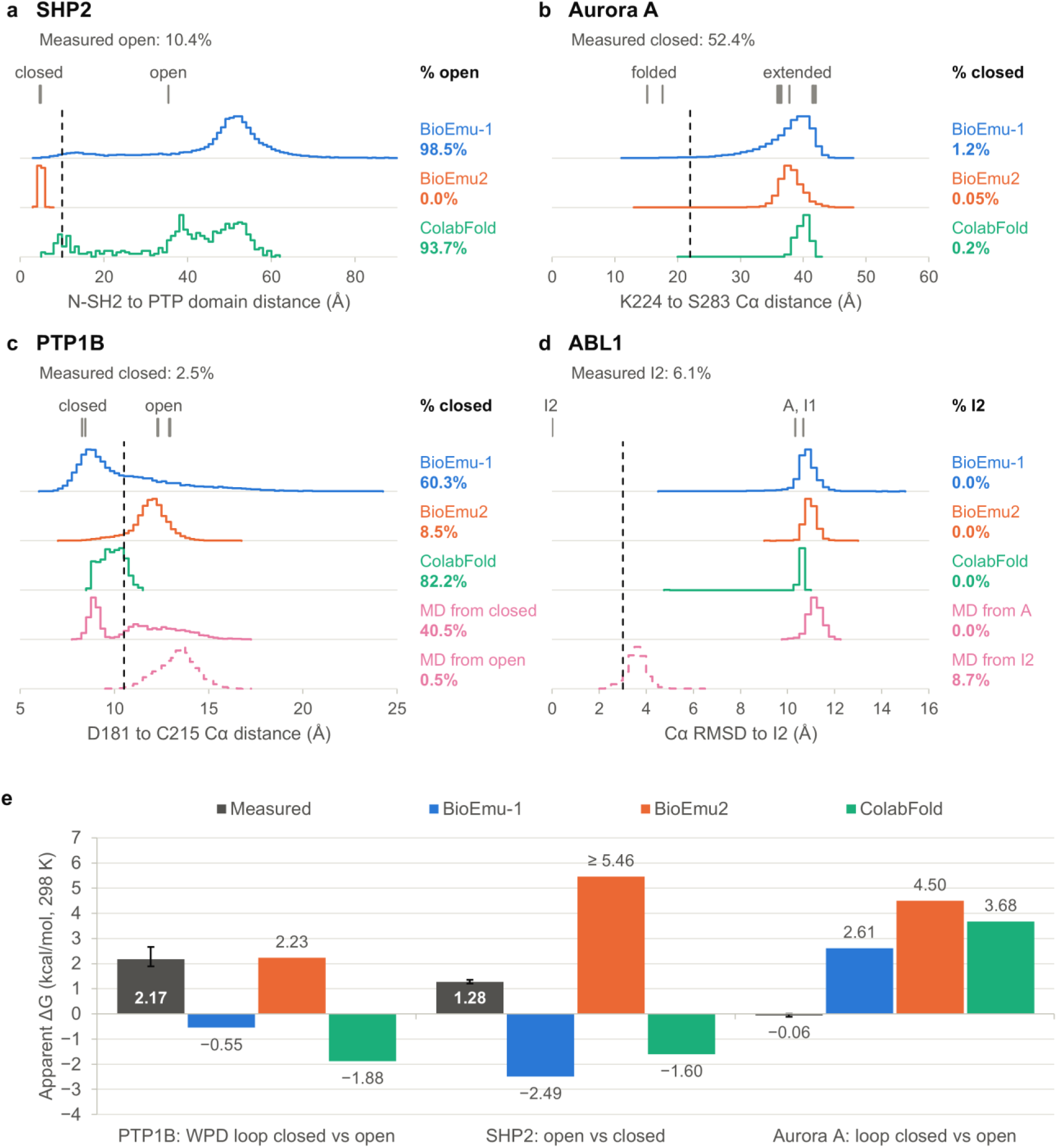
Sampled distributions and populations against measured values. (a–d) Distributions of the samples along the variable that defines the states of SHP2, Aurora A, PTP1B and ABL1, the four systems with a measured population, for BioEmu-1, BioEmu2, ColabFold and, where it was run, all-atom MD (each start state separately), each scaled to its own maximum. The panels show filter-passing samples (three pooled runs of 10,000 samples for each BioEmu model, the 500 ColabFold models and MD frames from 10 to 250 ns), and the MD percentages reflect the start state and are not populations. The dashed line marks the cut-off of the named state, the percentages give the percentage of samples on that side of it, and grey ticks mark the reference structures. (a) SHP2, shortest Cα distance between the N-SH2 and PTP domains in 1 Å bins, with the open state above 10 Å. The tick labelled open is the structure of the E76K mutant (PDB ID: 6CRF). (b) Aurora A, Cα distance between K224 and S283 in 1 Å bins, with the closed state below 22 Å. (c) PTP1B, Cα distance between D181 and C215 in 0.25 Å bins, with the closed state below 10.5 Å, the midpoint between the mean closed and mean open reference distances. This is the secondary definition, whereas Table 5 uses the backbone classes. (d) ABL1, Cα RMSD to the I2 structure in 0.25 Å bins, with the 3.0 Å class threshold. The I2 class used for the ABL1 populations is assigned from backbone rules rather than from this RMSD, so its percentages differ. (e) Apparent free-energy differences between the two measured states of PTP1B, SHP2 and Aurora A, ΔG = −RT ln(p₁/p₂) at 298.15 K, where p₁ and p₂ are the populations of the two states, from the measured populations (error bars, ±1 reported uncertainty) and from the mean population over runs. The SHP2 bar for BioEmu2 is a lower bound.

Next, we examined PTP1B, the only system in which a BioEmu model fell within the tolerance. The WPD loop of PTP1B carries the catalytic Asp181 and closes over the active site. In apo PTP1B, NMR relaxation dispersion indicates that the loop is closed in 2.5 ± 1.4% of molecules, ^24^ which gives a tolerance range of 0% to 5.3%. With the loop counted as closed within 1.5 Å Cα RMSD of a closed crystal structure, BioEmu2 placed 2.05% to 2.50% of its assigned samples in the closed state, so all three runs fell within the tolerance. These closed samples matched the reference structures closely (186 to 226 samples per run within 1.5 Å; closest 0.60 to 0.81 Å). Nevertheless, this agreement has to be treated with caution. One reason is that the tolerance range includes zero, so an ensemble without any closed samples would also satisfy it. Another reason is that the measured uncertainty was derived from fits at only two residues.

Moreover, we found that the agreement depends on how the closed state is defined. A class threshold of 1.25 Å lowered the closed percentage of BioEmu2 to 0.90% to 1.17%, and one of 2.0 Å raised it to 5.21% to 5.91%, which placed one run outside the tolerance. We also counted closed samples on a second measure from the literature, the Cα distance between Asp181 and Cys215 (Figure 9c, Table S11, Figure S2).^49^ On this measure, BioEmu2 stayed within the tolerance in all three runs for cut-offs up to 9.87 Å, the threshold used by Yeh et al., but not from 10.075 Å upwards. At our secondary boundary of 10.5 Å, it placed 8.46% of its samples in the closed state.

BioEmu2 therefore gives a closed population near the measured value under some reasonable definitions of the closed state but not under others. BioEmu-1 and ColabFold, in contrast, placed far more samples in the closed state than the measurement allows (71.8% and 96.0%), and BioEmu-1 fell outside the tolerance at every distance cut-off.

Moving on, we examined SHP2, for which none of the methods reproduced the measured population. Wild-type SHP2 is 10.4 ± 1.1% open in solution,^25^ but none of the methods matched this value at any cut-off of the interface distance from 8 to 35 Å (Figure 9a, Table S12). BioEmu2 kept the N-SH2 domain against the PTP domain in every filter-passing sample (median 4.98 Å; maximum 7.31 Å), so its rate of open samples is below 0.010% with 95% confidence. In contrast, BioEmu -1 placed 83.6% to 99.5% of its filter-passing samples in the open state across this range of cut-offs, and ColabFold 75.5% to 98.6% of its models. However, in 88.2% of the open BioEmu-1 samples, the N-SH2 domain packed against the C-terminal SH2 domain, a contact that none of the crystal structures shows.

Finally, we examined Aurora A, in which all three methods placed the activation loop in the open state in almost every sample (Figure 9b). Single-molecule fluorescence measurements of unphosphorylated, ligand-free Aurora A found the loop closed in 52.4 ± 2.3% of molecules. ^26^ In contrast, BioEmu-1 placed 1.14% to 1.28% of its samples in the closed state, BioEmu2 0.02% to 0.09% and ColabFold 0.20%, so all three lay outside the tolerance of ±4.5 pp. This result did not depend on the cut-off. With any cut-off from 18 to 35.8 Å, BioEmu2 placed at most 8.7% of its samples in the closed state, BioEmu-1 at most 27.5% and ColabFold at most 0.2% (Supporting Information). However, the measured protein carried two dye-labelled cysteines and was attached to a surface, which the models do not represent.

To compare the size of these errors across the three proteins on one scale, we converted each population into an apparent free-energy difference between the two states (Figure 9e). On this scale, an error of 1.36 kcal/mol at 298 K corresponds to a 10-fold error in the ratio between the two states. BioEmu2 differed from the measured value for PTP1B by only 0.05 kcal/mol. However, it made the closed state of SHP2 too stable by at least 4.18 kcal/mol and the open loop of Aurora A too stable by 4.56 kcal/mol. BioEmu-1 inverted the balance of PTP1B and SHP2 (errors of −2.72 and −3.76 kcal/mol) and made the open Aurora A loop too stable by 2.67 kcal/mol. ColabFold behaved like BioEmu-1, with errors of −4.05, −2.87 and +3.74 kcal/mol.

The errors therefore did not follow a simple pattern. The models neither placed roughly equal percentages of samples in the two states nor erred in the same direction in every protein. Instead, BioEmu2 placed nearly all of its samples in a single state. This agrees with the caution in the BioEmu2 model card.^11^

For MBP, CypA and T4L L99A, the measurements give a value without an error bar or only an upper limit, so we compared these systems without a tolerance (Table 5). In MBP, paramagnetic relaxation enhancement data rule out more than 1% of the sugar-bound closed form in the apo protein and show a partially closed form at about 5%.^29^ Bio Emu2 came closest to this limit, with 1.22% of its samples in the sugar-bound form, against 72.4% for BioEmu-1 and 54.4% for ColabFold. Every BioEmu2 run, however, lay slightly above the 1% limit, and none of the methods populated the partially closed form.

In CypA, BioEmu2 placed the Phe113 side chain in the minor rotamer in 7.9% of its samples, below the upper limit of 10%,^30^ but only 0.25% to 0.38% of its samples also adopted the Ser99 rotamer of the crystal minor state. In T4L L99A, whose excited state is populated at about 3%, ^27^ BioEmu2 placed none of its 30,000 samples in that state, and BioEmu-1 seven of its 29,756 filter-passing samples.

Taken together, only one of the six comparisons with a measured error bar fell within the tolerance, and that agreement depends on how the closed state of PTP1B is defined. These results suggest that the sampling frequencies of either model should not be read as populations or free energies. They also suggest that the narrow ensembles of BioEmu2 can place almost all of its samples in one state even when solution measurements show a second state.

### 3.8 ABL1 Populations and Their Response to Perturbation

Lastly, we assessed whether the populations of ABL1 change in the same direction as the measured populations when the protein is perturbed by mutations, phosphorylation or ligands. ABL1 is the only system with measured populations for three states, an active state and two inactive states (I1 and I2), and for their changes on mutation, phosphorylation and ligand binding. ^2^ The three states differ mainly in the conformation of the DFG motif and the activation loop, so we assigned them from backbone rules rather than by RMSD (Figure 10a).

**Figure 10.**
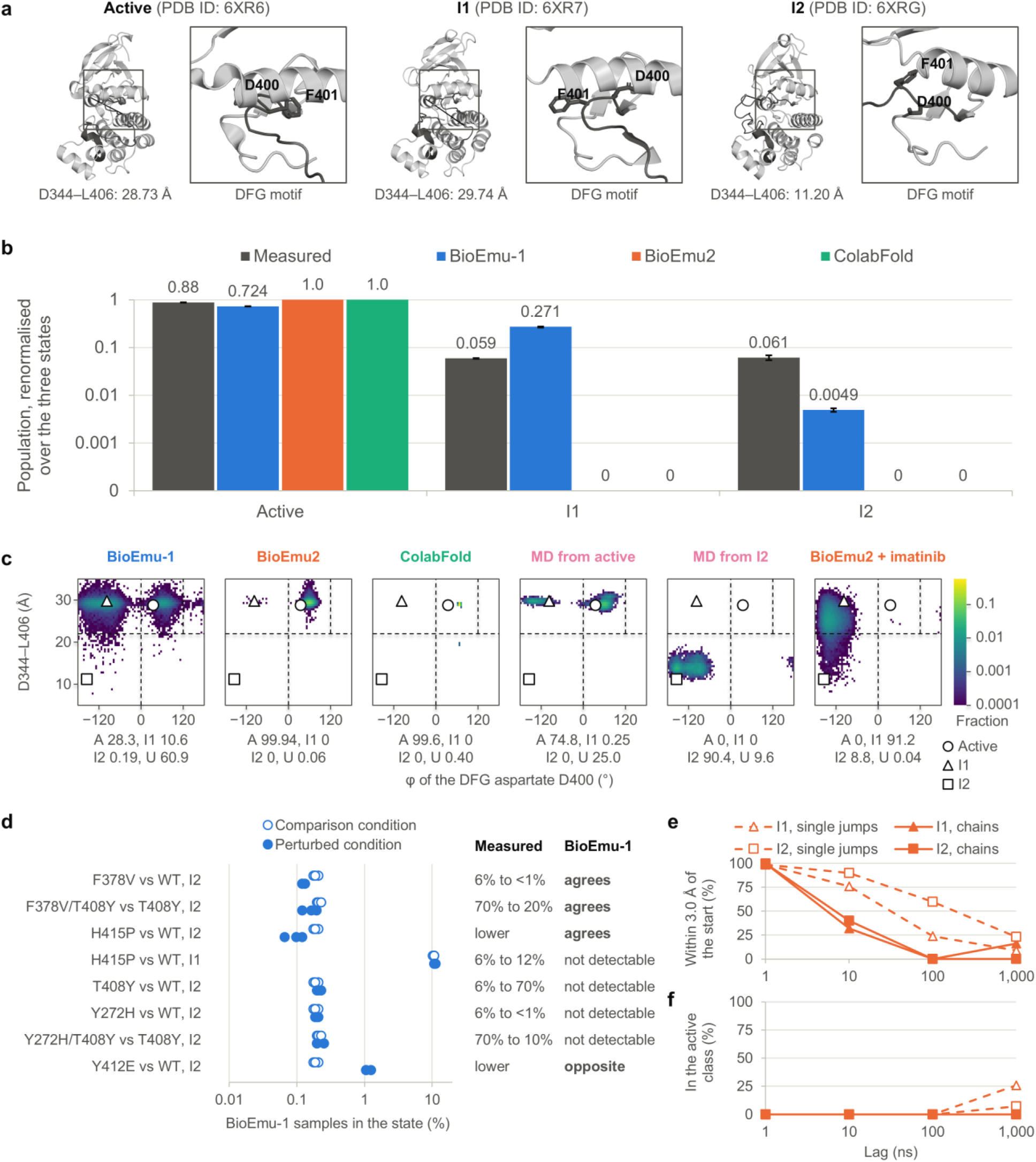
ABL1 states, populations, perturbations and BioEmu2 propagation. (a) NMR structures of the active state (PDB ID: 6XR6), I1 (PDB ID: 6XR7) and I2 (PDB ID: 6XRG). Each image shows the medoid of the models that the class rules assign to that state, superposed on the core of the C-terminal lobe. The DFG motif and activation loop are dark grey, the D344–L406 Cα distance is dashed and a close-up shows D400 and F401. (b) Wild-type populations of the active, I1 and I2 states, renormalised over the three states, for the NMR measurement (error bars, ±1 reported uncertainty) and for each method (mean over runs, with bars showing the range over runs, and ColabFold at the 32:64 alignment depth). (c) Distribution of filter-passing samples for each method by the φ angle of D400 and the D344–L406 Cα distance, as the fraction of samples per 5° by 0.5 Å bin on a logarithmic colour scale. Dashed lines mark the class cut-offs, with the loop closed below 22 Å and DFG in for φ from 0° to 120° and out below 0°. A sample is assigned only if the A399 ψ angle is in the β region, and the numbers give the percentages in the active (A), I1, I2 and unassigned (U) classes. The maps show three pooled runs of 10,000 samples for BioEmu -1, BioEmu2 and BioEmu2 with imatinib, 500 ColabFold models at the 32:64 alignment depth, and all-atom MD frames from 10 to 250 ns of the three replicas started from each state. The MD percentages therefore reflect the start state and are not populations. (d) Percentage of filter-passing BioEmu-1 samples in the named state for the three runs of the comparison (open circles) and perturbed (filled circles) conditions, with the measured change. For Y412E, the measured change is that of Y412 phosphorylation. (e, f) BioEmu2 propagation from five NMR models each of I1 and I2, given as the percentage of structures within 3.0 Å of the start structure (e) and in the active class (f), for single jumps of the given lag (32 endpoints per start, open symbols and dashed lines) and for chains of 1 -ns steps (filled symbols and solid lines).

We first compared the wild-type populations with the NMR values of 88.0% active, 5.9% I1 and 6.1% I2 (Figure 10b, Table S13). We found that none of the methods reproduced this balance, and that none of the runs lay within the tolerance of any state. BioEmu2 placed 99.94% of its filter -passing samples in the active class and none of 29,997 in I1 or I2, so each inactive population was at most 0.010% at 95% confidence. By contrast, BioEmu-1 left 60.9% of its filter-passing samples unassigned, and over the three states its populations were 0.724, 0.271 and 0.0049, against the measured 0.880, 0.059 and 0.061. ColabFold placed 96.2% to 99.8% of its models in the active class at the three alignment depths. As apparent free energies relative to the active state, the measured values are 1.60 kcal/mol for I1 and 1.58 kcal/mol for I2, against 0.58 and 2.96 kcal/mol for BioEmu -1 and at least 5.46 kcal/mol for BioEmu2.

We next asked which ABL1 structures the two models reached. Both came within 3.0 Å of the I1 reference in every run (closest 2.12 Å for BioEmu-1 and 2.15 Å for BioEmu2), and neither came within 3.0 Å of I2, whose activation loop is closed (closest 4.51 and 9.17 Å, Figure 9d). The two models therefore reached similar structures. However, BioEmu2 sampled backbones close to I1 without the DFG flip that defines the I1 class, and it never closed the activation loop (Figure 10c).

Meanwhile, in all-atom MD, the active state and I2 did not interconvert in 250 ns. Replicas started from I2 stayed in the I2 class (77.4% to 99.0% of frames), although most of their frames lay more than 3.0 Å from the I2 structure. Two of the three replicas started from the active state stayed active (98.6% and 99.9% of frames). In the third, the DFG backbone changed after 71.9 ns, but the aspartate never entered the back pocket and the activation loop stayed open. Only 25.8% of its frames were therefore active, and most of the rest were unassigned rather than I1.

Next, to test whether the models shift the populations in the same direction as the measurements, we compared each perturbed condition with the condition that it was measured against (Figure 10d, Table S14). BioEmu2 could not respond to most perturbations, because it did not place any sample in I2 in the wild type or in any condition except imatinib. Imatinib moved BioEmu2 out of the active class (91.2% in I1 and 8.8% in I2; Figure 10c). The imatinib-bound crystal structure has the DFG motif flipped out and the activation loop closed, so with imatinib Bio Emu2 reproduced the flipped DFG motif but closed the loop in only a minority of samples. With dasatinib, which binds the active form, 99.47% of samples stayed active.

BioEmu-1, run for the seven conditions without ligands or partner domains, kept I2 between 0.095% and 1.12% in every condition. The change that it produced had the same sign as the measured change in three of eight comparisons, was not detectable in four and had the opposite sign in one (Y412E raised I2, although phosphorylation of Y412 lowers both inactive states). The largest measured effect, T408Y (I2 from 6% to 70%), left the BioEmu-1 I2 population overlapping the wild-type range (0.199% to 0.231% against 0.177% to 0.210% over runs).

To test how BioEmu2 propagation connects the three states, we propagated NMR models of each state. Endpoints from the active models stayed active at every lag, and at 100 ns almost every other endpoint stayed in its start class (319 of 320 from I1 and 316 of 320 from I2). At 1,000 ns, 96 of the 320 I1-start endpoints were in the active class, but only eight of them lay within 3.0 Å of the active structure. Every change ran towards the active state, which BioEmu2 populates almost exclusively, so these endpoints may show relaxation towards the preferred state of the model rather than the connections between the states in solution.

To test whether BioEmu2 follows these changes step by step, we also chained 1,000 consecutive steps of 1 ns from five models of each state. None of the chains started from I1 or I2 reached the active class in any of its 10,010 frames, whereas single jumps of 1,000 ns from the same five starts placed 26% of the I1-start and 7.5% of the I2-start endpoints in the active class (Figure 10f). The chains also moved away from their start structures faster than single jumps of the same lag. At 10 ns, 32% of the I1-chain frames and 40% of the I2-chain frames lay within 3.0 Å of the start, against 76% and 90% of the single-jump endpoints (Figure 10e). Chained short steps and one long jump from the same structures therefore reached different ensembles. This suggests that the lag of BioEmu2 cannot be read as a physical time, although this comparison rests on five chains per state, whose frames are not independent (Supporting Information).

Taken together, these results suggest that neither model reproduces the populations of ABL1 or the way in which mutations shift them, and that the lag of BioEmu2 propagation should not be read as a physical time.

## 4. Conclusions

To conclude, in this study, we compared BioEmu-1 with the preview release of BioEmu2 on 22 systems, from the local geometry of the samples to the recovery of experimental states, the populations of these states and the response of these populations to perturbation. Four main findings emerged.

First, BioEmu2 samples almost never fail the backbone filter, whereas a median of 5.60% of BioEmu-1 samples per system fail it, so the second generation produces cleaner structures. Second, this cleaner geometry did not translate into reaching more states. BioEmu2 reached every state in 10 of the 22 systems and BioEmu-1 in 15. The difference lay mainly in states that need a ligand, nucleic acid, partner or phosphorylation that the sampled sequence lacks, and some of these states became accessible to BioEmu2 when the missing phosphorylation or DNA was supplied. Third, the two generations prefer different states in most systems, and BioEmu2 tends to favour the open form of proteins whose closed form was solved with a ligand. Fourth, neither generation reproduced the populations measured in solution. Only one of six comparisons fell within the experimental tolerance, and BioEmu2 tended to place almost all its samples in a single state.

For users of these models, these results would suggest three practical points. First, BioEmu2 samples rarely fail the backbone filter, but their local peptide geometry is still worth checking, and a few hundred of them capture almost all the states that BioEmu2 reaches at 3.0 Å. Second, a known partner or modification may be worth supplying when a bound or modified state is of interest. Third, the sampling frequencies of either model should not be read as free energies. States that lie far from the preferred state of BioEmu2 may still need a method with broader coverage. We caveat that the results from our work come from the preview weights, so a later release of BioEmu2 may behave differently and present an improvement, and the same kind of tests that we conducted may be repeated on it.

## Supporting Information

Supplementary methods (training data of the two models, condition labels, population targets, follow-up runs, scoring, backbone geometry, side-chain rebuild and statistical details); supplementary results (variation between runs, backbone geometry, sensitivity analyses of recovery, domain arrangements and comparison with small-angle X-ray scattering, pairwise RMSD and cluster count, populations, ABL1 and all-atom MD); Tables S1 to S17 and Figures S1 and S2 (PDF).

## Acknowledgement

This work was performed on the OzSTAR national facility at Swinburne University of Technology. The OzSTAR program receives funding in part from the Astronomy National Collaborative Research Infrastructure Strategy allocation provided by the Australian Government. Computations for this study were run on the facility’s OzSTAR and Ngarrgu Tindebeek supercomputers. The hardware for Ngarrgu Tindebeek was purchased through a grant from the Victorian Higher Education State Investment Fund. Claude Code (Anthropic; Claude Opus 5.5) was used to edit the language of the manuscript and Supporting Information, as well as to generate the code for data analysis and graph plotting. A human author was involved at every stage of this process. Specifically, the authors decided how to analyse the data, directed each code-generation step, checked the code and its outputs, interpreted all results, and proofread the final manuscript. The authors take full responsibility for the content of this article.

## Notes

### Competing Interest Statement

The authors have declared no competing interest.

