## Supplementary Information for "Does the BioEmu2 Preview Improve on BioEmu-1 for Protein Conformational Ensembles?"

#### Contents

S1. Supplementary methods; S2. Supplementary results; References. Tables S1 to S17; Figures S1 and S2.

Table S1. Release dates of the reference structures of each system, compared with the training data of each model.

Table S2. Condition labels of the 49 states.

Table S3. Components added in the eight BioEmu2 follow-up conditions.

Table S4. Families and finer labels of the 22 systems.

Figure S1. Difference between the two models against run-to-run variation for the 71 classes of the base condition.

Table S5. Results of the bioemu-benchmarks evaluator for the 12 benchmark-derived systems.

Table S6. Backbone geometry of filter-passing samples beyond the backbone filter, per system and method.

Table S7. The HIGH_QUALITY check of BioEmu2 and a heavy-atom clash score for BioEmu-1 samples with rebuilt side chains and for BioEmu2 samples, per system.

Table S8. Graded recovery of BioEmu-1 and BioEmu2 and their paired difference.

Table S9. Pairwise RMSD and cluster count of each ensemble, per system and method.

Table S10. Populations of the measured states outside ABL1 before renormalisation, with the unassigned fraction, for each method.

Table S11. Closed WPD-loop population of PTP1B on the Cα–Cα distance between Asp181 and Cys215.

Figure S2. Dependence of two population comparisons on the definition of the states.

Table S12. SHP2 N-SH2/PTP interface distance for each method and the structure of the open samples.

Table S13. ABL1 wild-type populations against the NMR values of Xie et al.

Table S14. ABL1 responses to perturbation for BioEmu-1 and BioEmu2.

Table S15. All-atom MD occupancy by start state.

Table S16. All-atom MD by start state over all production frames.

Table S17. Software, versions and hardware used in this study.

### S1. Supplementary methods

#### S1.1 Training data of the two models

The ood60 benchmark set was chosen for low similarity to the BioEmu-1 training data. Its proteins were selected because (a) they have been deposited after the training cut-off of AlphaFold2, and (b) share no more than 40% sequence similarity with the BioEmu-1 training set.[1] This choice does not apply to BioEmu2, which was trained on experimental structures from the PDB in the copy prepared for the structure-prediction model Boltz-1, together with predicted structures from the AlphaFold Protein Structure Database and new molecular dynamics data.[2] The BioEmu2 model card gives no release-date cut-off for these data. The reference structures released before the Boltz-1 cut-off[3] include eight of the 10 ood60 structures and both AdK structures. AdK is also the worked example in the BioEmu2 documentation,[4] although neither fact shows that AdK was in the training data. The details for each system are given in Table S1.

**Table S1.** Release dates of the reference structures of each system, compared with the training data of each model.

| System | Origin | Reference entries (PDB IDs) | Released | Released before the Boltz-1 cut-off | Held out of BioEmu-1 training |
|---|---|---|---|---|---|
| ABL1 | Chosen for this study | 6 | 2002 to 2020 | 6 of 6 | Likely (43.9%) |
| T4L L99A | Chosen for this study | 4 | 2008 to 2011 | 4 of 4 | No (32.2%) |
| CypA | Chosen for this study | 4 | 2009 | 4 of 4 | No |
| Aurora A | Chosen for this study | 13 | 2003 to 2019 | 13 of 13 | No (35.8%) |
| p38α | Chosen for this study | 4 | 2004 to 2016 | 4 of 4 | Yes (test protein) |
| ERK2 | Chosen for this study | 5 | 1998 to 2022 | 4 of 5 | Likely (50.3%) |
| MBP | Chosen for this study | 6 | 1994 to 2007 | 6 of 6 | Yes (test protein) |
| PTP1B | Chosen for this study | 6 | 1994 to 2018 | 6 of 6 | No |
| SHP2 | Chosen for this study | 5 | 1999 to 2020 | 5 of 5 | No |
| SRC | Chosen for this study | 7 | 1999 to 2026 | 5 of 7 | No (39.0%) |
| AdK | Chosen for this study | 2 | 1994 to 1996 | 2 of 2 | Yes (test protein) |
| RbsB | Benchmark, domain motion | 2 | 1995 to 1998 | 2 of 2 | Yes |
| DppA | Benchmark, domain motion | 2 | 1995 to 1996 | 2 of 2 | Yes |
| Hexokinase (*K. lactis*) | Benchmark, domain motion | 2 | 2010 | 2 of 2 | Yes |
| β-PGM | Benchmark, domain motion | 2 | 2009 to 2010 | 2 of 2 | Yes |
| Pol β | Benchmark, domain motion | 2 | 1994 | 2 of 2 | Yes |
| Biotin carboxylase | Benchmark, domain motion | 2 | 2000 | 2 of 2 | Yes |
| LapD module | Benchmark, low similarity (ood60) | 2 | 2019 | 2 of 2 | Yes |
| EcoT38I | Benchmark, low similarity (ood60) | 2 | 2022 | 0 of 2 | Yes |
| NanR | Benchmark, low similarity (ood60) | 2 | 2020 to 2021 | 2 of 2 | Yes |
| BusR | Benchmark, low similarity (ood60) | 2 | 2021 | 2 of 2 | Yes |
| Transposase HzTransib | Benchmark, low similarity (ood60) | 2 | 2019 | 2 of 2 | Yes |

In the last column, "yes" means that the protein is one of the published BioEmu-1 test proteins, which Lewis et al.[1] removed from the BioEmu-1 training data at 40% sequence similarity, and "likely" means that a test protein shares more than 40% sequence identity with it (best identity in brackets). The 84 unique entries comprise 79 released before 30 September 2021 and five released after it, namely two EcoT38I structures (PDB IDs: 7E8R and 7EDB), one ERK2 structure (PDB ID: 8AOJ) and two SRC structures (PDB IDs: 9NS0 and 9NS1). All 84 were released before BioEmu2 was trained in August and September 2026.[2] Release dates are from the RCSB PDB.

#### S1.2 Rules for the condition labels

Four labels were revised after a review of the structural evidence (Table S2), namely those of the folded-loop state of Aurora A (from B to C), unphosphorylated ERK2 (from C to A), the NMR intermediate of SRC (from A to C) and the open state of hexokinase (from B to C). The scored loop of the apo ERK2 structure (PDB ID: 5UMO) lies 0.29 and 0.48 Å from the two inactive reference structures, so the inactive loop conformation does not appear to depend on the ATP-site ligand of those references.

**Table S2.** Condition labels of the 49 states.

| No. | System | State | Reference structures (PDB IDs) | Label | Evidence |
|---|---|---|---|---|---|
| 1 | ABL1 | A | 6XR6 (NMR); 2F4J; 4TWP | A | 6XR6 solution NMR, no ligand, no mutation recorded; 2F4J VX-680 + H396P; 4TWP axitinib + T315I |
| 2 | ABL1 | I1 | 6XR7 (NMR); 1M52 | A | 6XR7 solution NMR, deposited with the wild-type sequence but calculated from restraints measured on the M309L/H415P variant,[5] no ligand; 1M52 PD173955 |
| 3 | ABL1 | I2 | 6XRG (NMR) | A | 6XRG solution NMR of the G269E/M309L/T408Y variant that stabilises I2 |
| 4 | Aurora A | Active loop, TPX2-bound | 1OL5; 5G1X; 3E5A; 3HA6 | B | TPX2 or N-Myc peptide bound in all four; pT287/pT288 or pT288; ADP or inhibitor |
| 5 | Aurora A | Active loop, no activator | 1MQ4; 4J8N; 6I2U; 4CEG; 5DNR | B | 1MQ4 ADP + Mg; 4J8N no ligand but T287D; 6I2U CoA + pT287/pT288; 4CEG ADP + pT288 + C290A/C393A; 5DNR ATP + pT288 |
| 6 | Aurora A | DFG-inter | 5L8K; 5L8L | B | ADP + pT288 + nanobody (chain B) + C290A/C393A in both |
| 7 | Aurora A | Inactive, folded loop | 2WTV; 4J8M | C | 2WTV MLN8054-type inhibitor (ZZL) + pT287 + mutations; 4J8M inhibitor (CJ5) + T287D |
| 8 | p38α | DFG-in, inactive loop | 1R39; 5ETC | A | Sulfate only in both |
| 9 | p38α | Active-like loop | 3PY3; 3OD6 | B | 3PY3 dually phosphorylated (pT180/pY182); 3OD6 Y323T (octyl glucoside is an additive) |
| 10 | ERK2 | Inactive, unphosphorylated | 8AOJ; 4GT3 | A | 8AOJ active-site ligand N8L (plus CME, DMSO, EDO, sulfate); 4GT3 ATP + Mg |
| 11 | ERK2 | Active, dually phosphorylated | 2ERK; 6OPG; 5V61 | B | Dually phosphorylated (pT183/pY185) in all; 6OPG AMP-PNP; 5V61 inhibitor + RSK peptide |
| 12 | SRC | Inactive, αC-out | 2SRC; 2H8H; 1KSW | B | pY527 (pY530 in the construct numbering) in all three; AMP-PNP, inhibitor or NBS; 1KSW T338G |
| 13 | SRC | Active, αC-in | 1Y57; 1YI6; 9NS0 (NMR) | B | 1Y57 inhibitor (MPZ); 1YI6 pY335/pY416; 9NS0 solution NMR with pY419 |
| 14 | SRC | NMR intermediate | 9NS1 (NMR) | C | Solution NMR, no ligand, no phosphorylation; sequence conflict A314I inside the scored αC segment (303-323) |
| 15 | PTP1B | WPD loop open | 2CM2; 2HNP; 6B8E; 1T49 | A | 2CM2 MPD only; 2HNP nothing; 6B8E glycerol/Tris + C32S/C92V; 1T49 allosteric inhibitor |
| 16 | PTP1B | WPD loop closed | 1SUG; 6B8E; 5K9W | A | 1SUG glycerol and Tris only; 6B8E glycerol/Tris + C32S/C92V; 5K9W ligand OTA |
| 17 | SHP2 | Closed | 4DGP; 2SHP; 5EHR; 7JVM | A | 4DGP no ligand (SEQADV lists only a deletion of 408-411); 2SHP cacodylate but engineered T2K/F41L/F513S, so it does not qualify; 5EHR SHP099; 7JVM |

| No. | System | State | Reference structures (PDB IDs) | Label | Evidence |
|---|---|---|---|---|---|
| | | | | | inhibitor |
| 18 | SHP2 | Open | 6CRF | B | E76K activating mutation |
| 19 | MBP | Open | 1OMP; 1JW4; 2V93 (NMR) | A | 1OMP and 1JW4 no ligand |
| 20 | MBP | Partially closed | 2V93 (NMR) | A | PRE ensemble of apo MBP carrying spin-label sites (D67C/S237C, label MXT) |
| 21 | MBP | Sugar-bound closed | 1ANF; 3MBP; 4MBP | B | Maltose or maltodextrin bound in all three |
| 22 | T4L L99A | Ground | 3DMV; 3DMX | A | 3DMV additives only (BME, HED, phosphate, chloride); 3DMX benzene; both C54T/C97A/L99A as in the construct |
| 23 | T4L L99A | Excited | 2LCB (NMR); 2LC9 (NMR) | A | 2LCB excited state of L99A (C54T/C97A/L99A) from CPMG chemical shifts; 2LC9 adds G113A/R119P |
| 24 | CypA | Major | 3K0M; 3K0N | A | No ligand, wild type |
| 25 | CypA | Minor | 3K0N; 3K0O; 3K0P | A | 3K0N wild-type multiconformer model holding the minor substate; 3K0O and 3K0P S99T |
| 26 | AdK | Open | 4AKE | A | No ligand |
| 27 | AdK | Closed | 1AKE | B | Ap5A bound |
| 28 | RbsB | Open | 1URP | A | No ligand |
| 29 | RbsB | Closed | 2DRI | B | Ribose bound |
| 30 | DppA | Open | 1DPE | A | $Cd^{2+}$ ions only (treated as an additive); no peptide |
| 31 | DppA | Closed | 1DPP | B | Gly-Leu dipeptide bound |
| 32 | Hexokinase | Open | 3O80 | C | AMP-PNP bound |
| 33 | Hexokinase | Closed | 3O8M | B | Glucose bound |
| 34 | β-PGM | Open | 2WFA | B | $BeF_3^-$ and $Mg^{2+}$ bound |
| 35 | β-PGM | Closed | 2WF5 | B | Glucose 6-phosphate and $MgF_3^-$ bound |
| 36 | Pol β | Apo | 1BPD | A | Phosphate only |
| 37 | Pol β | DNA ternary complex | 2BPG | B | Gapped DNA and ddCTP bound |
| 38 | Biotin carboxylase | Apo | 1DV1 | A | Phosphate only |
| 39 | Biotin carboxylase | ATP-bound | 1DV2 | B | ATP bound; E288K |
| 40 | LapD module | Apo | 6PWJ | A | $Mg^{2+}$ and PEG only |
| 41 | LapD module | c-di-GMP-bound | 6PWK | B | c-di-GMP and $Mg^{2+}$ bound; chain A binds two c-di-GMP (EAL site with $Mg^{2+}$, second site at residues 248-405) |
| 42 | EcoT38I | Apo | 7E8R | A | Glycerol only |
| 43 | EcoT38I | DNA-bound | 7EDB | B | DNA bound: homodimer (chains A, B) on one 13-bp duplex (chains E, F); two $Ca^{2+}$ at chain A's active site |
| 44 | NanR | Sialic-acid-bound | 6ON4 | B | N-acetylneuraminic acid (SLB) and $Zn^{2+}$ bound |
| 45 | NanR | DNA-bound | 6WFQ (cryo-EM) | B | DNA bound |
| 46 | BusR | DNA-bound a | 7OZ3 (cryo-EM) | B | DNA and c-di-AMP bound |
| 47 | BusR | DNA-bound b | 7B5Y (cryo-EM) | B | DNA and c-di-AMP bound |
| 48 | Transposase | Apo | 6PQN | A | Glycerol, phosphate, $Na^+$ and two $Zn^{2+}$ only |
| 49 | Transposase | DNA-bound | 6PQX (cryo-EM) | B | Hairpin DNA bound |

A, apo-matched; B, condition-dependent; C, uncertain. The evidence column lists the ligands, partner chains, phosphorylated residues and engineered mutations in each reference file, excluding crystallisation additives. The labels were assigned after the samples had been scored, and four were revised after a review of the structural evidence (rows 7, 10, 14 and 32). Residue numbers in the evidence column follow the deposited files. For SRC, pY527 in the deposited chicken numbering corresponds to Y530 in the UniProt numbering of the construct.

#### S1.3 Population targets

For T4L L99A, Mulder et al. report 97% ground state at 25 °C and standard errors for the enthalpy and entropy of the exchange, but not for the population,[6] and Korzhnev et al. fix the excited-state population at 3.4% from that earlier work.[7] For MBP, Tang et al. report that their fit to the paramagnetic data levels off at a minor-species population of about 5% and that the residual dipolar couplings agree best at 5% to 7%, and the same data exclude more than 1% of the sugar-bound closed form[8]; neither the main text nor the Supplementary Information of that study gives a standard uncertainty for these values. For CypA, only an upper bound of 10% on the minor population is available.[9] These values were therefore compared with the models descriptively, without a tolerance.

For Aurora A, the target of 52.4 ± 2.3% closed was derived from the reported equilibrium constant K (closed over open, 1.1 ± 0.1) as K/(1 + K), because the two reported percentages sum to 98%.[10] Using the reported 52 ± 1% instead narrowed the tolerance to ±2.0 pp but no result was changed, and the apparent free-energy errors changed by less than 0.01 kcal/mol. The measured protein carried the C290A and C393A substitutions and two dye-labelled cysteines, and it was attached to a surface, and the reported uncertainty is the propagated error of the histogram fit only, so it does not cover any effect of the labels, the surface attachment or the 10% glycerol in the buffer. The 22 Å cut-off of the K224–S283 Cα distance lies in the gap between the crystal structures: K224 and S283 are 15.1 and 17.6 Å apart in the two folded-loop structures (PDB IDs: 2WTV and 4J8M) and 35.9 to 42.0 Å apart in the eleven extended-loop structures. The measurement reports only whether the dyes touch so the closed count was repeated for every cut-off from 18 to 35.8 Å. The tolerance is ±4.5 pp.

#### S1.4 Follow-up runs with a partner or a phosphorylation

The components of each condition are listed in Table S3. No ligand condition was run for the folded-loop state of Aurora A (PDB IDs: 2WTV and 4J8M), for two reasons. First, both crystal structures contain an inhibitor, but the single-molecule data show the loop closed in about half of the molecules without any ligand, so this state does not clearly need a ligand. Second, the two inhibitors give loop conformations 5.4 Å apart, so a run with either inhibitor would test one complex rather than the ligand-free protein that the measurement describes.

The partner runs were compared with the probability that a random set of 1,000 samples from a protein-only run contains a sample within the threshold, computed exactly as for recovery. The phosphorylation runs, which used the same random seed as the unmodified sequence, were compared with the first 1,000 samples of each protein-only run.

**Table S3.** Components added in the eight BioEmu2 follow-up conditions.

| Condition | Components added | Source structure (PDB ID) | Target state (reference PDB IDs) |
|---|---|---|---|
| Pol β with DNA | Template and primer DNA strands (8 and 7 nucleotides); ddCTP (DCT); one $Mg^{2+}$ (MG) | 2BPG | DNA ternary complex (2BPG) |
| NanR with DNA | 15-base-pair operator DNA | 6WFQ | DNA-bound (6WFQ) |
| NanR with sialic acid | N-acetylneuraminic acid (SLB); one $Zn^{2+}$ (ZN) | 6ON4 | Sialic-acid-bound (6ON4) |
| BusR with DNA and cyclic di-AMP | 46-base-pair promoter DNA; one cyclic di-AMP (2BA) | 7B5Y | DNA-bound a (7OZ3) and DNA-bound b (7B5Y)[a] |
| LapD module with cyclic di-GMP | Two cyclic di-GMP (C2E); one $Mg^{2+}$ (MG) | 6PWK | c-di-GMP-bound (6PWK) |
| ERK2, pT185/pY187 | Phosphothreonine (TPO) at T185; phosphotyrosine (PTR) at Y187 | 2ERK, 6OPG, 5V61 | Active, dually phosphorylated (2ERK, 6OPG, 5V61) |
| p38α, pT180/pY182 | TPO at T180; PTR at Y182 | 3PY3 | Active-like loop (3PY3, 3OD6) |
| SRC, pY530 | PTR at Y530 | 2SRC, 2H8H, 1KSW | Inactive, αC-out (2SRC, 2H8H, 1KSW) |

Codes follow the PDB Chemical Component Dictionary. Only molecules that touch the reference protein chain in the bound structure were added, and crystallisation additives were left out. In every bound structure except that of Pol β, the reference chain also touches at least one other protein chain, which was not included. [a] The same input was used for both DNA-bound states of BusR, because the 152-base-pair DNA of the DNA-bound structure (PDB ID: 7OZ3) was too large to sample.

#### S1.5 Scoring details

Residues without a Cα atom in one or more reference structures of a system were left out of its superposition and scored residues. For Aurora A, this removed five superposition residues and residues 285 and 287 of the scored region, so 24 of its 26 scored residues were used. For SRC, the superposition residues (343 to 503) include the scored activation loop (407 to 435), so a refit without the loop was also scored as a sensitivity analysis.

#### S1.6 Backbone geometry beyond the filter

Ramachandran outliers and favoured residues were assigned from the φ and ψ angles with the Top500 density grids of Lovell et al.,[11] in the four classes general, glycine, proline and pre-proline, as distributed in PyRAMA 2.0.2. A residue was an outlier when the density at its (φ, ψ) was below 0.05% (general class) or 0.2% (the other classes), and favoured when it was at least 2%. Each peptide bond was classed by its ω dihedral angle (Cα–C–N–Cα) as cis (|ω| below 30°), twisted (30° to 150°) or trans. The N–Cα–C angle (τ) and the backbone bond lengths were compared with the targets of Engh and Huber[12]: τ 111.0 ± 2.7° (113.1 ± 2.5° for glycine and 112.1 ± 2.6° for proline), N–Cα 1.459 ± 0.020 Å, Cα–C 1.525 ± 0.026 Å, C–N 1.336 ± 0.023 Å and C=O 1.229 ± 0.019 Å. A value more than 4 standard deviations from its target was counted as an outlier, as in MolProbity.[13] Values were pooled over the samples of each system and summarised by the median over systems.

#### S1.7 HIGH_QUALITY check, side-chain rebuild and clash score

The HIGH_QUALITY check is part of the BioEmu2 code. It rejects a sample when more than five backbone bonds lie outside fixed length windows (for example 1.05 to 1.65 Å for the peptide C–N bond), or when the counts of clashes or of other bond and valence errors exceed set limits. For BioEmu2, the verdict written by the model was used for every sample. To test whether BioEmu-1 samples would also pass this check, the check was rerun with the same code (bioemu2.sampling.check_physicality in high-quality mode) on a random subset of 1,000 filter-passing samples per run of both models, the subset used for the pairwise RMSD and geometry analyses; for BioEmu2, the rerun gave the same verdict as the model for all 66,000 samples.

The side chains of BioEmu-1 samples were rebuilt with HPacker[14] in the same way as the side-chain reconstruction of bioemu 1.2.0 (backbone N, Cα, C and O as input, five refinement iterations), without the molecular dynamics relaxation that bioemu can add afterwards. The same rebuilt side chains gave the BioEmu-1 rotamers of Phe113 in CypA and Phe114 in T4L L99A. The check reports only the first limit that a sample exceeds, so every limit was also tested separately on part of the subset (375 or 750 samples per system). To estimate how much the rebuild itself contributes, the side chains of 1,000 BioEmu2 samples per run of CypA and T4L L99A were removed and rebuilt in the same way.

The heavy-atom clash score was the number of pairs of heavy atoms in different residues, more than three covalent bonds apart, whose van der Waals radii[15] overlap by at least 0.4 Å, per 1,000 heavy atoms. Pairs of nitrogen and oxygen atoms were counted only below 2.4 Å, because closer approaches between them are usually hydrogen bonds, and disulfide sulfur pairs were not counted. Clashing pairs were also split into pairs of two backbone atoms (N, Cα, C and O), pairs of a backbone atom and a side-chain atom, and pairs of two side-chain atoms, with Cβ counted as a side-chain atom because HPacker rebuilds it. This score uses heavy atoms only and is therefore not the MolProbity clashscore.

#### S1.8 Statistical details

A simulation of 4,000 studies per setting showed that the paired binary test has low power with 22 systems. For example, its power is 0.155 for an improvement of 1.0 in log-odds.

**Table S4.** Families and finer labels of the 22 systems.

| System | Family (14) | Finer label (20) |
|---|---|---|
| ABL1 | Protein kinase | tyrosine kinase abl |
| Aurora A | Protein kinase | protein kinase Other Aur |
| p38α | Protein kinase | protein kinase CMGC MAPK |
| ERK2 | Protein kinase | protein kinase CMGC MAPK |
| SRC | Protein kinase | protein kinase TK Src |
| PTP1B | Protein tyrosine phosphatase | protein tyrosine phosphatase classical |
| SHP2 | Protein tyrosine phosphatase | protein tyrosine phosphatase classical |
| MBP | Periplasmic binding protein | periplasmic binding protein |
| RbsB | Periplasmic binding protein | periplasmic binding protein type1 (PF13407) |
| DppA | Periplasmic binding protein | sbp bac 5 (PF00496) |
| T4L L99A | T4 lysozyme | t4 lysozyme |
| CypA | Cyclophilin | cyclophilin |
| AdK | Adenylate kinase | adenylate kinase (PF00406) |
| Hexokinase | Hexokinase | hexokinase (PF00349, PF03727) |
| β-PGM | HAD hydrolase | had hydrolase (PF00702) |
| Pol β | DNA polymerase X | dna polymerase x (PF14791, PF14792) |
| Biotin carboxylase | ATP-grasp | biotin carboxylase (PF00289, PF02785, PF02786) |
| LapD module | GGDEF-EAL | ggdef eal (PF00990, PF00563) |
| EcoT38I | SacI-like restriction endonuclease | saci restriction endonuclease (PF09566, PF29187) |
| NanR | GntR-family regulator | gntr fcd (PF00392, PF07729) |
| BusR | GntR-family regulator | gntr trka c (PF00392, PF02080) |
| Transposase | Transib transposase | transib transposase |

The 14 families were used for the two-stage bootstrap and the sign-flip test, and the 20 finer labels were used in the sensitivity analysis. Finer labels follow the Pfam domains of each system (accessions in brackets).

#### S2. Supplementary results

##### S2.1 Variation between runs

The 22 systems in the base condition have 49 states and 71 classes, because each system has one unassigned class in addition to its states. For each class, we compared the difference between the populations of the two models with the variation between the three runs of each model (Figure S1).

The largest run-to-run SD, 0.82 pp, was that of the closed state of RbsB in BioEmu2, whose population was 55.0 ± 0.8% (mean ± SD). In the closest of the 54 classes in which the two models differed by more than 1 pp, the unassigned class of Aurora A, the models differed by 1.95 pp, whereas the larger SD was 0.53 pp. In the remaining 17 classes, the populations of the two models agreed to within 1 pp. In 15 of these classes, both models placed fewer than 1% of their samples in the class, and in six they placed none, so this agreement mostly reflects states that neither model sampled often. The other two were unassigned classes: both models left almost every BusR sample unassigned, and both left about 41% of the AdK samples unassigned (41.0% for BioEmu-1 and 40.8% for BioEmu2).

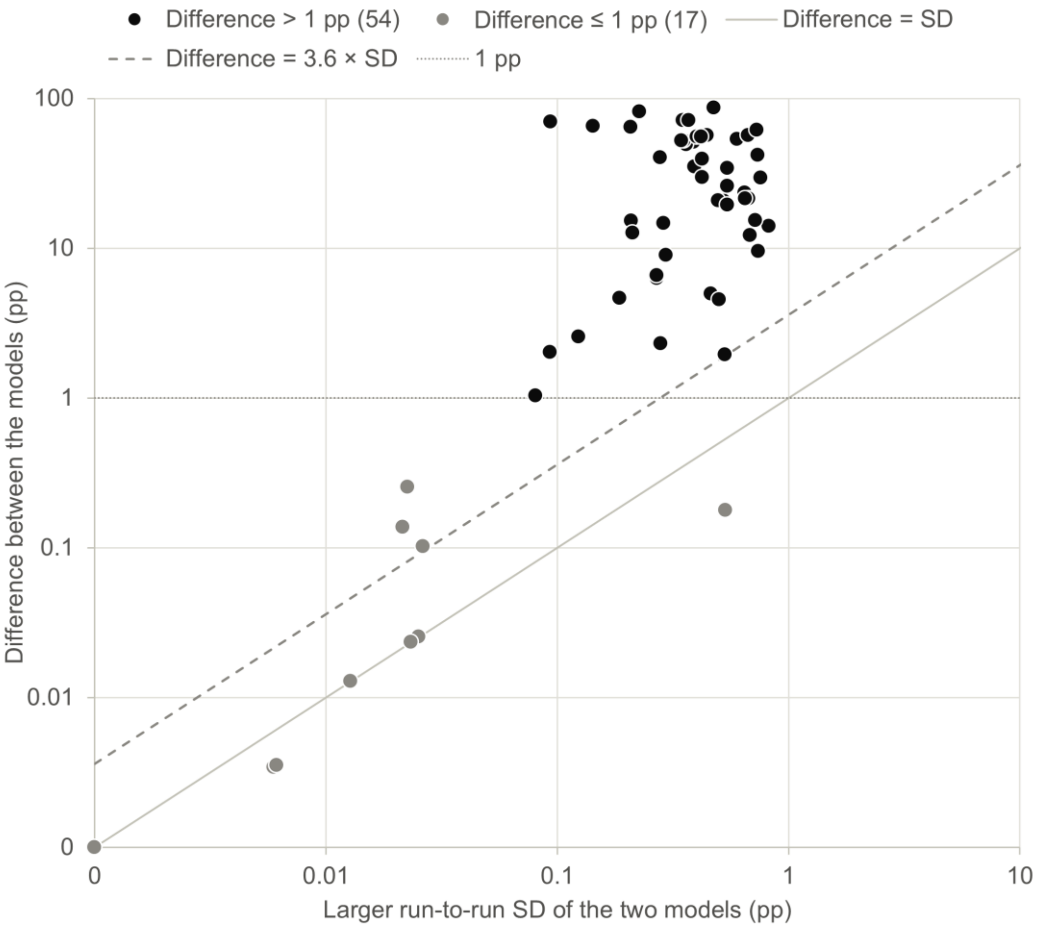


**Figure S1. Difference between the two models against run-to-run variation for the 71 classes of the base condition.** Each point is one class (a state or the unassigned class of a system). The horizontal axis gives the larger of the two run-to-run standard deviations (SDs) of the class population (BioEmu-1 or BioEmu2, three runs each), and the vertical axis gives the absolute difference between the mean populations of the two models. Both quantities are in percentage points (pp) of filter-passing samples on logarithmic axes, and values of zero are drawn at the axis minimum and labelled 0. The solid line marks a difference equal to the SD, the dashed line a difference 3.6 times the SD and the dotted line a difference of 1 pp. All 54 classes whose difference exceeds 1 pp lie on or above the dashed line.

#### S2.2 Backbone geometry

The bioemu-benchmarks evaluator, which applies its own sample filter, backbone RMSD and scored regions, gave the same picture as our scoring (Table S5).

The BioEmu2 failure fraction ranged from 0% for SRC and T4L L99A to 0.090% for NanR. The BioEmu-1 failures were spread over the three checks of the filter, and a sample could fail more than one. Averaged over all systems, 4.73% of samples had at least one peptide C–N distance of 2.0 Å or more, 3.56% had at least one Cα–Cα distance of 4.5 Å or more between consecutive residues, and 4.02% had a clash. Broken chains were more frequent than clashes: in 14 of the 22 systems, one of the two break criteria rejected more samples than the clash criterion.

**Table S5.** Results of the bioemu-benchmarks evaluator for the 12 benchmark-derived systems.

| Quantity | BioEmu-1 | BioEmu2 |
|---|---|---|
| Domain-motion coverage at 3.0 Å, bootstrapped (11 systems) | 0.837 / 0.825 / 0.841 | 0.727 / 0.727 / 0.727 |
| ood60 coverage at 3.0 Å, bootstrapped (BusR, local metric) | 0.293 / 0.260 / 0.100 | 0.000 / 0.000 / 0.000 |
| References reached at 3.0 Å, mean over 12 systems | 0.875 / 0.833 / 0.875 | 0.667 / 0.667 / 0.667 |
| Samples passing the evaluator's own filter | 76.2% to 99.2% | 99.75% to 100% |
| Reached-at-3.0 Å calls that agree with this study's scoring (reference and run pairs) | 72 of 72 | 72 of 72 |

The evaluator was version 0.0.2.post1, and values are given for runs 1, 2 and 3. The domain-motion task of the evaluator covers 11 of the 12 systems (AdK, the six domain-motion cases and the four ood60 cases that also appear in the domain-motion set), and its ood60 task covers BusR.

The median share of Ramachandran outliers was 0.72% for BioEmu-1, 0.46% for BioEmu2, 0.59% for all-atom MD and 1.54% for ColabFold (Table S6). Whether the Gly274–Ser275 cis peptide bond of the BioEmu2 Pol β samples occurs in other Pol β structures was not checked.

In the HIGH_QUALITY check (Table S7), a median of 23.4% of the rebuilt BioEmu-1 samples passed, against 65,998 of the 66,000 BioEmu2 samples. Most BioEmu-1 samples failed because of their backbone, which the rebuild leaves in place, since a median of 62.4% had more than five backbone bonds outside the length windows of the check. The heavy-atom clash score had a median of 45.5 for the rebuilt BioEmu-1 samples against 4.9 for BioEmu2. A control rebuild of the BioEmu2 samples of CypA and T4L L99A suggested that the rebuild alone can account for roughly half of the BioEmu-1 clash score, but for only a small part of its failures of the check. Clashes between two backbone atoms, which do not depend on the side chains, were about six times more frequent in BioEmu-1 (median 10.2 against 1.6 per 1,000 heavy atoms). The BioEmu-1 backbone therefore appears to be less regular, although its exact pass rate and clash score also depend on the rebuild.

**Table S6.** Backbone geometry of filter-passing samples beyond the backbone filter, per system and method.

| System | Method | Samples | Ramachandran outliers (% of residues) | Ramachandran favoured (%) | Cis non-proline peptides (% of peptide bonds) | Twisted peptides (%) | Mean deviation of trans ω from 180° (°) | τ SD (°) | τ outliers (%) | N–Cα outliers (%) | C–N outliers (%) |
|---|---|---|---|---|---|---|---|---|---|---|---|
| ABL1 | BioEmu-1 | 3000 | 1.117 | 91.449 | 0.002 | 0.435 | 7.178 | 0.802 | 0 | 0 | 48.202 |
| ABL1 | BioEmu2 | 3000 | 0.253 | 94.838 | 0.002 | 0.211 | 7.515 | 3.276 | 0.23 | 1.265 | 0.151 |
| ABL1 | ColabFold | 500 | 1.731 | 95.295 | 0 | 0.42 | 4.144 | 0.792 | 0 | 0 | 4.827 |
| ABL1 | All-atom MD | 3000 | 0.586 | 91.994 | 0 | 0.322 | 7.835 | 3.353 | 0.256 | 2.111 | 0.107 |
| Aurora A | BioEmu-1 | 3000 | 0.908 | 93.538 | 0.006 | 0.511 | 6.133 | 0.781 | 0 | 0 | 45.07 |
| Aurora A | BioEmu2 | 3000 | 0.496 | 94.243 | 0.011 | 0.241 | 7.537 | 3.325 | 0.297 | 1.311 | 0.16 |
| Aurora A | ColabFold | 500 | 4.921 | 91.117 | 0 | 2.549 | 4.405 | 0.771 | 0 | 0 | 3.024 |
| p38α | BioEmu-1 | 3000 | 0.684 | 94.816 | 0.004 | 0.278 | 5.348 | 0.777 | 0 | 0 | 43.701 |
| p38α | BioEmu2 | 3000 | 0.423 | 94.399 | 0.009 | 0.253 | 7.404 | 3.294 | 0.256 | 1.349 | 0.171 |
| p38α | ColabFold | 499 | 2.105 | 95.573 | 0 | 0.227 | 3.826 | 0.77 | 0 | 0 | 3.109 |
| ERK2 | BioEmu-1 | 3000 | 1.317 | 92.355 | 0.004 | 0.202 | 6.377 | 0.812 | 0 | 0 | 46.743 |
| ERK2 | BioEmu2 | 3000 | 0.784 | 93.978 | 0.006 | 0.24 | 7.421 | 3.342 | 0.31 | 1.401 | 0.185 |
| ERK2 | ColabFold | 498 | 3.192 | 93.134 | 0 | 0.146 | 4.074 | 0.806 | 0 | 0 | 3.661 |
| SRC | BioEmu-1 | 3000 | 0.646 | 95.678 | 0.003 | 0.26 | 5.758 | 0.823 | 0 | 0 | 44.763 |
| SRC | BioEmu2 | 3000 | 0.342 | 94.848 | 0.011 | 0.205 | 7.662 | 3.328 | 0.271 | 1.406 | 0.173 |
| SRC | ColabFold | 498 | 1.672 | 95.153 | 0 | 0.432 | 4.224 | 0.825 | 0 | 0 | 4.901 |
| PTP1B | BioEmu-1 | 3000 | 1.224 | 93.161 | 0.003 | 0.258 | 6.228 | 0.799 | 0 | 0 | 45.696 |
| PTP1B | BioEmu2 | 3000 | 0.559 | 94.831 | 0.013 | 0.271 | 7.481 | 3.347 | 0.279 | 1.305 | 0.154 |
| PTP1B | ColabFold | 499 | 2.701 | 93.949 | 0 | 0.407 | 4.465 | 0.79 | 0 | 0 | 3.367 |
| PTP1B | All-atom MD | 3000 | 0.84 | 92.268 | 0 | 0.302 | 7.658 | 3.466 | 0.307 | 2.42 | 0.144 |
| SHP2 | BioEmu-1 | 3000 | 1.373 | 93.928 | 0.006 | 0.501 | 5.739 | 0.851 | 0 | 0 | 46.573 |
| SHP2 | BioEmu2 | 3000 | 0.664 | 94.006 | 0.018 | 0.275 | 7.667 | 3.429 | 0.299 | 1.535 | 0.203 |
| SHP2 | ColabFold | 489 | 3.039 | 92.899 | 0.001 | 0.647 | 4.395 | 0.861 | 0 | 0 | 6.569 |
| MBP | BioEmu-1 | 3000 | 0.636 | 94.992 | 0.001 | 0.066 | 6.231 | 0.879 | 0 | 0 | 45.706 |
| MBP | BioEmu2 | 3000 | 0.492 | 95.883 | 0.008 | 0.171 | 7.412 | 3.313 | 0.244 | 1.397 | 0.182 |
| MBP | ColabFold | 500 | 0.226 | 98.655 | 0 | 0.001 | 4.057 | 0.872 | 0 | 0 | 1.79 |
| T4L L99A | BioEmu-1 | 3000 | 0.264 | 96.567 | 0 | 0.029 | 5.202 | 0.796 | 0 | 0 | 42.58 |
| T4L L99A | BioEmu2 | 3000 | 0.102 | 96.507 | 0 | 0.054 | 7.067 | 3.162 | 0.22 | 1.323 | 0.135 |
| T4L L99A | ColabFold | 500 | 0.178 | 99.447 | 0 | 0 | 3.611 | 0.777 | 0 | 0 | 2.67 |
| T4L L99A | All-atom MD | 3000 | 0.28 | 95.047 | 0 | 0.229 | 7.506 | 3.343 | 0.298 | 2.424 | 0.141 |
| CypA | BioEmu-1 | 3000 | 1.88 | 89.527 | 0.002 | 0.498 | 7.841 | 0.978 | 0 | 0 | 50.456 |
| CypA | BioEmu2 | 3000 | 1.222 | 92.212 | 0.002 | 0.398 | 7.826 | 3.45 | 0.287 | 1.3 | 0.148 |
| CypA | ColabFold | 499 | 1.3 | 96.802 | 0 | 0.01 | 4.58 | 0.96 | 0 | 0 | 3.063 |
| CypA | All-atom MD | 3000 | 1.25 | 91.055 | 0 | 0.602 | 8.129 | 3.538 | 0.298 | 2.09 | 0.109 |
| AdK | BioEmu-1 | 3000 | 0.759 | 93.943 | 0 | 0.248 | 7.011 | 0.932 | 0 | 0 | 47.87 |
| AdK | BioEmu2 | 3000 | 0.119 | 97.228 | 0.001 | 0.171 | 7.416 | 3.282 | 0.217 | 1.287 | 0.144 |
| AdK | ColabFold | 500 | 0 | 99.779 | 0 | 0 | 3.606 | 0.918 | 0 | 0 | 3.341 |
| AdK | All-atom MD | 3000 | 0.184 | 96.021 | 0 | 0.259 | 7.78 | 3.462 | 0.288 | 2.44 | 0.151 |

| System | Method | Samples | Ramachandran outliers (% of residues) | Ramachandran favoured (%) | Cis non-proline peptides (% of peptide bonds) | Twisted peptides (%) | Mean deviation of trans ω from 180° (°) | τ SD (°) | τ outliers (%) | N–Cα outliers (%) | C–N outliers (%) |
|---|---|---|---|---|---|---|---|---|---|---|---|
| RbsB | BioEmu-1 | 3000 | 0.89 | 94.744 | 0 | 0.241 | 6.863 | 0.906 | 0 | 0 | 47.136 |
| RbsB | BioEmu2 | 3000 | 0.449 | 96.314 | 0 | 0.189 | 7.452 | 3.29 | 0.196 | 1.278 | 0.15 |
| RbsB | ColabFold | 500 | 0.374 | 98.772 | 0 | 0 | 4.091 | 0.894 | 0 | 0 | 2.08 |
| DppA | BioEmu-1 | 3000 | 0.231 | 96.119 | 0 | 0.081 | 5.729 | 0.845 | 0 | 0 | 43.819 |
| DppA | BioEmu2 | 3000 | 0.294 | 94.888 | 0.001 | 0.247 | 7.588 | 3.391 | 0.268 | 1.428 | 0.2 |
| DppA | ColabFold | 500 | 0.096 | 97.962 | 0 | 0.002 | 4.489 | 0.843 | 0 | 0 | 3.126 |
| Hexokinase | BioEmu-1 | 3000 | 0.373 | 96.024 | 0.002 | 0.068 | 5.353 | 0.923 | 0 | 0 | 43.381 |
| Hexokinase | BioEmu2 | 3000 | 0.493 | 94.673 | 0.01 | 0.216 | 7.534 | 3.362 | 0.271 | 1.521 | 0.198 |
| Hexokinase | ColabFold | 499 | 1.406 | 96.449 | 0 | 0.049 | 4.026 | 0.921 | 0 | 0 | 3.758 |
| β-PGM | BioEmu-1 | 3000 | 0.495 | 94.858 | 0 | 0.294 | 6.011 | 0.852 | 0 | 0 | 45.196 |
| β-PGM | BioEmu2 | 3000 | 0.155 | 96.9 | 0.033 | 0.404 | 7.249 | 3.237 | 0.221 | 1.3 | 0.149 |
| β-PGM | ColabFold | 500 | 0.269 | 98.134 | 0 | 0.181 | 3.345 | 0.836 | 0 | 0 | 3.936 |
| Pol β | BioEmu-1 | 3000 | 0.334 | 96.596 | 0.003 | 0.098 | 5.026 | 0.829 | 0 | 0 | 43.145 |
| Pol β | BioEmu2 | 3000 | 0.309 | 95.466 | 0.314 | 0.179 | 7.424 | 3.274 | 0.217 | 1.439 | 0.182 |
| Pol β | ColabFold | 500 | 2.708 | 95.67 | 0 | 0.026 | 3.6 | 0.824 | 0 | 0 | 2.651 |
| Biotin carboxylase | BioEmu-1 | 3000 | 0.843 | 96.435 | 0.001 | 0.129 | 5.234 | 0.948 | 0 | 0 | 42.275 |
| Biotin carboxylase | BioEmu2 | 3000 | 1.081 | 94.883 | 0.006 | 0.2 | 7.367 | 3.381 | 0.281 | 1.401 | 0.18 |
| Biotin carboxylase | ColabFold | 497 | 0.817 | 97.875 | 0 | 0.073 | 3.823 | 0.946 | 0 | 0 | 3.682 |
| LapD module | BioEmu-1 | 3000 | 0.642 | 95.468 | 0.004 | 0.287 | 5.058 | 0.782 | 0 | 0 | 45.399 |
| LapD module | BioEmu2 | 3000 | 0.409 | 94.837 | 0.007 | 0.164 | 7.38 | 3.298 | 0.263 | 1.505 | 0.19 |
| LapD module | ColabFold | 499 | 1.302 | 96.301 | 0 | 0.11 | 3.547 | 0.784 | 0 | 0 | 5.12 |
| EcoT38I | BioEmu-1 | 3000 | 0.512 | 95.473 | 0.003 | 0.208 | 4.918 | 0.814 | 0 | 0 | 44.89 |
| EcoT38I | BioEmu2 | 3000 | 0.56 | 94.558 | 0.009 | 0.187 | 7.294 | 3.274 | 0.24 | 1.387 | 0.162 |
| EcoT38I | ColabFold | 500 | 0.7 | 97.053 | 0 | 0.106 | 3.705 | 0.811 | 0 | 0 | 4.178 |
| NanR | BioEmu-1 | 3000 | 2.007 | 94.102 | 0.188 | 2.03 | 4.771 | 0.769 | 0 | 0 | 45.866 |
| NanR | BioEmu2 | 3000 | 0.463 | 94.607 | 0.016 | 0.235 | 7.528 | 3.23 | 0.303 | 1.511 | 0.196 |
| NanR | ColabFold | 500 | 6.611 | 90.074 | 0 | 4.706 | 3.666 | 0.786 | 0 | 0 | 4.365 |
| BusR | BioEmu-1 | 3000 | 1.247 | 94.521 | 0.02 | 0.594 | 5.315 | 0.864 | 0 | 0 | 47.648 |
| BusR | BioEmu2 | 3000 | 0.464 | 94.184 | 0.008 | 0.322 | 7.628 | 3.353 | 0.33 | 1.549 | 0.202 |
| BusR | ColabFold | 498 | 5.347 | 92.169 | 0 | 1.169 | 3.774 | 0.921 | 0 | 0 | 6.991 |
| Transposase | BioEmu-1 | 3000 | 0.661 | 95.262 | 0.004 | 0.329 | 5.325 | 0.766 | 0 | 0 | 43.54 |
| Transposase | BioEmu2 | 3000 | 0.411 | 94.596 | 0.009 | 0.226 | 7.497 | 3.299 | 0.3 | 1.582 | 0.215 |
| Transposase | ColabFold | 499 | 3.343 | 94.424 | 0 | 0.842 | 4.033 | 0.771 | 0 | 0 | 3.663 |

Samples are 1,000 random filter-passing samples per run for BioEmu-1 and BioEmu2 (3,000 per system), all filter-passing ColabFold models and 500 frames per all-atom replica after the first 10 ns. Ramachandran classes follow the Top500 grids of Lovell et al.[11] A peptide bond is cis when |ω| is below 30° and twisted when |ω| is between 30° and 150°. Bond lengths and the N–Cα–C angle (τ) are compared with the targets of Engh and Huber,[12] and an outlier lies more than 4 standard deviations from its target. τ SD is the standard deviation of τ. Across the 22 systems, the medians for Ramachandran outliers, C–N outliers and τ SD were 0.722%, 45.297% and 0.826° for BioEmu-1. The corresponding medians were 0.456%, 0.176% and 3.306° for BioEmu2 and 1.539%, 3.662% and 0.824° for ColabFold. Over the five all-atom MD systems, the medians were 0.586%, 0.141% and 3.462°. The N–Cα outliers of BioEmu2 and all-atom MD reflect a mean N–Cα length of about 1.470 Å against the 1.459 Å target, which is a shift of the mean rather than scatter.

**Table S7.** The HIGH_QUALITY check of BioEmu2 and a heavy-atom clash score for BioEmu-1 samples with rebuilt side chains and for BioEmu2 samples, per system.

| System | BioEmu-1 rebuilt: HIGH_QUALITY pass (%) | BioEmu-1 rebuilt: first failure on backbone bonds (%) | BioEmu-1 rebuilt: pass without the backbone bond limit (%) | BioEmu-1 rebuilt: median clash score | BioEmu-1 rebuilt: backbone–backbone clashes per 1,000 heavy atoms | BioEmu2: HIGH_QUALITY pass (%) | BioEmu2: median clash score | BioEmu2: backbone–backbone clashes per 1,000 heavy atoms |
|---|---|---|---|---|---|---|---|---|
| ABL1 | 9.567 | 79.9 | 41.6 | 66.552 | 10.157 | 100 | 4.294 | 1.515 |
| Aurora A | 40.4 | 47.633 | 72.8 | 46.441 | 9.281 | 99.967 | 4.774 | 1.544 |
| p38α | 23.833 | 56.6 | 53.6 | 45.767 | 10.842 | 100 | 5.162 | 1.599 |
| ERK2 | 6.3 | 78.667 | 23.2 | 61.544 | 9.129 | 100 | 4.52 | 1.322 |
| SRC | 5.933 | 81.9 | 31.2 | 46.396 | 10.539 | 100 | 4.143 | 1.457 |
| PTP1B | 14.733 | 63.467 | 34.4 | 62.929 | 11.811 | 100 | 4.958 | 1.651 |
| SHP2 | 0.833 | 97.233 | 17.333 | 47.53 | 10.753 | 100 | 6.088 | 1.603 |
| MBP | 22.9 | 66.733 | 65.333 | 38.929 | 8.512 | 100 | 4.171 | 1.799 |
| T4L L99A | 81.7 | 6.833 | 86.267 | 45.211 | 8.792 | 100 | 2.299 | 1.699 |
| CypA | 44.233 | 53.7 | 94.933 | 49.012 | 9.233 | 100 | 2.372 | 0.795 |
| AdK | 40.167 | 52.967 | 83.467 | 48.943 | 10.198 | 100 | 4.23 | 1.752 |
| RbsB | 35.633 | 61.4 | 90.267 | 36.482 | 9.914 | 99.967 | 3.498 | 1.655 |
| DppA | 7.7 | 75.633 | 30.133 | 46.948 | 7.91 | 100 | 5.683 | 1.381 |
| Hexokinase | 22.933 | 69.733 | 71.2 | 37.931 | 9.658 | 100 | 5.04 | 1.619 |
| β-PGM | 58.367 | 28.3 | 79.467 | 43.351 | 9.497 | 100 | 2.929 | 1.692 |
| Pol β | 54.667 | 36.667 | 88.533 | 34.125 | 10.795 | 100 | 4.451 | 1.823 |
| Biotin carboxylase | 10.133 | 54.333 | 21.333 | 46.532 | 7.687 | 100 | 5.78 | 1.406 |
| LapD module | 13 | 82.633 | 69.333 | 39.533 | 11.47 | 100 | 5.091 | 1.633 |
| EcoT38I | 29.133 | 59.433 | 74.133 | 42.685 | 10.796 | 100 | 5.108 | 1.939 |
| NanR | 32.9 | 65.3 | 92 | 35.611 | 14.679 | 100 | 7.7 | 2.289 |
| BusR | 42.567 | 55.567 | 94.133 | 33.333 | 13.728 | 100 | 6.548 | 2.038 |
| Transposase | 13.033 | 77.567 | 53.067 | 40.503 | 10.911 | 100 | 6.038 | 1.977 |

Each value uses 1,000 random filter-passing samples per run (3,000 per system). Over all 30,000 samples of each system, the verdict that BioEmu2 wrote passed at least 99.98% of them. BioEmu-1 side chains were rebuilt with HPacker without relaxation. The check reports only the first limit that a sample exceeds, and the column for the backbone bond limit gives the share of samples that failed first on more than five backbone bonds outside the length windows. The pass rate without that limit and the backbone–backbone clashes come from a part of the subset (375 or 750 samples per system). In a control, BioEmu2 samples were rebuilt in the same way (side chains removed and rebuilt with HPacker, 3,000 samples per system). The HIGH_QUALITY pass rate was then 99.767% for CypA and 95.933% for T4L L99A, with median clash scores of 20.553 and 27.586. The rebuild did not move any backbone atom (largest shift 0.0000 Å).

#### S2.3 Sensitivity analyses of recovery

The graded recovery of the two models at each threshold and budget is given in Table S8.

**Table S8.** Graded recovery of BioEmu-1 and BioEmu2 and their paired difference.

| Threshold (Å) | Budget (samples) | BioEmu-1 | BioEmu2 | Difference (95% CI) | Systems B2 higher / B1 higher / equal | Sign-flip *p* |
|---|---|---|---|---|---|---|
| 1.5 | 100 | 0.376 | 0.385 | +0.009 (−0.120 to 0.113) | 7 / 10 / 5 | 0.435 |
| 1.5 | 1,000 | 0.464 | 0.434 | −0.030 (−0.130 to 0.056) | 6 / 10 / 6 | 0.186 |
| 1.5 | 10,000 | 0.537 | 0.481 | −0.056 (−0.176 to 0.056) | 3 / 7 / 12 | 0.270 |
| 1.5 | Budget-averaged | 0.460 | 0.434 | −0.026 (−0.127 to 0.060) | 6 / 11 / 5 | 0.196 |
| 2.0 | 100 | 0.515 | 0.478 | −0.037 (−0.155 to 0.072) | 7 / 13 / 2 | 0.322 |
| 2.0 | 1,000 | 0.636 | 0.563 | −0.074 (−0.170 to 0.042) | 6 / 12 / 4 | 0.227 |
| 2.0 | 10,000 | 0.705 | 0.636 | −0.068 (−0.167 to 0.043) | 2 / 6 / 14 | 0.219 |
| 2.0 | Budget-averaged | 0.626 | 0.562 | −0.064 (−0.153 to 0.042) | 7 / 13 / 2 | 0.192 |
| 3.0 | 100 | 0.706 | 0.660 | −0.046 (−0.148 to 0.063) | 6 / 10 / 6 | 0.372 |
| 3.0 | 1,000 | 0.779 | 0.670 | −0.108 (−0.251 to 0.030) | 2 / 8 / 12 | 0.109 |
| 3.0 | 10,000 | 0.838 | 0.670 | −0.168 (−0.337 to 0.000) | 1 / 8 / 13 | 0.125 |
| 3.0 | Budget-averaged | 0.776 | 0.669 | −0.107 (−0.244 to 0.028) | 5 / 11 / 6 | 0.106 |

Values are means over the 22 systems and the paired difference BioEmu2 − BioEmu-1, with the 95% two-stage family-bootstrap confidence interval (CI, 20,000 resamples) and the exact sign-flip test on the 14 family means. B1, BioEmu-1; B2, BioEmu2.

To test whether the graded-recovery results depended on specific analytical choices, we conducted four sensitivity analyses. First, when systems were grouped into 20 smaller families instead of 14 (Table S4), the 3.0 Å difference at 10,000 samples was −0.168 (95% CI −0.333 to −0.025). Second, when a sample counted for a state only if that state was also its nearest one, the budget-averaged 3.0 Å difference was −0.137 (−0.269 to −0.005). While both intervals excluded zero in favour of BioEmu-1, they should be interpreted with caution: because they come from a bootstrap over few families, the intervals may be too narrow.

The remaining two sensitivity analyses evaluated the influence of single families and of one system. Leaving out one family at a time kept the 3.0 Å difference negative in all 14 cases (−0.064 to −0.140). Similarly, removing CypA, whose two states differ by only 0.11 Å in the backbone, changed the differences very little. Without CypA, the budget-averaged differences were −0.027, −0.067 and −0.112 (95% CI −0.254 to 0.027) at 1.5, 2.0 and 3.0 Å, against −0.026, −0.064 and −0.107 with all 22 systems, and the sign-flip *p* values were unchanged. No analysis gave an interval that excluded zero at 1.5 or 2.0 Å.

When the systems were grouped by origin, the direction of the difference was the same in each group at 2.0 and 3.0 Å, although BioEmu2 was slightly ahead at 1.5 Å in the study and ood60 systems. At 2.0 Å, BioEmu-1 had the higher budget-averaged recovery in the study systems (difference −0.038), the domain-motion benchmark systems (−0.094) and the ood60 systems (−0.088). The gap was largest in the ood60 systems at 3.0 Å (0.543 against 0.300), and BioEmu2 did not reach every state in any of these five systems, whereas BioEmu-1 did so in two. Without AdK, the difference was −0.075 at 2.0 Å and −0.113 at 3.0 Å, against −0.064 and −0.107 with all 22 systems.

At 500 samples, the expected numbers of states recovered within 3.0 Å for BioEmu-1, BioEmu2 and ColabFold were 20.3, 19.0 and 20 in the 11 study systems (27 states), 11.4, 11.0 and 11 in the six domain-motion systems (12 states) and 5.1, 3.0 and 5 in the five ood60 systems (10 states). Within 2.0 Å, BioEmu-1 recovered more states than ColabFold in 500 samples (28.9 against 25, with 25.4 for BioEmu2), and most of this advantage came from the 11 benchmark systems (14.2 against 11, with 12.5 for BioEmu2), whereas the study systems gave 14.6, 12.9 and 14. ColabFold reached every state within 3.0 Å in 11 of the 22 systems, against an expected 12.7 for BioEmu-1 and 10.0 for BioEmu2 in 500 samples, estimated from 200 random subsets per run. With 10,000 samples per run, the number of states recovered within 3.0 Å rose to 40.7 for BioEmu-1 but stayed at 33.0 for BioEmu2.

For the binary outcome at 3.0 Å with the two-of-three rule, leaving out one family at a time gave exact McNemar *p* values from 0.0625 to 0.25, and in none of these comparisons did BioEmu2 reach every state of a system in which BioEmu-1 did not. The largest *p* value came from leaving out the five kinases. Without CypA, which both models reached, the BioEmu-1, BioEmu2 and Both counts of Table 2 each fall by one, and the *p* values are unchanged.

To test whether the states that only BioEmu-1 reached depended on samples with poor local geometry, we counted the states again using only samples that passed the HIGH_QUALITY check. The counts used the subsets of Table S7, namely 1,000 random filter-passing samples per run, with the side chains of the BioEmu-1 samples rebuilt with HPacker. For BioEmu-1, only the samples that passed the check were counted, which left between 25 (SHP2) and 2,451 (T4L L99A) of the 3,000 checked samples per system, with a median of 701.5. For BioEmu2, all 3,000 checked samples of each system were counted, as 65,998 of its 66,000 checked samples passed. As in the binary outcome, a state was counted as reached when a counted sample lay within 3.0 Å of it in at least two of the three runs.

With only its passing samples, BioEmu-1 reached 36 of the 49 states and every state in 12 systems, against 33 states and 10 systems for BioEmu2. With all of its checked samples, BioEmu-1 reached 38 states and every state in 14 systems, so the check removed two states, namely the inactive state of ERK2 and the DFG-in inactive loop of p38α. Five states were reached only by the passing BioEmu-1 samples, namely the c-di-GMP-bound state of the LapD module, the active-like loop of p38α, the DNA-bound state of NanR, the active state of ERK2 and the apo state of Pol β. Two states were reached only by BioEmu2, namely the DFG-in inactive loop of p38α and the closed state of SHP2, for which BioEmu-1 kept only 25 passing samples.

These five states are among the 10 states that only BioEmu-1 reached in the full runs (Table 3 of the main text). For each of them, the share of nearby samples that passed the check was similar to or higher than the share among all checked samples of the same system. The subsets were too small to test the other five states. Only one to three checked samples lay within 3.0 Å of the DNA-bound a state of BusR and of the inactive state of ERK2, and none of them passed the check. No checked sample lay within 3.0 Å of the NMR intermediate of SRC, the sialic-acid-bound state of NanR or the inactive, folded loop of Aurora A, which were reached in only one or two runs of the full data. These results suggest that the lead of BioEmu-1 in state recovery does not come mainly from samples with poor local geometry. Testing the rarest states would need the check, or a relaxation, of every sample near them.

#### S2.4 Domain arrangements and comparison with SAXS

For Pol β, BioEmu-1 placed 29 of its 28,485 filter-passing samples within 3.0 Å Cα RMSD of the extended apo structure (PDB ID: 1BPD), whereas none of the BioEmu2 samples came closer than 3.19 Å. Over the scored residues, the Cα Rg was 28.1 Å for the apo structure and 22.7 Å for the DNA complex (PDB ID: 2BPG). The median Cα Rg of the filter-passing samples was 22.3 Å for BioEmu-1 and 21.7 Å for BioEmu2, and only 8.0% and 0.4% of their samples were closer in Rg to the apo structure than to the DNA complex. Comparing these Cα values with the measured scattering Rg is approximate, because it ignores the side chains and the hydration shell. However, the Cα Rg of the apo structure differs from its computed scattering Rg (28.9 Å) by only about 0.8 Å, which is much smaller than the 6 to 7 Å gap between the samples and the measured value.

In NanR, BioEmu2 packed the DNA-binding domain against the C-terminal domain, 26.6 Å and 77° from its position in the sialic-acid-bound structure. In BusR, it brought the two end domains into contact in 97.4% of its samples, compared with 0.2% for BioEmu-1. The long helix that connects them was intact in only 44% of the BioEmu2 samples, against 93% of the BioEmu-1 samples and 77% to 100% in the references. As a result, the BioEmu2 BusR chains had an Rg of 18.0 Å, compared with 36.1 and 43.2 Å for the references and 37.2 Å for BioEmu-1. Solution data cannot test these single chains directly, because free BusR is a tetramer (Rg 44 Å)[16] and free NanR is a dimer (Rg 32.3 Å).[17] However, every chain of the apo BusR tetramer (PDB ID: 7B5T) is extended (Cα Rg 38 to 44 Å), and the BioEmu2 NanR chains (median Cα Rg 18.9 Å over residues 30 to 244) were more compact than the chain in a dimer model fitted to the NanR scattering data (26.1 Å). With DNA and cyclic di-AMP, the median Cα Rg of BusR over all 215 residues rose only from 17.9 to 19.7 Å, and that of NanR was 19.1 Å with DNA and 19.3 Å with sialic acid.

With DNA and cyclic di-AMP, the share of BusR chains with a Cα Rg above 30 Å rose from 0.06% to 6.2%, but the median distances to the two DNA-bound states did not decrease (22.1 and 25.6 Å, against 21.0 and 23.8 Å without the partner). With pY530, the centres of the SH2 and kinase domains of SRC lay within 2.0 Å of their inactive crystal distance in 97.1% to 99.6% of samples, so the remaining difference from the inactive state appears to lie mainly within the kinase domain. We did not check whether the phosphotyrosine was bound in the SH2 pocket, and all three inactive crystal structures contain an ATP-site ligand that these runs omitted.

The domains and their placement (Figure 6) were defined as follows. To find the rigid domains of Pol β, NanR and BusR, we computed the standard deviation of every Cα–Cα distance over 1,500 random filter-passing samples of each model, grouped the residues by average-linkage clustering of these deviations, and kept groups of at least 25 residues as domains, so that a domain is a set of residues that move together in the samples of both models. Each sample was then superposed on the largest domain of a reference structure. The shift of another domain is the distance between the centres of its Cα atoms in the sample and in the reference, and its rotation is the angle of the further rotation that superposes it on its position in the reference. The internal RMSD of a domain was computed after superposing that domain alone.

#### S2.5 Pairwise RMSD and cluster count

**Table S9.** Pairwise RMSD and cluster count of each ensemble, per system and method.

| System | Method | Samples per run | Pairwise RMSD of scored residues, median (IQR) (Å) | Pairwise RMSD of alignment residues, median (Å) | Clusters at 3.0 Å | Clusters holding ≥ 1% of samples | Clusters in 500 samples |
|---|---|---|---|---|---|---|---|
| ABL1 | BioEmu-1 | 1000 | 3.682 (3.002 to 4.628) | 0.923 | 138.333 | 6.667 | 81.667 |
| ABL1 | BioEmu2 | 1000 | 2.475 (1.977 to 3.237) | 0.804 | 11.333 | 3.333 | 10 |
| ABL1 | ColabFold | 500 | 0.5 (0.368 to 0.653) | 0.137 | 2 | 1 | 2 |
| ABL1 | All-atom MD | 500 | 2.571 (2.033 to 3.133) | 1.159 | 4.5 | 2.5 | 4.5 |
| Aurora A | BioEmu-1 | 1000 | 5.252 (3.406 to 8.107) | 0.605 | 278.667 | 7.667 | 166 |
| Aurora A | BioEmu2 | 1000 | 3.662 (2.964 to 4.664) | 0.81 | 104.667 | 9.333 | 69 |
| Aurora A | ColabFold | 500 | 1.206 (0.884 to 1.619) | 0.137 | 3 | 1 | 3 |
| p38α | BioEmu-1 | 1000 | 4.903 (3.717 to 6.487) | 0.502 | 169 | 12.667 | 102.333 |
| p38α | BioEmu2 | 1000 | 3.622 (2.921 to 4.43) | 1.163 | 32.333 | 7 | 22.333 |
| p38α | ColabFold | 499 | 1.51 (0.965 to 2.597) | 0.221 | 3 | 2 | 3 |
| ERK2 | BioEmu-1 | 1000 | 4.358 (3.585 to 5.405) | 1.087 | 203.333 | 12 | 128 |
| ERK2 | BioEmu2 | 1000 | 2.529 (2.124 to 2.983) | 0.864 | 12.667 | 2 | 9.333 |
| ERK2 | ColabFold | 498 | 1.879 (1.198 to 2.407) | 0.473 | 2 | 1 | 2 |
| SRC | BioEmu-1 | 1000 | 2.24 (1.499 to 3.497) | 1.112 | 43 | 3.333 | 26.333 |
| SRC | BioEmu2 | 1000 | 2.09 (1.669 to 2.528) | 1.193 | 5 | 1 | 3.667 |
| SRC | ColabFold | 498 | 0.374 (0.302 to 0.452) | 0.213 | 1 | 1 | 1 |
| PTP1B | BioEmu-1 | 1000 | 1.855 (1.241 to 2.849) | 1.016 | 18.667 | 2 | 11.333 |
| PTP1B | BioEmu2 | 1000 | 1.18 (0.946 to 1.519) | 0.874 | 1 | 1 | 1 |
| PTP1B | ColabFold | 499 | 0.604 (0.364 to 0.947) | 0.291 | 1 | 1 | 1 |
| PTP1B | All-atom MD | 500 | 1.291 (0.956 to 1.675) | 1.079 | 1.167 | 1 | 1.167 |
| SHP2 | BioEmu-1 | 1000 | 30.322 (13.341 to 55.504) | 0.656 | 610.667 | 10.333 | 337.667 |
| SHP2 | BioEmu2 | 1000 | 2.817 (2.151 to 3.729) | 1.067 | 7 | 3 | 5.667 |
| SHP2 | ColabFold | 489 | 40.27 (22.107 to 55.403) | 0.445 | 291 | 15 | 291 |
| MBP | BioEmu-1 | 1000 | 1.656 (1.287 to 2.335) | 1.656 | 12.333 | 2.667 | 11 |
| MBP | BioEmu2 | 1000 | 1.533 (1.25 to 1.974) | 1.533 | 1 | 1 | 1 |
| MBP | ColabFold | 500 | 1.456 (0.345 to 3.004) | 1.456 | 1 | 1 | 1 |
| T4L L99A | BioEmu-1 | 1000 | 1.294 (0.895 to 1.951) | 1.514 | 15 | 2 | 7.667 |
| T4L L99A | BioEmu2 | 1000 | 0.76 (0.631 to 0.941) | 0.859 | 1 | 1 | 1 |
| T4L L99A | ColabFold | 500 | 0.384 (0.228 | 0.424 | 1 | 1 | 1 |

| System | Method | Samples per run | Pairwise RMSD of scored residues, median (IQR) (Å) | Pairwise RMSD of alignment residues, median (Å) | Clusters at 3.0 Å | Clusters holding ≥ 1% of samples | Clusters in 500 samples |
|---|---|---|---|---|---|---|---|
| | | | to 0.665) | | | | |
| T4L L99A | All-atom MD | 500 | 1.44 (1.079 to 1.919) | 1.58 | 1.167 | 1 | 1.167 |
| CypA | BioEmu-1 | 1000 | 1.329 (1.1 to 1.648) | 1.329 | 1 | 1 | 1 |
| CypA | BioEmu2 | 1000 | 0.692 (0.64 to 0.751) | 0.692 | 1 | 1 | 1 |
| CypA | ColabFold | 499 | 0.195 (0.158 to 0.237) | 0.195 | 1 | 1 | 1 |
| CypA | All-atom MD | 500 | 0.857 (0.777 to 0.944) | 0.857 | 1 | 1 | 1 |
| AdK | BioEmu-1 | 1000 | 3.431 (2.669 to 4.671) | 3.431 | 149.667 | 5 | 83.333 |
| AdK | BioEmu2 | 1000 | 3.054 (2.35 to 4.094) | 3.054 | 16.667 | 4.667 | 15.333 |
| AdK | ColabFold | 500 | 1.525 (0.801 to 2.264) | 1.525 | 1 | 1 | 1 |
| AdK | All-atom MD | 500 | 2.672 (2.092 to 3.417) | 2.672 | 4.833 | 3.333 | 4.833 |
| RbsB | BioEmu-1 | 1000 | 2.411 (1.666 to 3.977) | 2.411 | 56.333 | 4.667 | 33.667 |
| RbsB | BioEmu2 | 1000 | 2.26 (1.267 to 3.582) | 2.26 | 3 | 2.333 | 2.667 |
| RbsB | ColabFold | 500 | 0.945 (0.444 to 1.898) | 0.945 | 1 | 1 | 1 |
| DppA | BioEmu-1 | 1000 | 3.254 (1.711 to 7.049) | 3.254 | 101.333 | 6.333 | 59 |
| DppA | BioEmu2 | 1000 | 2.707 (1.839 to 3.913) | 2.707 | 6.667 | 3 | 5.333 |
| DppA | ColabFold | 500 | 0.871 (0.545 to 1.413) | 0.871 | 1 | 1 | 1 |
| Hexokinase | BioEmu-1 | 1000 | 2.101 (1.425 to 3.398) | 2.101 | 26.667 | 3 | 18.667 |
| Hexokinase | BioEmu2 | 1000 | 1.573 (1.399 to 1.796) | 1.573 | 1 | 1 | 1 |
| Hexokinase | ColabFold | 499 | 0.666 (0.44 to 0.945) | 0.666 | 1 | 1 | 1 |
| β-PGM | BioEmu-1 | 1000 | 1.711 (1.313 to 2.413) | 1.711 | 24.667 | 1.667 | 13.333 |
| β-PGM | BioEmu2 | 1000 | 1.432 (1.176 to 1.788) | 1.432 | 1.333 | 1 | 1 |
| β-PGM | ColabFold | 500 | 1.522 (0.609 to 1.963) | 1.522 | 1 | 1 | 1 |
| Pol β | BioEmu-1 | 1000 | 3.055 (1.845 to 6.432) | 3.055 | 113.667 | 5 | 72 |
| Pol β | BioEmu2 | 1000 | 3.1 (2.441 to 3.983) | 3.1 | 28 | 4.667 | 19.333 |
| Pol β | ColabFold | 500 | 0.911 (0.692 to 1.166) | 0.911 | 1 | 1 | 1 |
| Biotin carboxylase | BioEmu-1 | 1000 | 1.134 (0.872 to 1.515) | 1.134 | 2.667 | 1 | 1.667 |
| Biotin carboxylase | BioEmu2 | 1000 | 1.552 (1.322 to 1.907) | 1.552 | 2.667 | 1 | 2 |
| Biotin carboxylase | ColabFold | 497 | 0.674 (0.509 to 1.002) | 0.674 | 1 | 1 | 1 |
| LapD module | BioEmu-1 | 1000 | 10.863 (5.162 to 15.08) | 10.863 | 272 | 10.333 | 164.667 |
| LapD module | BioEmu2 | 1000 | 1.957 (1.696 to 2.312) | 1.957 | 2.333 | 1 | 1.667 |
| LapD module | ColabFold | 499 | 3.789 (2.144 to 9.053) | 3.789 | 15 | 7 | 15 |
| EcoT38I | BioEmu-1 | 1000 | 4.863 (3.119 | 4.863 | 141.333 | 9 | 92.667 |

| System | Method | Samples per run | Pairwise RMSD of scored residues, median (IQR) (Å) | Pairwise RMSD of alignment residues, median (Å) | Clusters at 3.0 Å | Clusters holding ≥ 1% of samples | Clusters in 500 samples |
|---|---|---|---|---|---|---|---|
| | | | to 8.325) | | | | |
| EcoT38I | BioEmu2 | 1000 | 4.871 (3.465 to 6.877) | 4.871 | 79 | 11.333 | 54 |
| EcoT38I | ColabFold | 500 | 1.291 (0.886 to 2.288) | 1.291 | 1 | 1 | 1 |
| NanR | BioEmu-1 | 1000 | 2.808 (1.971 to 4.197) | 2.808 | 69.333 | 5.333 | 48.667 |
| NanR | BioEmu2 | 1000 | 8.563 (6.238 to 11.266) | 8.563 | 529.667 | 13.667 | 309.667 |
| NanR | ColabFold | 500 | 2.256 (1.518 to 3.106) | 2.256 | 4 | 2 | 4 |
| BusR | BioEmu-1 | 1000 | 10.467 (8.151 to 12.924) | 12.023 | 652.333 | 7 | 374 |
| BusR | BioEmu2 | 1000 | 15.878 (13.624 to 18.31) | 14.186 | 1000 | 0 | 500 |
| BusR | ColabFold | 498 | 3.095 (2.012 to 4.389) | 4.952 | 9 | 4 | 9 |
| Transposase | BioEmu-1 | 1000 | 2.005 (1.648 to 2.488) | 2.005 | 18 | 1.333 | 11.333 |
| Transposase | BioEmu2 | 1000 | 2.488 (2.193 to 2.849) | 2.488 | 6.333 | 1.667 | 5 |
| Transposase | ColabFold | 499 | 0.849 (0.716 to 1.002) | 0.849 | 1 | 1 | 1 |

For each run, every pair among 1,000 random filter-passing samples (all filter-passing ColabFold models, and 500 frames per all-atom replica after the first 10 ns) was superposed on the alignment residues, and the Cα RMSD was computed over the scored residues. Cluster counts use the method of Daura et al.[18] at a 3.0 Å cut-off on the same RMSD. Values are means over runs (replicas for all-atom MD). Over the 22 systems, the median pairwise RMSD of BioEmu2 was smaller than that of BioEmu-1 in 16 systems (median ratio 0.83, exact sign-flip test on the 14 family means of the log ratio, $p$ = 0.13). Because small clusters dominate the cluster count, the number of clusters holding at least 1% of the samples and the count on a random 500 samples per run, for comparison with the 500 ColabFold models, are also given. IQR, interquartile range.

The reference spread of BioEmu2 was at least that of BioEmu-1 in five systems: NanR (ratio 2.51), RbsB (1.24), AdK (1.08), the transposase (1.03) and MBP (1.00). In each of the five kinases, the pairwise RMSD of BioEmu2 lay between those of the other two methods (Table S9) (2.24 to 5.25 Å for BioEmu-1, 2.09 to 3.66 Å for BioEmu2 and 0.37 to 1.88 Å for ColabFold).

#### S2.6 Populations

The populations before renormalisation are given in Table S10. At the 1.5 Å class threshold of PTP1B, 9.3% of the BioEmu2 samples and 38.9% of the BioEmu-1 samples were unassigned, so the closed share before renormalisation was 2.07% and 43.8%, respectively. In all-atom MD, the PTP1B WPD loop opened in all three replicas started from the closed structure, but these replicas were still closed in 70.8%, 98.3% and 0.1% of their assigned frames, and no replica started from the open structure closed, so MD gives no population (Table S15).

The class whose population differed most between the two models kept the sign of its difference at every class threshold tested in 19 of the 21 systems. In ABL1, the difference was −0.02 pp at 2.0 Å but +18 to +54 pp at the other thresholds. In SRC, it was −31 pp at 2.5 Å and +12 pp at 3.5 Å, so the relative preference of the two models in SRC depends on the threshold. When SRC was rescored with the activation loop excluded from the superposition, the BioEmu2 population of the αC-in state fell from 44.1% to 21.0%, although this did not change which states were reached. The Asp181–Cys215 distance of PTP1B needs only Cα atoms, so every method could be scored on it. It leaves a gap of 3.83 Å between the closed (8.29 to 8.45 Å) and open (12.28 to 12.96 Å) crystal structures, so the cut-off could be placed anywhere in that gap, and we therefore tested nine cut-offs across it, from 8.689 to 12.0 Å (Table S11, Figure S2a and b).

**Table S10.** Populations of the measured states outside ABL1 before renormalisation, with the unassigned fraction, for each method.

| System · state | Method | Before renormalisation (%) | Unassigned (%) | Renormalised (%) |
|---|---|---|---|---|
| PTP1B · WPD closed | BioEmu-1 | 43.83 (42.98–44.28) | 38.93 | 71.76 |
| PTP1B · WPD closed | BioEmu2 | 2.07 (1.86–2.26) | 9.26 | 2.29 |
| PTP1B · WPD closed | ColabFold | 85.57 | 10.82 | 95.96 |
| SHP2 · open | BioEmu-1 | 98.52 (98.45–98.59) | 0.00 | 98.52 |
| SHP2 · open | BioEmu2 | 0.00 | 0.00 | 0.00 |
| SHP2 · open | ColabFold | 93.66 | 0.00 | 93.66 |
| Aurora A · loop closed | BioEmu-1 | 1.20 (1.14–1.28) | 0.00 (not determined: 0.00) | 1.20 |
| Aurora A · loop closed | BioEmu2 | 0.05 (0.02–0.09) | 0.00 | 0.05 |
| Aurora A · loop closed | ColabFold | 0.20 | 0.00 | 0.20 |
| MBP · sugar-bound closed | BioEmu-1 | 72.41 (72.11–72.79) | 27.09 | Not renormalised |
| MBP · sugar-bound closed | BioEmu2 | 1.22 (1.04–1.42) | 36.65 | Not renormalised |
| MBP · sugar-bound closed | ColabFold | 54.40 | 5.00 | Not renormalised |
| MBP · partially closed | BioEmu-1 | 0.003 (0.000–0.010) | 27.09 | Not renormalised |
| MBP · partially closed | BioEmu2 | 0.00 | 36.65 | Not renormalised |
| MBP · partially closed | ColabFold | 0.00 | 5.00 | Not renormalised |
| T4L L99A · excited (backbone class) | BioEmu-1 | 0.024 (0.010–0.050) | 22.58 | Not renormalised |
| T4L L99A · excited (backbone class) | BioEmu2 | 0.00 | 1.18 | Not renormalised |
| T4L L99A · excited (backbone class) | ColabFold | 0.00 | 0.00 | Not renormalised |
| CypA · Phe113 minor rotamer | BioEmu-1 | Not measurable from its own output | | |
| CypA · Phe113 minor rotamer | BioEmu2 | 7.88 (7.53–8.10) | 0.03 | Not renormalised |
| CypA · Phe113 minor rotamer | ColabFold | 66.53 | 0.00 | Not renormalised |

Values are percentages of filter-passing samples, mean over runs with the range over runs in brackets, at the class level used in Table 5 (backbone classes for PTP1B, MBP and T4L L99A, and classes from the interface distance, loop distance or side-chain rotamer for SHP2, Aurora A and CypA). The renormalised value is the population over the two measured states, as given in Table 5. All-atom MD is not listed, because its replicas stayed near their start state, so a value pooled over replicas is not a population. Table S15 gives its occupancy for each start state. For Aurora A, BioEmu-1 samples outside the two loop classes that BioEmu-1 can compute would be counted as not determined rather than unassigned, and there were none. SHP2 and Aurora A did not have any unassigned samples, because their two classes cover every filter-passing sample. For T4L L99A, the class that also requires the Phe114 trans rotamer gave the same values as the backbone class for BioEmu2 and ColabFold. BioEmu-1 does not write any side chains, so its CypA rotamer is not measurable from its own output. The value after side-chain rebuilding is in Table 5.

**Table S11.** Closed WPD-loop population of PTP1B on the Cα–Cα distance between Asp181 and Cys215.

| Cut-off (Å) | Note | BioEmu-1 | BioEmu2 | ColabFold |
|---|---|---|---|---|
| 8.689 | Closed bound of Yeh et al.[19] | 24.8% (23.6 to 25.4) (0/3) | 0.49% (0.47 to 0.50) (3/3) | 0% (1/1) |
| 9.0 | | 34.2% (33.2 to 35.3) (0/3) | 1.05% (0.98 to 1.10) (3/3) | 10.0% (0/1) |
| 9.5 | | 46.0% (45.4 to 46.4) (0/3) | 2.68% (2.57 to 2.80) (3/3) | 29.3% (0/1) |
| 9.87 | Cut-off used in the Results of Yeh et al.[19] | 52.2% (51.3 to 52.8) (0/3) | 4.43% (4.24 to 4.67) (3/3) | 48.7% (0/1) |
| 10.075 | Open bound of Yeh et al.[19] | 55.0% (54.4 to 55.5) (0/3) | 5.54% (5.42 to 5.74) (0/3) | 59.3% (0/1) |
| 10.5 | Midpoint of the closed and open structures (secondary boundary defined before sampling) | 60.3% (59.7 to 60.7) (0/3) | 8.46% (8.19 to 8.67) (0/3) | 82.2% (0/1) |
| 11.0 | | 65.8% (65.4 to 66.3) (0/3) | 14.1% (13.7 to 14.4) (0/3) | 97.4% (0/1) |
| 11.5 | | 71.1% (70.4 to 71.8) (0/3) | 27.3% (27.1 to 27.6) (0/3) | 100% (0/1) |
| 12.0 | Below the smallest open value (12.28 Å) | 75.9% (75.0 to 76.6) (0/3) | 51.6% (51.0 to 52.1) (0/3) | 100% (0/1) |

A filter-passing sample counts as closed when this distance is below the cut-off. Values are the mean over runs (range over runs), with the number of runs within the tolerance of 0% to 5.3% in brackets. ColabFold has one set of 500 models. The median distance, with the middle half of samples in brackets, was 9.73 Å (8.70 to 11.91) for BioEmu-1, 11.97 Å (11.44 to 12.47) for BioEmu2 and 9.88 Å (9.40 to 10.36) for ColabFold. Under the two-boundary rule of Yeh et al.[19] (closed below 8.689 Å, open above 10.075 Å), 30.3% of the BioEmu-1 samples, 5.05% of the BioEmu2 samples and 59.3% of the ColabFold models lay between the two boundaries.

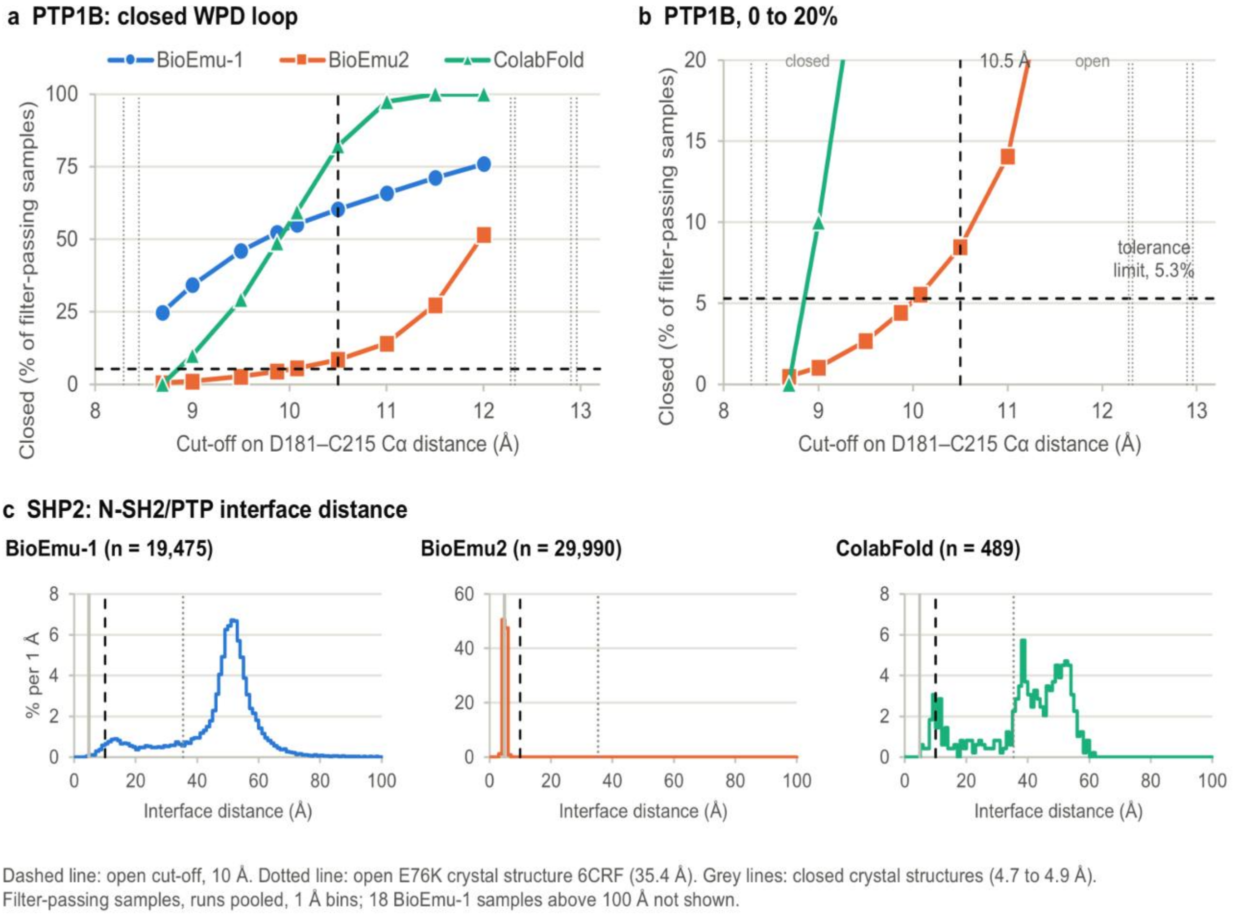


**Figure S2. Dependence of two population comparisons on the definition of the states.** (a) Closed WPD-loop population of PTP1B against the cut-off on the Cα–Cα distance between Asp181 and Cys215 for BioEmu-1, BioEmu2 and ColabFold (mean over runs). The dashed horizontal line is the upper limit of the tolerance of the measured 2.5 ± 1.4% (5.3%), the dashed vertical line marks the secondary boundary of 10.5 Å and dotted lines mark the closed and open crystal structures. (b) The same data with the population axis from 0% to 20%. (c) Distribution of the SHP2 N-SH2/PTP interface distance for each method (filter-passing samples, runs pooled, 1 Å bins). The dashed line marks the open cut-off of 10 Å, the dotted line marks the crystal structure of the open E76K mutant (PDB ID: 6CRF) at 35.4 Å and the grey lines mark the closed crystal structures (4.7 to 4.9 Å). Eighteen BioEmu-1 samples above 100 Å are not shown.

For SHP2, because "open" here means only that the interface distance exceeds 10 Å, we checked the structure of the open BioEmu-1 samples (Table S12). In 94.8% of them, all three domains were within 2.0 Å Cα RMSD of a crystal structure, so in most samples the domains kept their fold. However, in 88.2% the N-SH2 domain packed against the C-terminal SH2 domain, a contact that none of the five crystal structures shows. Whether this arrangement resembles the released, mobile N-SH2 domain inferred from NMR cannot be decided from these data.

**Table S12.** SHP2 N-SH2/PTP interface distance for each method and the structure of the open samples.

| Quantity | BioEmu-1 | BioEmu2 | ColabFold |
|---|---|---|---|
| Filter-passing samples | 19,475 | 29,990 | 489 |
| Interface distance, median (middle half), Å | 50.4 (44.0 to 54.4) | 4.98 (4.69 to 5.26) | 42.6 (35.3 to 50.4) |
| Interface distance, 5th to 95th percentile, Å | 14.4 to 63.4 | 4.24 to 5.66 | 9.5 to 55.2 |
| Open at 8 / 10 / 15 / 20 / 35 Å cut-off | 99.5 / 98.5 / 94.6 / 91.5 / 83.6% | 0% at all | 98.6 / 93.7 / 86.9 / 84.7 / 75.5% |
| Open samples (interface distance above 10 Å) | 19,186 | 0 | 458 |
| All three domains within 2.0 Å of a crystal structure | 94.8% | (closed: 98.2%) | 100% |
| Shift of the N-SH2 centre from its closed position, median (middle half), Å | 56.6 (51.2 to 58.9) | (closed: 2.1) | 47.4 (32.9 to 59.5) |
| Rotation of N-SH2, median (middle half), degrees | 131 (117 to 147) | (closed: 10.5) | 106 (86 to 127) |
| No N-SH2 Cα within 8 Å of the PTP domain | 42.1% | (closed: 0%) | 18.6% |
| N-SH2 in contact with C-SH2 | 88.2% | (closed: 42.3%) | 90.8% |
| N-SH2 RMSD to the 6CRF position after superposition on PTP, median (closest), Å | 27.7 (4.52) | (closed: 55.9) | 55.9 (11.8) |

The interface distance is the shortest Cα–Cα distance between N-SH2 residues 58 to 62 and PTP residues 459 to 465, and a sample is open when it exceeds the cut-off. The closed crystal structures (PDB IDs: 4DGP, 2SHP, 5EHR and 7JVM) have an interface distance of 4.69 to 4.93 Å, an N-SH2 shift of 0 to 1.51 Å, a rotation of 0° to 5.5°, 49 to 53 N-SH2/PTP Cα contacts and no N-SH2/C-SH2 contact. The E76K structure (PDB ID: 6CRF) has an interface distance of 35.4 Å, a shift of 56.4 Å, a rotation of 133°, three N-SH2/PTP Cα contacts and no N-SH2/C-SH2 contact. It lies 56.9 Å from the closed structures by N-SH2 RMSD. Contacts are Cα pairs closer than 8 Å. The domains are N-SH2 (residues 3 to 103), C-SH2 (112 to 216) and PTP (247 to 525). The fold of each domain is the Cα RMSD to the nearest crystal structure after superposition on that domain alone, and the pose is measured after superposition on the PTP domain of the closed structure (PDB ID: 4DGP). Among the open BioEmu-1 samples, the median fold RMSDs were 1.13 Å (N-SH2), 0.78 Å (C-SH2) and 1.01 Å (PTP). Values in brackets for BioEmu2 describe its closed samples, because it did not produce any open sample. BioEmu-1 also placed 289 filter-passing samples at 10 Å or below. In these, the N-SH2 domain was rotated by a median of 31°, and the C-SH2 domain was within 2.0 Å of a crystal structure in 79.2% of them.

For MBP, because the partially closed form was inferred from paramagnetic data rather than solved as a crystal structure, we also counted the BioEmu2 samples that were not open. Their share was 37.0% to 38.3% at a threshold of 1.5 Å but only 1.1% to 1.4% at 3.0 Å, and none of them lay within the 1.5 Å class threshold of the partially closed form (closest 2.12 Å), so they appear to be spread around the open state rather than gathered in a second conformation. Counted as not open, that is, more than 1.5 Å from every open structure, 99.5% of the BioEmu-1 samples, 37.9% of the BioEmu2 samples (1.2% at 3.0 Å) and 59.4% of the ColabFold models were not open.

For CypA, the Phe113 rotamer alone may not correspond to the minor state seen by NMR. ColabFold placed 66.5% of its models in the minor rotamer. The rebuilt BioEmu-1 samples placed Phe113 in the minor rotamer in 3.6% of cases (3.2% to 4.2% per run). When the side chains of BioEmu2 samples were rebuilt in the same way, HPacker kept the Phe113 rotamer that BioEmu2 had written in 99.5% of them, which suggests that the backbone largely fixes this rotamer, but the rebuild changed Ser99 in 24.1% of these samples, so the rebuilt BioEmu-1 share with both minor-state rotamers was not compared with the measured state.

For T4L L99A, the ground and excited structures are only 2.3 Å apart, so the backbone classes can overlap and the Phe114 rotamer is the more decisive test. After the rebuild, BioEmu-1 placed Phe114 in the trans rotamer in 0.3% of samples (0.1% to 0.4% per run), and HPacker kept the Phe114 rotamer of every rebuilt BioEmu2 sample. In all-atom MD, replicas started from the ground state never reached the excited state, and replicas started from the excited state kept only 0.2% to

9.6% of frames in its backbone class and did not reach the ground state, although Phe114 stayed trans in 90.9% to 100% of frames.

#### S2.7 ABL1

**Table S13.** ABL1 wild-type populations against the NMR values of Xie et al.[5]

| Method | Active | I1 | I2 | Runs within tolerance |
|---|---|---|---|---|
| Measured[5] ± tolerance | 0.880 ± 0.014 | 0.059 ± 0.002 | 0.061 ± 0.014 | |
| BioEmu-1 | 0.724 (0.719–0.734) | 0.271 (0.261–0.276) | 0.0049 (0.0045–0.0053) | 0 of 3 |
| BioEmu2 | 1.000 | 0 | 0 | 0 of 3 |
| ColabFold 32:64 | 1.000 | 0 | 0 | 0 of 1 |
| ColabFold 16:32 | 1.000 | 0 | 0 | 0 of 1 |
| ColabFold 8:16 | 0.983 | 0.017 | 0 | 0 of 1 |
| All-atom MD | Not a population: replicas started from the active state were 25.8% to 99.9% active, and replicas started from I2 were 77.4% to 99.0% I2 | | | |

Values are renormalised populations of the three states (mean over runs, with the range over runs in brackets), with one set for each ColabFold alignment depth. Tolerances are twice the reported standard uncertainty. Before renormalisation, BioEmu-1 placed 28.3% of its filter-passing samples in the active state, 10.6% in I1 and 0.19% in I2, and left 60.9% unassigned. Of the unassigned BioEmu-1 samples, 99.3% failed the class rule that the ψ angle of A399 lies in the β region (|ψ| of at least 90°). BioEmu2 placed 99.94% in the active state and left 0.06% unassigned.

**Table S14.** ABL1 responses to perturbation for BioEmu-1 and BioEmu2.

| Condition | Comparison | Method | State | Comparison (%) | Condition (%) | Model direction | Measured direction | Agreement | Apparent ΔΔG, model | ΔΔG, measured |
|---|---|---|---|---|---|---|---|---|---|---|
| T408Y | WT | BioEmu-1 | I2 | 0.191 | 0.213 | No detectable change | Up | No detectable change | −0.06 | −1.45 |
| T408Y | WT | BioEmu2 | I2 | 0.000 | 0.000 | State absent | Up | Cannot respond (state absent) | Undefined | −1.45 |
| Y272H | WT | BioEmu-1 | I2 | 0.191 | 0.199 | No detectable change | Down | No detectable change | −0.02 | ≥ 1.07 |
| Y272H | WT | BioEmu2 | I2 | 0.000 | 0.000 | State absent | Down | Cannot respond (state absent) | Undefined | ≥ 1.07 |
| F378V | WT | BioEmu-1 | I2 | 0.191 | 0.125 | Down | Down | Same | 0.25 | ≥ 1.07 |
| F378V | WT | BioEmu2 | I2 | 0.000 | 0.000 | State absent | Down | Cannot respond (state absent) | Undefined | ≥ 1.07 |
| Y272H/T408Y | T408Y | BioEmu-1 | I2 | 0.213 | 0.217 | No detectable change | Down | No detectable change | −0.01 | 1.15 |
| Y272H/T408Y | T408Y | BioEmu2 | I2 | 0.000 | 0.000 | State absent | Down | Cannot respond (state absent) | Undefined | 1.15 |
| F378V/T408Y | T408Y | BioEmu-1 | I2 | 0.213 | 0.161 | Down | Down | Same | 0.17 | 0.74 |
| F378V/T408Y | T408Y | BioEmu2 | I2 | 0.000 | 0.000 | State absent | Down | Cannot respond (state absent) | Undefined | 0.74 |
| H415P | WT | BioEmu-1 | I1 | 10.577 | 10.984 | No detectable change | Up | No detectable change | −0.02 | −0.43 |
| H415P | WT | BioEmu2 | I1 | 0.000 | 0.007 | No detectable change | Up | No detectable change | ≤ 0.24 | −0.43 |
| H415P | WT | BioEmu-1 | I2 | 0.191 | 0.095 | Down | Down | Same | 0.41 | |
| H415P | WT | BioEmu2 | I2 | 0.000 | 0.000 | State absent | Down | Cannot respond (state absent) | Undefined | |
| pY412 | WT | BioEmu2 | I1 | 0.000 | 0.007 | No detectable change | Down | No detectable change | ≤ 0.24 | |
| pY412 | WT | BioEmu2 | I2 | 0.000 | 0.000 | State absent | Down | Cannot respond (state absent) | Undefined | |
| Y412E | WT | BioEmu-1 | I2 | 0.191 | 1.124 | Up | Down | Opposite | −1.05 | |
| SH3-SH2 construct | WT | BioEmu2 | I2 | 0.000 | 0.000 | State absent | Up | Cannot respond (state absent) | Undefined | −1.03 |

| Condition | Comparison | Method | State | Comparison (%) | Condition (%) | Model direction | Measured direction | Agreement | Apparent ΔΔG, model | ΔΔG, measured |
|---|---|---|---|---|---|---|---|---|---|---|
| GNF-5 | SH3-SH2 construct | BioEmu2 | I2 | 0.000 | 0.000 | State absent | Up | Cannot respond (state absent) | Undefined | −0.61 |
| Imatinib | WT | BioEmu2 | I2 | 0.000 | 8.757 | Up | Expected: most drug-bound molecules in I2 (closed activation loop, DFG out) | | ≤ −4.01 | |
| Dasatinib | WT | BioEmu2 | I2 | 0.000 | 0.000 | State absent | Expected: most drug-bound molecules active | | Undefined | |

Populations are percentages of filter-passing samples (mean over three runs) of the state named, in the comparison condition and in the perturbed condition. A change was called only when the three run values of the perturbed condition lay entirely above or entirely below those of its comparison. The apparent $\Delta\Delta G = -RT \ln(p_1/p_0)$ at 298.15 K, where $p_1$ and $p_0$ are the populations in the perturbed condition and in its comparison, is given in kcal/mol. Measured directions and ΔΔG values are from Xie et al.[5] For imatinib and dasatinib, the direction is the expected binding state of the drug rather than a measured population shift. "Undefined" means that the state was absent in both conditions. The symbols ≤ and ≥ give bounds from the exact one-sided 95% upper bound where a state was absent in one condition, or from a measured bound. BioEmu-1 was run for the seven conditions without ligands or partner domains, with Y412E as a phosphomimetic in place of pY412. WT, wild type.

For BioEmu2, H415P raises I1 from 5.9% to 12.1% in solution (ΔΔG −0.43 kcal/mol), but BioEmu2 placed at most 0.010% of samples in I1 in any run with H415P, as it did with pY412, so neither change was detectable (Table S14). Where the BioEmu-1 direction agreed with the measured one, the apparent ΔΔG was 0.17 to 0.41 kcal/mol, against measured values of 0.74 kcal/mol for F378V/T408Y and at least 1.07 kcal/mol for F378V.

In the propagation test, starting from BioEmu2's own wild-type samples allowed only active starts, and all 1,280 endpoints from these 10 starts and all 5,005 frames of five 1,000-step trajectories stayed active. From the NMR starts, the I1-start endpoints at 1,000 ns were 96 active, 219 I1 and five unassigned, and the I2-start endpoints were 21 active, 11 I1, 272 I2 and 16 unassigned. The endpoints also drifted from their references: only 58 of the 320 I2-start endpoints lay within 3.0 Å of I2 at 1,000 ns, against 190 at 100 ns. The lag is the nominal lag of the model, not a measured time.

Of the 5,005 chain frames per state from the NMR starts, the active-start chains had 5,004 active frames and one that failed the backbone filter; the I1-start chains had 5,001 I1 and 4 unassigned frames; and the I2-start chains had 3,606 I2, 1,369 I1, 28 unassigned and 2 failing frames. No frame of an I1-start or I2-start chain was active. The I1 frames of the I2-start chains came from three chains whose activation loop opened past the 22 Å cut-off while the DFG motif stayed flipped out; these frames lay at least 5.83 Å from the I1 structure, and two of the three chains ended in the I1 class. Chains and single jumps from the same five starts were compared at each lag, using all chain frames in a window around the lag (8 to 12, 80 to 120 and 900 to 1,000 ns) and the 160 single-jump endpoints. The fraction within 3.0 Å of the start structure, for chains and jumps respectively, was 0.32 and 0.76 at 10 ns, 0.00 and 0.24 at 100 ns and 0.16 and 0.09 at 1,000 ns for the I1 starts. For the I2 starts, it was 0.40 and 0.90, 0.00 and 0.60 and 0.00 and 0.23, and for the active starts 1.00 and 1.00, 0.88 and 0.98 and 0.75 and 0.79.

#### S2.8 Simulations

The all-atom occupancy of each state by replica is given in Table S15, and Table S16 gives, for each start state, the basin stability, the best progress towards the other state and the time of the first frame in the other state over all production frames. Replicas of AdK started from the open state stayed open, with 83.5% of frames open after the first 10 ns (73.1% to 90.4% per replica) and no frame closer than 3.95 Å to the closed state. Replicas started from the closed structure were closed in only 0.6% of frames after the first 10 ns (0% to 1.9% per replica). In CypA, Ser99 turned from its minor-state rotamer to trans during equilibration in the replicas started from the minor substate, so production did not start from the full minor substate. In T4L L99A, replicas started from the ground state came no closer than 1.74 to 1.91 Å to the excited state, and those started from the excited state left their basin but came no closer than 2.16 to 3.60 Å to the ground state.

**Table S15.** All-atom MD occupancy by start state.

| System | Start state | Occupancy, % (replicas 1, 2, 3) | WPD-closed share of the two states, % |
|---|---|---|---|
| T4L L99A | Ground | Ground 87.0, 30.1, 90.9; excited 0.0, 0.0, 0.0; unassigned 13.0, 69.9, 9.1 | – |
| T4L L99A | Excited | Ground 0.0, 0.0, 0.0; excited 0.2, 9.6, 9.0; unassigned 99.8, 90.4, 91.0 | – |
| CypA | Major (Phe113 χ1) | Major 95.8, 15.1, 98.4; minor 4.2, 84.9, 1.6; unassigned 0.0, 0.0, 0.0 | – |
| CypA | Minor (Phe113 χ1) | Major 21.8, 52.4, 35.2; minor 78.2, 47.5, 42.6; unassigned 0.0, 0.0, 22.2 | – |
| PTP1B | WPD open | WPD open 39.1, 60.6, 74.5; WPD closed 0.0, 0.0, 0.0; unassigned 60.9, 39.4, 25.5 | 0.0, 0.0, 0.0 |
| PTP1B | WPD closed | WPD open 7.5, 1.1, 33.8; WPD closed 18.0, 60.8, 0.0; unassigned 74.5, 38.1, 66.2 | 70.8, 98.3, 0.1 |
| AdK | Open | Open 86.9, 73.1, 90.4; closed 0.0, 0.0, 0.0; unassigned 13.1, 26.9, 9.6 | – |
| AdK | Closed | Open 76.7, 9.1, 34.1; closed 0.0, 0.0, 1.9; unassigned 23.3, 90.9, 64.0 | – |

Values are percentages of filter-passing frames from 10.1 to 250 ns (2,400 frames per replica) for replicas 1, 2 and 3 of each start state. For T4L L99A, PTP1B and AdK, a frame is assigned to a state when its Cα RMSD to that state is below the class threshold (1.5 Å for T4L L99A and PTP1B, 3.0 Å for AdK) and lower than its RMSD to the other state. Frames that meet this condition for none of the states are unassigned. For CypA, state assignment uses the Phe113 side-chain rotamer (χ1 angle near +60° for the major state and near −60° for the minor state), because its two backbone states are only 0.11 Å apart. For PTP1B, the last column gives the WPD-closed share of the two states, renormalised as in Table 5. ABL1 replicas are reported with the other ABL1 results.

**Table S16.** All-atom MD by start state over all production frames.

| Start state | Basin stability | Best progress | First frame in the other state (ns) |
|---|---|---|---|
| T4L L99A from ground | 0.71 (0.33 to 0.91) | 0.21 (0.17 to 0.24) | Never |
| T4L L99A from excited | 0.08 (0.02 to 0.13) | −0.34 (−0.56 to 0.06) | Never |
| CypA from major (Phe113 χ1) | 0.71 (0.18 to 0.98) | 1.00 | 54.6, 8.7, 31.0 |
| CypA from minor (Phe113 χ1) | 0.57 (0.45 to 0.79) | 1.00 | 50.7, 0.9, 112.4 |
| PTP1B from WPD open | 0.59 (0.40 to 0.75) | 0.47 (0.43 to 0.50) | Never |
| PTP1B from WPD closed | 0.27 (0.02 to 0.62) | 0.73 (0.70 to 0.79) | 73.6, 234.2, 8.8 |
| AdK from open | 0.84 (0.74 to 0.91) | 0.42 (0.38 to 0.45) | Never |
| AdK from closed | 0.03 (0.01 to 0.06) | 0.80 (0.77 to 0.84) | 12.4, 12.6, 48.2 |

Values are the mean (range) over three replicas, using all production frames (0.1 to 250 ns), because three replicas crossed to the other state within the first 10 ns. A frame is assigned to a backbone state when its Cα RMSD to that state is below the class threshold (1.5 Å for T4L L99A and PTP1B, 3.0 Å for AdK) and lower than its RMSD to the other state. Basin stability is the fraction of frames whose state label matches the start state. For T4L L99A, basin stability also requires the Phe114 rotamer of that state, and for PTP1B, the D181–C215 Cα distance of that state. Best progress is the highest value of 1 − RMSD(frame, other state)/*d*, where *d* is the smallest Cα RMSD between the reference structures of the two states, so it is 1 at the other state and negative when a frame lies farther from the other state than *d*. For CypA, whose two backbone states are 0.11 Å apart, progress between the states was measured on the Phe113 χ1 rotamer. The last column gives, for each replica, the time of the first frame in the other state.

#### S2.9 Software

**Table S17.** Software, versions and hardware used in this study.

| Component | Version or identifier |
|---|---|
| BioEmu2 | bioemu2-preview checkpoint (Hugging Face snapshot dd4b0c1b); code at repository commit 6d1ca744; CUDA 12 installation; Python 3.12; PyTorch 2.13.0; equilibrium mode, step scale 1.0, 200 denoising steps, batch size 32 |
| BioEmu-1 | bioemu 1.2.0 with its default checkpoint bioemu-v1.1; batch_size_100 set to 10; its own sample filter switched off |
| BioEmu-1 environment | PyTorch 2.13.0 built for CUDA 12.6; Python 3.11.18 |
| ColabFold | 1.5.4, the version bundled with bioemu 1.2.0; five AlphaFold2 models; 100 seeds; --max-msa 32:64 (ABL1 also 8:16 and 16:32) |
| Alignments | Public ColabFold MMseqs2 server (api.colabfold.com, the default of ColabFold 1.5.4), queried once per sequence on 4 and 5 November 2026 |
| Backbone filter | The filter shipped with bioemu 1.2.0 (Cα–Cα below 4.5 Å, C–N below 2.0 Å, no atom pair closer than 1.0 Å) |
| Side-chain rebuild of BioEmu-1 samples | HPacker 0.1.1 (commit 6cd5ec3 of github.com/gvisani/hpacker), run as in the side-chain reconstruction of bioemu 1.2.0, without relaxation |
| Scoring | be_scoring 0.1.0; Python 3.12.16; MDTraj 1.11.2; NumPy 2.3.5; SciPy 1.18.1; pandas 3.1.0; PyYAML 6.0.3 |
| Benchmark evaluator | bioemu-benchmarks 0.0.2.post1, the release whose numbers match the published BioEmu v1.1 results, in its own environment |
| Benchmark test cases | bioemu-benchmarks multi-conformation benchmark version 0.1, commit 63edbfa |
| All-atom preparation | AmberTools 22.3 (pdb4amber and tleap); ff19SB; OPC water |
| All-atom production | pmemd.cuda (Amber 20) |
| Hardware | NVIDIA A100-SXM4-80GB and V100 (BioEmu2); Tesla P100-PCIE-12GB (BioEmu-1, ColabFold and all-atom MD); NVIDIA A100-SXM4-80GB for the timing of both BioEmu models |
